# Retinoic acid breakdown is required for proximodistal positional identity during amphibian limb regeneration

**DOI:** 10.1101/2024.08.07.607055

**Authors:** Timothy J. Duerr, Melissa Miller, Sage Kumar, Dareen Bakr, Jackson R. Griffiths, Aditya K. Gautham, Danielle Douglas, S. Randal Voss, James R. Monaghan

**Author notes:** **Correspondence** James Monaghan, Northeastern University, 360 Huntington Avenue Boston, MA 02115.

## Abstract

Regenerating limbs retain their proximodistal (PD) positional identity following amputation. This positional identity is genetically encoded by PD patterning genes that instruct blastema cells to regenerate the appropriate PD limb segment. Retinoic acid (RA) is known to specify proximal limb identity, but how RA signaling levels are established in the blastema is unknown. Here, we show that RA breakdown via CYP26B1 is essential for determining RA signaling levels within blastemas. CYP26B1 inhibition molecularly reprograms distal blastemas into a more proximal identity, phenocopying the effects of administering excess RA. We identify *Shox* as an RA-responsive gene that is differentially expressed between proximally and distally amputated limbs. Ablation of *Shox* results in shortened limbs with proximal skeletal elements that fail to initiate endochondral ossification. These results suggest that PD positional identity is determined by RA degradation and RA-responsive genes that regulate PD skeletal element formation during limb regeneration.

## Introduction

Tissue regeneration requires a complex cellular choreography that results in restoration of missing structures. Salamander limb regeneration is no exception, where mesenchymal cells, including dermal fibroblasts and periskeletal cells, dedifferentiate into a more embryonic-like state and migrate to the tip of the amputated limb to form a blastema (Currie et al., 2016; Gerber et al., 2018; Lin et al., 2021). Mesenchymal cells within the blastema contain positional information which coordinates proximodistal (PD) pattern reestablishment in the regenerating limb, enabling autopod-forming blastema cells to distinguish themselves from stylopod-forming blastema cells (Kragl et al., 2009; Nacu et al., 2013; Vieira and McCusker, 2019). It has been proposed that continuous values of positional information exist along the PD axis and that thresholds of these values specify limb segments (Pescitelli and Stocum, 1981; Wolpert, 1969). These segments are genetically established by combinations of homeobox genes including Hox and *Meis* genes (Gardiner et al., 1995; Roensch et al., 2013; Takeuchi et al., 2022), and each limb segment contains a unique epigenetic profile around these homeobox genes (Kawaguchi et al., 2024). However, a mechanistic explanation for how continuous values of positional information are established and differentially interpreted by limb segments during limb regeneration is lacking.

Retinoic acid (RA) is a small, pleiotropic molecule that is pervasively involved during vertebrate morphogenesis, including initiating and patterning the developing limb. The prevailing model suggests that RA is synthesized in the lateral plate mesoderm during amniote limb development and diffuses into the limb bud to specify proximal limb identity through activation of *Meis* genes (Cooper et al., 2011; Delgado et al., 2020; Mercader et al., 2000; Niederreither et al., 2002; Roselló-Díez et al., 2011). An intrinsically timed, antagonizing gradient of fibroblast growth factors (FGFs) emanating from the apical ectodermal ridge (AER) then creates a zone of distal identity marked by *Hoxa13* expression (Mercader et al., 2000; Probst et al., 2011; Saiz-Lopez et al., 2015). This activates CYP26B1, which eliminates RA from the distal limb cells and creates a gradient of RA that patterns the developing limb PD axis (Yashiro et al., 2004). Perturbing the establishment of this gradient often results in PD patterning defects (Niederreither et al., 1999; Yashiro et al., 2004). Our understanding of the role of RA in urodele limb development is not as comprehensive as in amniotes, but some aspects have been elucidated. Similar to amniotes, a gradient of active RA signaling exists along the developing urodele PD axis (Monaghan and Maden, 2012) and disruption results in abnormal skeletal morphologies along the PD axis (Maden, 1998; Nguyen et al., 2017; Scadding and Maden, 1986).

As in limb development, RA concentration is thought to differentiate the positional identity of upper limb blastemas from lower limb blastemas, thereby ensuring regeneration of the appropriate PD limb structures from disparate amputation planes. RA synthesis occurs endogenously within the blastema in response to injury (Scadding and Maden, 1994; Viviano et al., 1995), and RA signaling is approximately 3.5 times higher in proximal blastemas (PBs) compared to distal blastemas (DBs) (Brockes, 1992). These endogenous levels of RA are necessary for proper regeneration (Lee et al., 2012; Maden, 1998; Scadding, 1999). Furthermore, administering RA to DBs reprograms blastema cells to a proximal identity, resulting in regeneration of more proximal structures in a concentration dependent manner (Maden, 1982; Thoms and Stocum, 1984). This reprogramming is associated with downregulation of distal limb patterning genes like *Hoxa13* and upregulation of proximal limb patterning genes like *Meis1* and *Meis2* (Gardiner et al., 1995; Nguyen et al., 2017; Polvadore and Maden, 2021). In agreement with this, perturbing *Meis1* and *Meis2* in RA treated DBs partially blocks limb duplication (Mercader et al., 2005). These studies collectively point to a model whereby heightened RA signaling in PBs specifies proximal positional identity through activation of proximal patterning genes and repression of distal patterning genes.

Despite strong evidence that RA regulates positional identity along the regenerating PD axis, the differences in RA signaling levels between PBs and DBs are not fully understood. To address this, we examined the spatiotemporal expression of limb patterning genes in PBs and DBs and compared it to expression of genes related to RA synthesis, degradation, and signaling. We found that *Cyp26b1* was more highly expressed in mesenchymal cells of DBs than PBs, suggesting that differences in RA signaling levels between PBs and DBs are due to RA degradation. Pharmacological inhibition of CYP26B1 in DBs increased RA signaling and resulted in concentration-dependent duplications of proximal limb segments. These duplications occurred by repressing distal limb patterning genes and activating proximal limb patterning genes. Two such genes, *Shox* and *Shox2*, are both RA-responsive and differentially expressed in PBs and DBs. Disruption of *Shox* yields phenotypically normal autopods but shortened stylopods and zeugopods that fail to initiate endochondral ossification. Moreover, we show that *Shox* is not required for limb regeneration but is crucial for endochondral ossification of stylopodial and zeugopodial skeletal elements during regeneration. Our results collectively suggest that RA breakdown via CYP26B1 is required for establishing positional identity along the regenerating PD axis, enabling the activation of genes such as *Shox* that confer proximal limb positional identity.

## Methods

### Animal procedures & drug treatments

Transgenic lines (RA reporter-tgSceI(RARE:GFP)^Pmx^, *Hoxa13* reporter-tm(*Hoxa13*^t/+^:*Hoxa13*-T2A-mCHERRY)^Etnka^) used in this study were bred at Northeastern University. d/d genotype axolotls (RRID:AGSC 101E; RRID:AGSC101L) were bred at Northeastern University or were obtained from the Ambystoma Genetic Stock Center (RRID: SCR_006372). Crispant lines were generated at both Northeastern University and the University of Kentucky. All surgeries were performed under anesthesia using 0.01% benzocaine and were approved by the Northeastern University Institutional Animal Care and Use Committee.

Talarozole and AGN 193109 were resuspended in dimethyl sulfoxide (DMSO) to 5 mM and stored at -20°C. Disulfiram was resuspended in DMSO to 20 mM and stored at - 20°C. Animals received amputations through the carpals on one limb and through the upper humerus on the contralateral limb. Four days post amputation (DPA), animals were placed in drug or DMSO. Animals were redosed every other day for 7 days. Following treatment, animals were removed from the drug water and placed into axolotl housing water. Regenerated limbs were collected from the animals 120 days post treatment for analysis of skeletal morphology.

### qRT-PCR

Blastema tissue (4-5 blastemas per sample, 3-6 samples per experiment) was collected from 3.5 cm animals head to tail (HT) aged 2.5 months and immediately frozen in liquid nitrogen. RNA isolation and qRT-PCR were performed with standard protocols, using 1 ng of cDNA per reaction. Each qRT-PCR reaction was run in duplicate. Primers for each gene are listed in Table S1. Gene expression was normalized to *Ef1α* and the Livak 2^−ΔΔCq^ method (Livak and Schmittgen, 2001) was used to quantify relative fold change in mRNA abundance. Statistical significance between groups was tested using either a two-tailed Student’s t-test or a one-way analysis of variance (ANOVA) with post-hoc Tukey’s honestly significant difference (HSD) test on ΔCt values. Linear regression analysis was performed to test for linear relationships across samples.

### Single-cell gene expression quantification and reanalysis

Existing single-cell RNA sequencing (scRNA-seq) data was accessed from NCBI SRA PRJNA589484 (Li et al., 2021). UMI counts for gene expression data were generated using kallisto (Bray et al., 2016) and bustools (Melsted et al., 2019) against *Ambystoma mexicanum* transcriptome v4.7 (Schloissnig et al., 2021). Isoform-level count matrices were generated using “bustools count” with the --em flag. Counts matrices were imported into Seurat v5 (Hao et al., 2024) in an RStudio IDE. Cells expressing fewer than 200 features and features expressed in fewer than 3 cells were filtered out; matrices were further filtered to include cells with between 500 and 25000 counts, <5% red blood cell gene content, and <15% mitochondrial gene content. Counts were normalized using the SCTransform function (Choudhary and Satija, 2022) with regression for mitochondrial content and red blood cell content. The counts layers were flattened with the IntegrateLayers function using RPCAIntegration, after which FindClusters was run with a resolution of 0.4 followed by RunUMAP using the first 30 dimensions, 50 nearest neighbors, and a minimum distance of 0.1. Following cluster annotation, SCTransform-normalized isoform-level expression for genes of interest was summed and log-transformed for plotting with ggplot2 using UMAP coordinates from Seurat (Wickham, 2016).

### HCR-FISH and imaging

Whole mount and tissue section version 3 hybridization chain reaction fluorescence *in-situ* hybridization (HCR-FISH) protocols were performed as previously described (Lovely et al., 2023) with slight modifications on the tissue section protocol. Namely, fresh frozen blastema tissue 3.5 cm animals (HT) aged 2.5 months were sectioned at 10 µm. Slides were then fixed with 4% paraformaldehyde (PFA) for 15 minutes at room temperature and washed with 1X phosphate buffered saline (PBS) three times for five minutes at room temperature before being stored in 70% ethanol at 4°C until use. Slides were washed again three times for five minutes with 1X PBS at room temperature before beginning HCR-FISH protocol. Probes for each gene were generated using ProbeGenerator (http://ec2-44-211-232-78.compute-1.amazonaws.com) and can be found in Table S2.

All tissue sections were imaged using a Zeiss LSM 880 confocal microscope at 20X magnification with airyscan fast settings. For each image, 4-6 optical sections were captured in the 10 µm section. Following acquisition, images were processed on Zen Black using airyscan processing with the automatic 2D setting, then a maximum intensity projection step was performed. If obvious issues in tile alignment were observed, an additional stitching step was performed prior to maximum intensity projection.

Whole mount samples stained using HCR-FISH were mounted in 1.5% low-melt agarose and refractive index matched with EasyIndex (LifeCanvas Technologies) overnight at 4°C (Lovely et al., 2023). Samples were then imaged in EasyIndex using a Zeiss Lightsheet Z1 at 20X magnification. All representative images are a single z-plane from the stack.

### HCR-FISH dot visualization and quantification

Dot detection in HCR-FISH images was performed using FIJI plugin RS FISH (Bahry et al., 2022; Schindelin et al., 2012). For visualization of HCR-FISH dots in figure pictures, RS-FISH dots were flattened atop the DAPI channel and background dots outside tissue sections were removed. For quantification, identified dots from RS-FISH were overlaid on a black background and the image was flattened. The ImageJ function “Find Maxima” with prominence greater than 0 was used to convert dots to single pixels. Cell outlines were obtained using Cellpose segmentation run on the DAPI channel using the “cyto2” model with default settings and a diameter of 60 (Pachitariu and Stringer, 2022; Stringer et al., 2021). A modified version of the region of interest conversion script provided by the Cellpose authors was used to obtain measurements of area and raw integrated density per cell. Pipeline documentation can be accessed at https://github.com/Monaghan-Lab/HCRFISH-DotCounting.

All measurements were concatenated and filtered for cells with an area greater than 60, as smaller areas can indicate poor segmentation or cells at a nonoptimal plane. To evaluate the expression levels for each gene within a sample, an unpaired, two-sided clustered Wilcoxon rank-sum test (n = 3-6 blastemas) was performed with FDR p-adjustment using the R package “clusrank” with test method “ds” (Datta and Satten, 2005; Jiang et al., 2020). Violin plots were generated in RStudio using the ggplot2 package (Wickham, 2016).

### HCR-FISH PD intensity measurements

Images of RS-FISH maxima were rotated in FIJI with no interpolation such that the top of the image represented the most proximal end of the blastema. An equal rotation was performed on the DAPI channel from the same tissue. The freehand selection tool was used to isolate the blastemal mesenchyme from the DAPI image, then this selection was overlaid onto the rotated maxima image and used to clear any points lying outside of the selection before adding a value of 25 to each pixel within the blastema selection. The value z at every (x,y) pixel in the image was then measured.

Measurements for each image were imported into R and filtered for z > 0 to select for points falling only within the blastemal mesenchyme. The y values were then rescaled to generate a pseudo-proximodistal axis with range [0, 1]. Z values were reassigned such that 25 represented a 0 or "negative" pixel and 255 represented a 1 or "positive" pixel, then grouped at each pseudo-y value and averaged to yield a proportion of positive pixels. The square root of these proportions was plotted as a smoothed curve along the pseudo-proximodistal axis using the "stat_smooth()" function in ggplot2 with an *n* of 2000 (Wickham, 2016). A sample script can be accessed at https://github.com/Monaghan-Lab/HCRFISH-DotCounting.

### Alcian blue/alizarin red staining

Whole mount alcian blue/alizarin red staining was performed using a modified version of a previously published protocol (Riquelme-Guzmán and Sandoval-Guzmán, 2023). Briefly, regenerated limbs were collected and immediately fixed in 4% PFA overnight at 4°C. The next day, limbs were washed three times for five minutes each with 1X PBS at room temperature and skinned. Limbs were dehydrated in 25%, 50%, and 100% ethanol for 20 minutes at each concentration before being placed in alcian blue mixture (5 mg alcian blue in 30 mL 100% ethanol, 20 mL acetic acid) and left on a rocker overnight at room temperature. The next day, limbs were rehydrated in 100%, 50%, and 25% ethanol for 20 minutes at room temperature at each concentration. Limbs were placed into trypsin solution (1% trypsin in 30% borax) and rocked for 45 minutes at room temperature. Samples were washed twice in 1% KOH for 30 minutes each before being placed in alizarin red mixture (5 mg alizarin red in 50 mL 1% KOH) and rocked overnight at room temperature. Limbs were again washed twice with 1% KOH for 30 minutes at room temperature and placed in 25% glycerol/1% KOH solution until samples cleared. The limbs were then dehydrated in 25%, 50%, and 100% ethanol for 20 minutes at room temperature at each concentration. Finally, limbs were placed in 25%, 50%, and 75% glycerol solutions (made in 100% ethanol) before being stored in 100% glycerol for imaging.

### Bulk RNA sequencing and analysis

Animals received a proximal amputation through the upper humerus of the left forelimb and a distal amputation through the carpals of the right forelimb. TAL was administered in the housing water as indicated above. Samples (n = 3 samples per condition, 2-3 blastemas per sample, 5 cm (HT) animals aged 3 months) were collected at 14 DPA and immediately flash frozen in liquid nitrogen before storing at -80°C. Samples were shipped to Genewiz for RNA sequencing using an Illumina HiSeq platform and 150-bp paired-end reads for an average sequencing depth of roughly 24.1 million reads per sample. Raw sequencing data are available at GEO (accession number GSE272731).

Reads were quality trimmed with Trimmomatic (Bolger et al., 2014) before quasi-mapping to the v.47 axolotl transcriptome (Nowoshilow et al., 2018) and quantification with salmon v0.13.1 (Patro et al., 2017). Differential expression analysis was performed on counts matrices with DESeq2 v1.34.0 (Love et al., 2014) using the Trinity v2.8.5 script (Grabherr et al., 2011) “run_DE_analysis.pl” with default parameters. Visualizations were produced with ggplot2, ggvenn, and ComplexHeatmap (Gu et al., 2016) where appropriate. Sample correlation heatmap was produced using Trinity script “PtR” on the gene counts matrix (Grabherr et al., 2011).

### Generating and genotyping *Shox* crispant axolotls

*Shox* crispants were generated using CRISPR/Cas9 according to previous protocols (Fei et al., 2018; Trofka et al., 2021). The following sgRNAs were used:

*Shox* sgRNA 2- GAGGGAGGACGTGAAGTCGG

*Shox* sgRNA 3- GGCCAGGGCCCGGGAGCTGG

NGS-based genotyping was conducted on a pool of 10 tail tips from crispant animals and analyzed using CRISPResso2 (Clement et al., 2019).

### Hematoxylin, eosin, and alcian blue staining

Samples for hematoxylin, eosin, and alcian blue staining (H&E&A) were collected and placed in 4% PFA overnight at 4°C. Samples were then washed with 1X PBS three times for five minutes each. Following the third wash, samples were placed in 1 mM ethylenediaminetetraacetic acid (EDTA) for four days at 4°C. The EDTA solution was changed every other day. After EDTA treatment, the samples were again washed with 1X PBS three times for five minutes each before being cryoprotected in 30% sucrose. Once the samples sunk in the 30% sucrose, the samples were mounted in optimal cutting temperature medium and frozen at -80°C until use. These blocks were then sectioned at 10 µm, and the resultant slides were baked at 65°C overnight. To improve the adherence of the skeletal structure to the slide, slides were placed in 4% PFA for 15 minutes at room temperature. Slides were washed three times for five minutes with 1X PBS, then the slides were placed in alcian blue solution (5 mg of alcian blue in 30 mL 100% ethanol and 20 mL acetic acid) for ten minutes at room temperature. The slides were then dehydrated with 100% EtOH for one minute at room temperature and allowed to air dry. Hematoxylin solution was added to the slides and incubated at room temperature for seven minutes. Slides were then dipped into tap water five times, then clean tap water another 15 times. Slides were dipped another 15 times in clean tap water before adding bluing buffer for two minutes at room temperature. Slides were again dipped in clean tap water five times, and eosin solution was pipetted onto the slides for two minutes at room temperature. Residual eosin was removed by dipping slides ten times in clean tap water, and the slides were air dried before imaging.

## Results

### The spatiotemporal expression of PD patterning genes differs in PBs and DBs

To explore how patterning genes regulate RA signaling in PBs and DBs, we examined the expression of known RA-responsive homeobox genes, including *Meis1* and *Meis2* (Mercader et al., 2000; Mercader et al., 2005). We assessed the expression of these genes in limbs amputated at the upper stylopod (US), lower stylopod (LS), upper zeugopod (UZ), and autopod levels at 10 DPA using qRT-PCR and found that *Meis1* and *Meis2* expression decreases in progressively distal amputations (Fig. 1A-C). We next reanalyzed scRNA-seq data from DBs to identify the cell types that express *Meis1* and *Meis2* (Fig. S1) (Li et al., 2021). We observed *Meis1* expression in mesenchymal and epithelial cells while *Meis2* was undetected (Fig. S2A-B).

**Figure 1:**
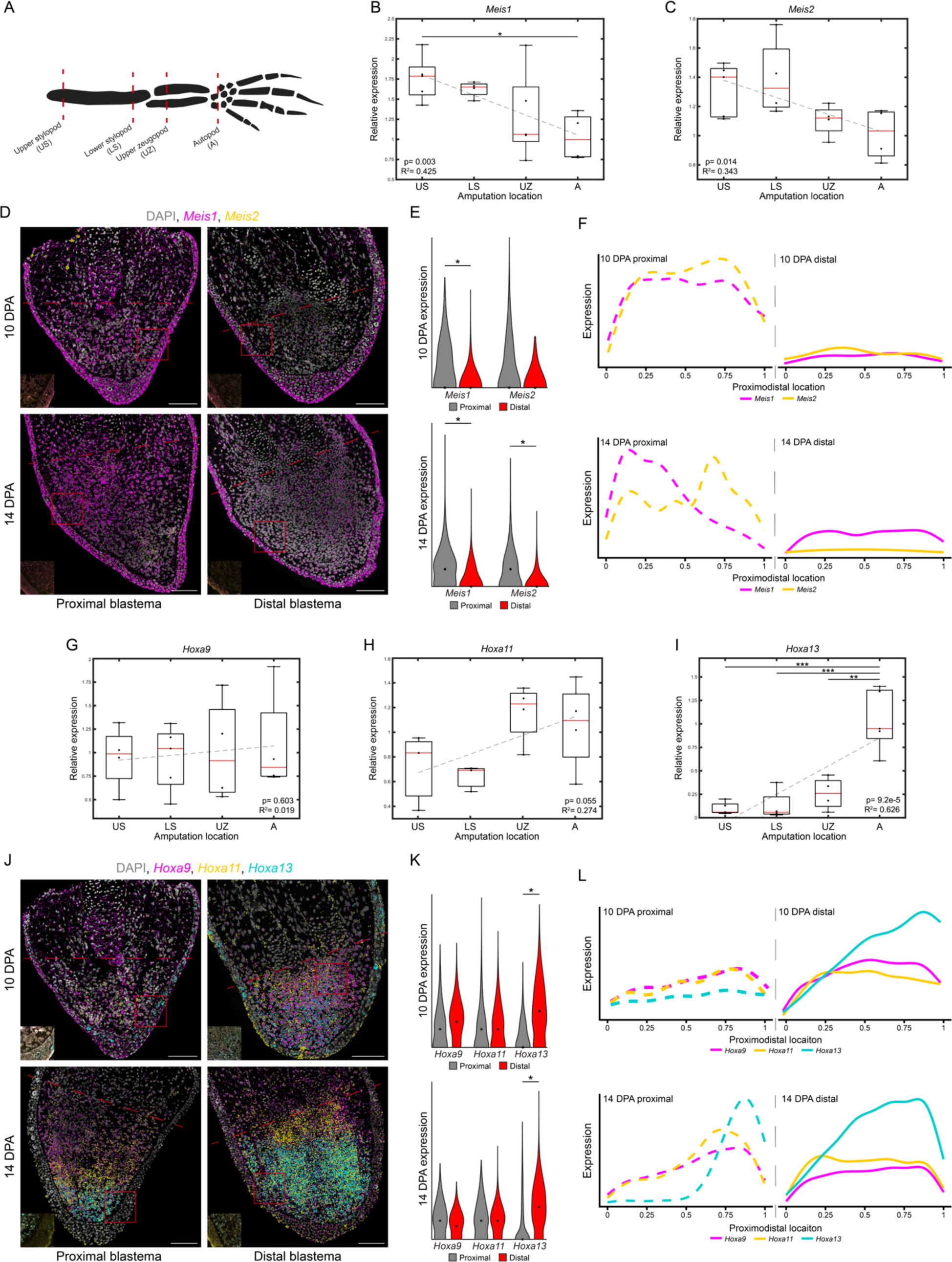
PD patterning genes are dynamically expressed during limb regeneration. (A) Schematic of PD amputation plane qRT-PCR experiment. (B-C) qRT-PCR of *Meis1* (B) and *Meis2* (C) at different PD amputation locations (n = 3-6, 4-5 blastemas per sample, 3.5 cm (HT) animals aged 2.5 months, 10 DPA). Each gene was normalized to *Ef1a* and analyzed with a one-way ANOVA using a Tukey-Kramer multiple comparison test. R^2^ and p values from linear regression analysis are shown. * = p < 0.05. (D) HCR-FISH for *Meis1* and *Meis2* in PBs and DBs at 10 and 14 DPA. Dashed lines indicate amputation plane. Scale bars = 200 µm or 20 µm (inset). (E) HCR-FISH dot quantification for mesenchymal *Meis1* and *Meis2* in PBs and DBs at 10 and 14 DPA (n = 3-6, 3.5 cm (HT) animals aged 2.5 months). Expression is the square root of RS-FISH dots within ROIs. Groups were analyzed using a clustered Wilcoxon rank sum test according to the Datta-Satten method. * = p < 0.05. (F) PD intensity plots for mesenchymal *Meis1* and *Meis2* in PBs and DBs at 10 and 14 DPA (n = 3-6, 3.5 cm (HT) animals aged 2.5 months). Lines represent average signal intensity (expression) along a normalized PD axis across each sample. (G-I) qRT-PCR of *Hoxa9* (G), *Hoxa11* (H), and *Hoxa13* (I) at different PD amputation locations (n = 3-6, 4-5 blastemas per sample, 3.5 cm (HT) animals aged 2.5 months, 10 DPA). Analyses as in Fig. 1B-C. ** = p < 0.01, *** = p < 0.001. (J) HCR-FISH for *Hoxa9*, *Hoxa11*, and *Hoxa13* in PBs and DBs at 10 and 14 DPA. Dashed lines indicate amputation plane. Scale bars = 200 µm or 20 µm (inset). (K) HCR-FISH dot quantification for mesenchymal *Hoxa9*, *Hoxa11*, and *Hoxa13* in PBs and DBs at 10 and 14 DPA (n = 3-6, 3.5 cm (HT) animals aged 2.5 months). Axes and analyses as in Fig. 1E. * = p < 0.05. (L) PD intensity plots for mesenchymal *Hoxa9*, *Hoxa11*, and *Hoxa13* in PBs and DBs at 10 and 14 DPA (n = 3-6, 3.5 cm (HT) animals aged 2.5 months). Axes and analyses as in Fig. 1F.

We then visualized *Meis1* and *Meis2* in DBs at 7, 10, and 14 DPA and found *Meis1* was expressed only in the epithelium while *Meis2* was nonexistent (Fig. 1D-E, S2C-D). Similarly, PBs at 7 DPA exhibited low mesenchymal *Meis1* and *Meis2* expression (Fig. S2C-D). At 10 DPA, *Meis1* was significantly higher in the mesenchyme of PBs, and *Meis2* was elevated in PBs but was generally lowly expressed (Fig. 1D-E). By 14 DPA, *Meis1* was significantly higher in PBs and localized to the proximal-most mesenchyme cells, creating a distal zone devoid of *Meis1* expression (Fig. 1D-F). A similar *Meis1* expression pattern was observed during axolotl (Fig. S2E), mouse, and chick limb development (Mercader et al., 2009; Roselló-Díez et al., 2014), suggesting an evolutionarily conserved role reutilized for limb regeneration. *Meis2* was significantly higher in PBs at 14 DPA but was generally lower than *Meis1* (Fig. 1D-E). Notably, *Meis2* appeared more abundant in the proximal portion of the developing limb bud (Fig. S2E). Epithelial expression of *Meis1* and *Meis2* did not differ between PBs and DBs at any time point or amputation location (Fig. S2D), underscoring the importance of mesenchymal cells in conveying PD positional identity within the blastema.

We next examined *Hoxa9*, *Hoxa11*, and *Hoxa13* which have a known role in establishing both the amphibian regenerating and developing PD axis by providing positional identity to each limb segment (Gardiner et al., 1995; Roensch et al., 2013; Takeuchi et al., 2022). We observed similar *Hoxa9* expression in blastemas at each amputation location (Fig. 1G), while *Hoxa11* was elevated in blastemas amputated at the UZ and autopod levels relative to US and LS amputations (Fig. 1H). *Hoxa13* was significantly more highly expressed in autopod amputations compared to any other amputation level (Fig. 1I), reflecting activation of more 5’ Hox genes in increasingly distal limb amputations. Additionally, we found that *Hoxa9*, *Hoxa11*, and *Hoxa13* were predominately expressed in mesenchymal cells (Fig. S2F-H).

We observed that *Hoxa9*, *Hoxa11*, and *Hoxa13* were expressed uniformly in the mesenchyme of DBs at 7, 10, and 14 DPA (Fig. 1J, Fig. S2I). In PBs at 7 DPA, only *Hoxa9* was expressed in the mesenchyme while *Hoxa11* and *Hoxa13* were absent (Fig. S2I). By 10 DPA, *Hoxa9* and *Hoxa11* were expressed in the mesenchyme of PBs, but *Hoxa13* remained low (Fig. 1J-K, Fig. S2J). At 14 DPA, *Hoxa9* was expressed throughout the mesenchyme of PBs at similar levels as DBs (Fig. 1J-K, Fig. S2J), and *Hoxa11* was detected in cells from the mid-blastema to the distal tip (Fig. 1J-L). *Hoxa13* was detected in the distal-most mesenchymal cells of PBs, although at lower levels than in DBs (Fig. 1J-L). This colinear activation of 5’ Hox genes mirrors *Hoxa9*, *Hoxa11*, and *Hoxa13* expression during limb development (Fig. S2K), supporting the hypothesis that progressive specification establishes PD positional identity in both processes (Roensch et al., 2013).

### RA signaling levels within blastemas are determined by *Cyp26b1* expression

We hypothesized that RA signaling pathway members regulate RA signaling levels in response to PD patterning genes. We reasoned that a candidate gene would be expressed in the blastema mesenchyme, show graded expression along the PD axis, and complement the spatiotemporal expression of *Meis1*, *Hoxa11*, and *Hoxa13*, which direct limb morphogenesis (Uzkudun et al., 2015). To this end, we focused on RA receptors (*Rara*, *Rarg*) and genes involved in RA synthesis (*Raldh1*, *Raldh2*, *Raldh3*) and RA degradation (*Cyp26a1*, *Cyp26b1*). Both *Rara* and *Rarg* were expressed in the mesenchyme, but their expression did not match *Meis1*, *Hoxa11*, and *Hoxa13* (Fig. S5). *Raldh1* and *Raldh3* were primarily expressed in the epithelium, and *Raldh2* expression did not differ within or between PBs and DBs (Fig. S6). For these reasons, we ruled out RARs and RALDHs as modulators of RA signaling in the blastema. Next, we investigated if RA signaling levels are controlled by degradation in PBs and DBs. We found that *Cyp26a1* was higher in US blastemas and decreased in distal amputations (Fig. 2A). Conversely, *Cyp26b1* expression was highest in autopod level amputations and decreased in more proximal amputation locations (Fig. 2B). *Cyp26a1* and *Cyp26b1* expression levels appeared to form gradients independent of limb segment, with similar levels in blastemas amputated at the LS and UZ (Fig. 2A-B). ScRNA-seq did not detect *Cyp26a1*, but *Cyp26b1* was highly expressed in the mesenchymal and epidermal of DBs (Fig. S7A-B). *Cyp26a1* was similar in PBs and DBs and showed no bias towards the epithelium or mesenchyme (Fig. 2C-D, Fig. S7C). This difference between our HCR-FISH and qRT-PCR results is likely due to the higher sensitivity of qRT-PCR, although both assays indicate that *Cyp26a1* is lowly expressed and did not match expression of *Hoxa13*, *Hoxa11*, or *Meis1* (Fig. S7E). In contrast, epithelial *Cyp26b1* was observed in PBs and DBs at 7 DPA but was more highly expressed in the mesenchyme of DBs (Fig. S7C). By 10 DPA, *Cyp26b1* was similar in the epithelium of PBs and DBs, but significantly higher in the mesenchyme of DBs (Fig. 2C-D). At 14 DPA, *Cyp26b1* was no longer differentially expressed between PBs and DBs (Fig. 2C-D), but was concentrated at the distal tip of PBs, tapering off proximally (Fig. 2E). Mesenchymal *Cyp26b1* expression at 7, 10, and 14 DPA closely mirrors *Hoxa11* and *Hoxa13* and appears anticorrelated with *Meis1* (Fig. 2E). These results indicate that *Cyp26b1* is graded in mesenchymal cells along the PD axis and associated PD patterning genes.

**Figure 2:**
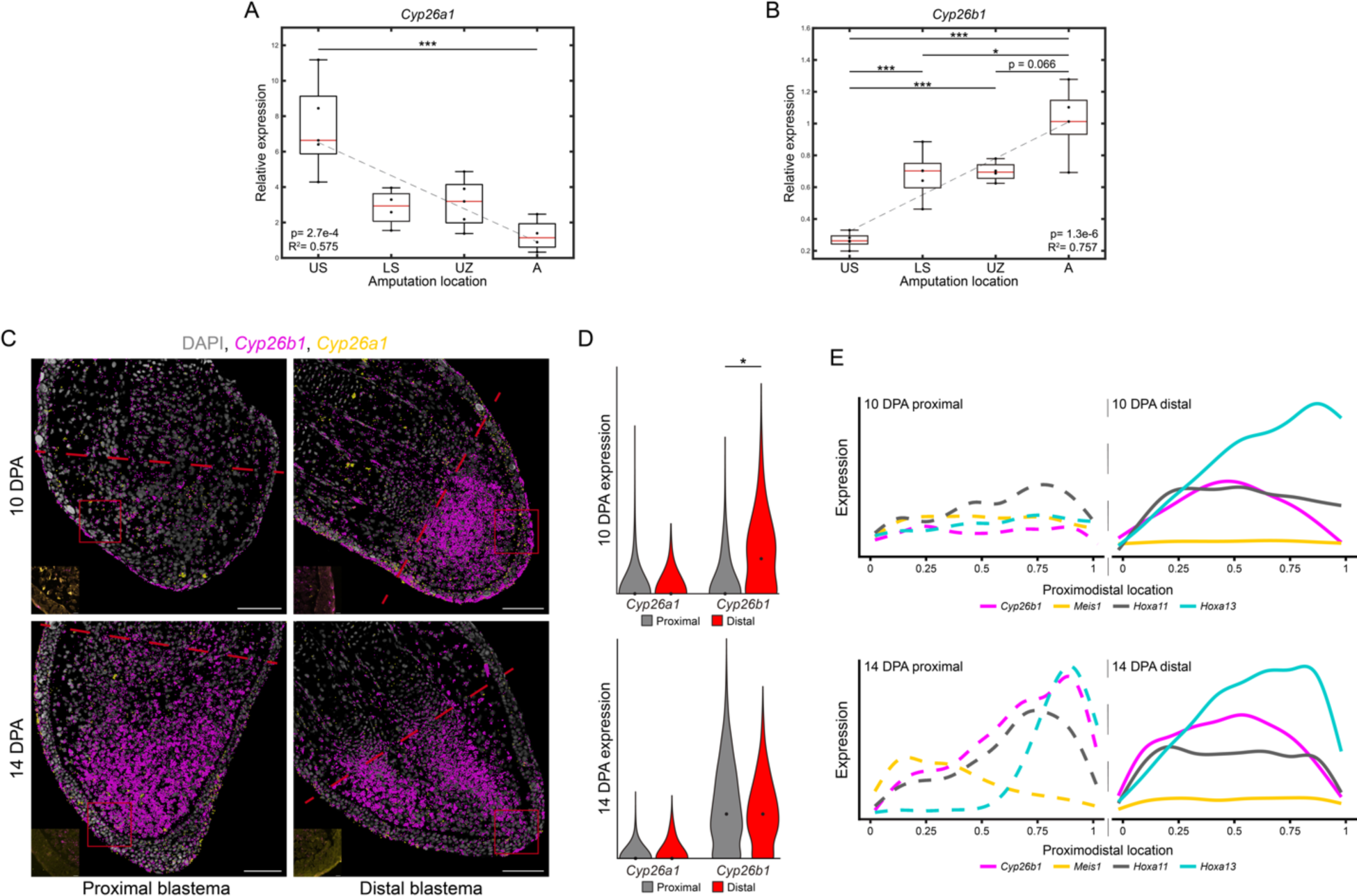
*Cyp26b1* is differentially expressed in PBs and DBs and correlates with *Meis1, Hoxa11*, and *Hoxa13* expression. (A-B) qRT-PCR of *Cyp26a1* (A) and *Cyp26b1* (B) at different PD amputation locations (n = 3-6, 4-5 blastemas per sample, 3.5 cm (HT) animals aged 2.5 months, 10 DPA). Analyses as in Fig. 1B-C. * = p < 0.05, *** = p < 0.001. (C) HCR-FISH for *Cyp26a1* and *Cyp26b1* in PBs and DBs at 10 and 14 DPA. Dashed lines indicate amputation plane. Scale bars = 200 µm or 20 µm (inset). (D) HCR-FISH dot quantification for mesenchymal *Cyp26a1* and *Cyp26b1* in PBs and DBs at 10 and 14 DPA (n = 3-6, 3.5 cm (HT) animals aged 2.5 months). Axes and analyses as in Fig. 1E. * = p < 0.05. (E) PD intensity plots for mesenchymal *Cyp26b1*, *Meis1*, *Hoxa11*, and *Hoxa13* in PBs and DBs at 10 and 14 DPA. Axes and analyses as in Fig. 1F.

### CYP26 inhibition increases RA signaling and duplicates proximal skeletal structures

We hypothesized that positional identity along the regenerating PD axis depends on RA degradation in the blastema. To test this, we used talarozole (TAL, or R115866) to inhibit CYP26 during limb regeneration. Three TAL concentrations (0.1 µM, 1 µM, and 5 µM) or DMSO were administered at 4 DPA (Thoms and Stocum, 1984) to animals with proximally amputated left limbs and distally amputated right limbs for 7 days (Fig. 3A). The skeletal structures of regenerates were then analyzed for abnormalities in morphology (Fig. 3A). At 14 DPA, we observed that drug treated blastemas were smaller than DMSO controls (Fig. S8A-B). After fully regenerating, DMSO treated limbs and proximally amputated limbs treated with 0.1 µM or 1 µM TAL exhibited normal skeletal morphology (Fig. 3B, Table S3). At 5 µM TAL, 15.4% of proximally amputated limbs regenerated without skeletal irregularities while limbs in the remaining 84.6% regressed to the scapula and failed to regenerate (Fig. 3B). Conversely, 92.3% of distally amputated limbs treated with 0.1 µM TAL exhibited whole (61.2%) or partial (30.1%) zeugopod duplications while 7.7% displayed no limb duplications (Fig. 3B). Increasing TAL to 1 µM resulted in full (66.7%) or partial (33.3%) stylopod duplications from distal amputations (Fig. 3B). Treatment with 5 µM TAL inhibited regeneration in 92.3% of distally amputated limbs, while 7.7% showed full humerus duplication. (Fig. 3B). These results mirror both the inhibition of limb regeneration by excess retinoids (Maden, 1983) and PD duplications observed in regenerating *Xenopus laevis* hindlimbs after TAL treatment (Cuervo and Chimal-Monroy, 2013).

**Figure 3:**
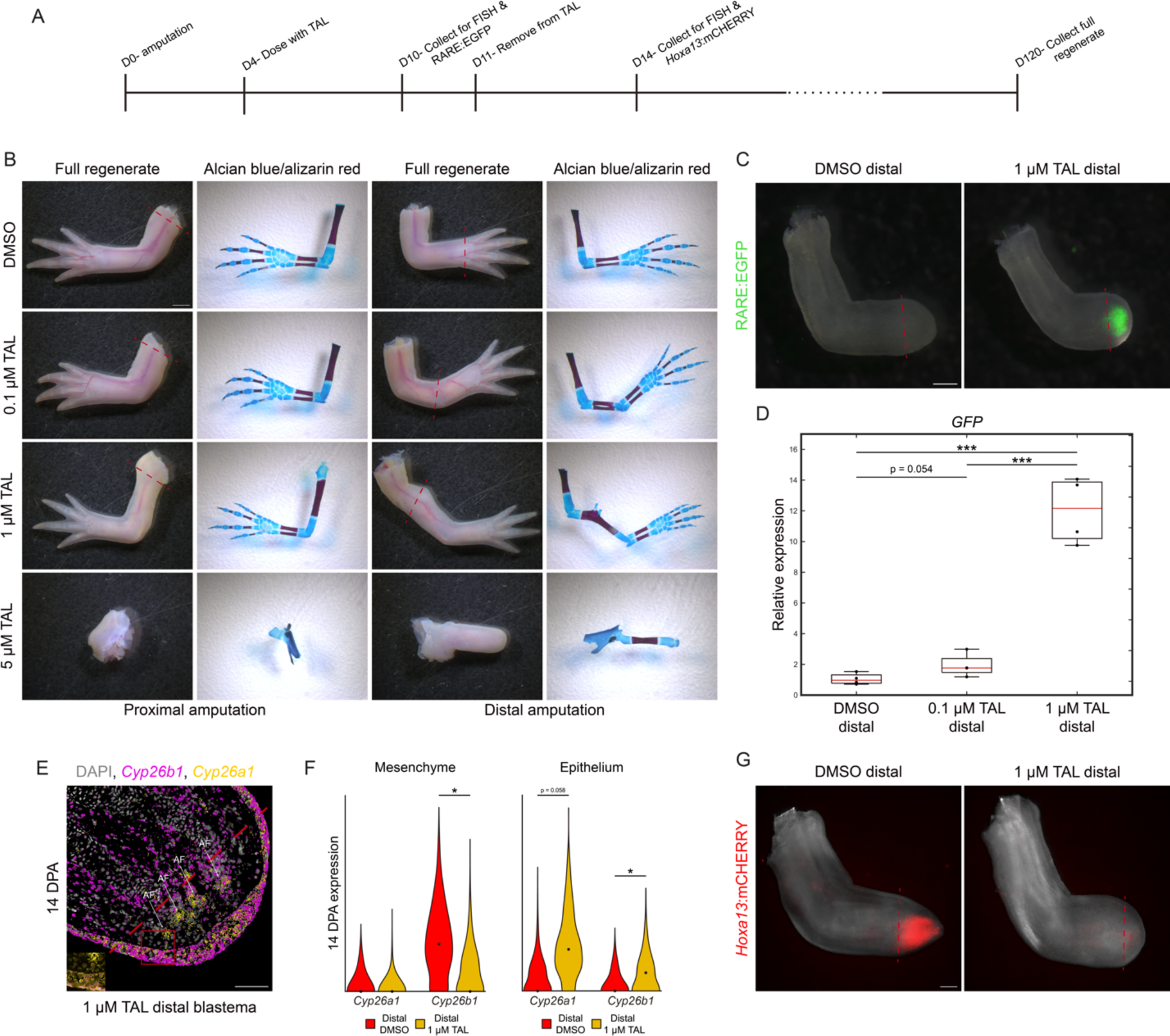
CYP26 inhibition phenocopies exogenous RA during limb regeneration. (A) Timeline of TAL experiments and tissue collection timepoints. (B) Brightfield images of regenerates and skeletal structures of PBs and DBs treated with DMSO or 0.1, 1, or 5 µM TAL. Dashed lines indicate amputation plane. Scale bar = 2 mm. (C) 10 DPA DBs from RA reporter animals treated with DMSO or 1 µM TAL (n = 8, 3 cm (HT) animals aged 2 months). Dashed lines indicate amputation plane. Scale bar = 500 µm. (D) qRT-PCR of *Gfp* in tissue from RA reporter animals. (n = 4, 4 blastemas per sample, 3.5 cm (HT) animals aged 2.5 months, blastemas collected at 10 DPA). Analyses as in Fig. 1B-C. *** = p < 0.001. (E) HCR-FISH for *Cyp26a1* and *Cyp26b1* in DBs administered 1 µM TAL at 14 DPA (n = 3-6, 3.5 cm (HT) animals aged 2.5 months). Dashed line indicates amputation plane. AF = autofluorescence. Scale bar = 200 µm or 20 µm (inset). (F) HCR-FISH dot quantification for mesenchymal and epithelial *Cyp26a1* and *Cyp26b1* in DBs treated with DMSO or 1 µM TAL at 14 DPA (n = 3-6, 3.5 cm (HT) animals aged 2.5 months). Axes and analyses as in Fig. 1E. * = p < 0.05. (G) 14 DPA DBs from *Hoxa13*:mCHERRY reporter animals treated with DMSO or 1 µM TAL (n = 8, 7.5 cm (HT) animals aged 6 months). Dashed lines indicate amputation plane. Scale bar = 500 µm.

Limb duplications after TAL treatment resembled the effects of administering excess RA to regenerating limbs. To determine if TAL increased RA signaling, we administered 0.1 µM or 1 µM TAL to RA reporter animals (Monaghan and Maden, 2012) and assessed GFP expression in DBs at 10 DPA. We observed increased GFP signal and *Gfp* expression following TAL administration (Fig. 3C-D), indicating TAL increased RA signaling in the blastema. Additionally, mesenchymal *Cyp26b1* decreased in DBs treated with 1 µM TAL while *Cyp26a1* was unchanged (Fig. 2C, Fig. 3E-F). However, *Cyp26a1* was elevated following TAL treatment, (Fig. 3E-F), suggesting that CYP26A1 degrades detrimental levels of RA in the epithelium but does not pattern the regenerating limb.

Previous studies have shown that excess RA decreases *Hoxa13* (Gardiner et al., 1995; Nguyen et al., 2017; Roensch et al., 2013), leading us to hypothesize that *Hoxa13* would similarly decrease following TAL treatment. To address this, we utilized *Hoxa13* reporter animals (Oliveira et al., 2022) to visualize how TAL impacts HOXA13. DMSO treated DBs demonstrated strong mCHERRY at 14 DPA, whereas expression was absent in DBs treated with 1 µM TAL (Fig. 3G). We also observed reduced *Hoxa13* at the transcript level (Fig. S8C-D), showing that *Hoxa13* decreases following TAL administration.

Our results indicate that TAL increases mesenchymal RA signaling in DBs, leading to both concentration-dependent increases in RA signaling and skeletal morphologies that mimic RA-induced limb duplications. Considering *Cyp26a1* and *Cyp26b1* expression (Fig. 2C-D, Fig. 3E-F), the final skeletal morphologies are likely due to inhibition of CYP26B1, not CYP26A1. Moreover, we observed no change in skeletal morphology of proximally amputated limbs treated with the same TAL concentration that caused full limb duplication in distally amputated limbs, indicating RA breakdown is crucial for positional identity in distally, not proximally, amputated limbs. However, most PBs treated with 5 µM TAL failed to regenerate, suggesting that some RA breakdown is necessary for limb regeneration in PBs (Fig. 3B, Table S3).

### Limb duplications require RA synthesis and RAR activity

To elucidate the mechanism behind TAL-induced proximalization of DBs, we cotreated animals with TAL and either disulfiram (DIS), a pan RALDH inhibitor, or AGN 193109 (RAA), a pan-RAR antagonist. Treatments were administered as above (Fig. 3A), with DIS or RAA administered alone or with 1 µM TAL. Limb duplications were not observed in DMSO treated limbs, regardless of treatment condition (Fig. S9A-B, Table S4, Table S5). Animals treated or with DIS or RAA alone showed minor skeletal abnormalities but no duplications in skeletal morphology (Table S4, Table S5). In distally amputated limbs treated with 1 µM TAL and 0.1 µM DIS, all limbs were duplications at the radius/ulna level (Fig. S9A, Table S4). Similarly, 80% of the limbs treated with 1 µM TAL and 1 µM DIS showed either half duplication of the radius/ulna or no duplication (Table S4). These results show that DIS inhibits full proximalization of TAL-treated DBs, indicating that RA synthesis is required for proximalization. We next administered 1 µM TAL and either 0.1 µM or 1 µM RAA to animals with proximal and distal amputations. None of the distally amputated limbs treated with 1 µM TAL and 0.1 µM RAA showed duplications (Fig. S9B, Table S5), and 83.3% of those treated with 1 µM TAL and 1 µM RAA (Table S5) also lacked duplication. This shows RA signaling through RARs is necessary for limb duplication.

### CYP26 inhibition transcriptionally reprograms DBs into a more PB-like identity

We next hypothesized that increasing TAL concentration in DBs would progressively reprogram DBs into a more PB-like identity. Additionally, we hypothesized that reprogramming would be driven by RA-responsive genes that are differentially expressed along the PD axis. We tested these hypotheses using bulk RNA-seq on DBs treated with DMSO, 0.1 µM TAL, 1 µM TAL, and PBs treated with DMSO at 14 DPA. Principal component analysis (PCA) showed segregation between treatment groups including PBs and DBs treated with DMSO (Fig. 4A, Fig. S10A). DBs treated with 0.1 or 1 µM TAL had more similar transcriptomes compared to either DMSO-treated PBs or DBs (Fig. 4A, Fig. S10A). However, TAL-treated DBs exhibited transcriptomes more akin to PBs, suggesting a shift towards a more PB-like positional identity. (Fig. 4A, Fig. S10A).

**Figure 4:**
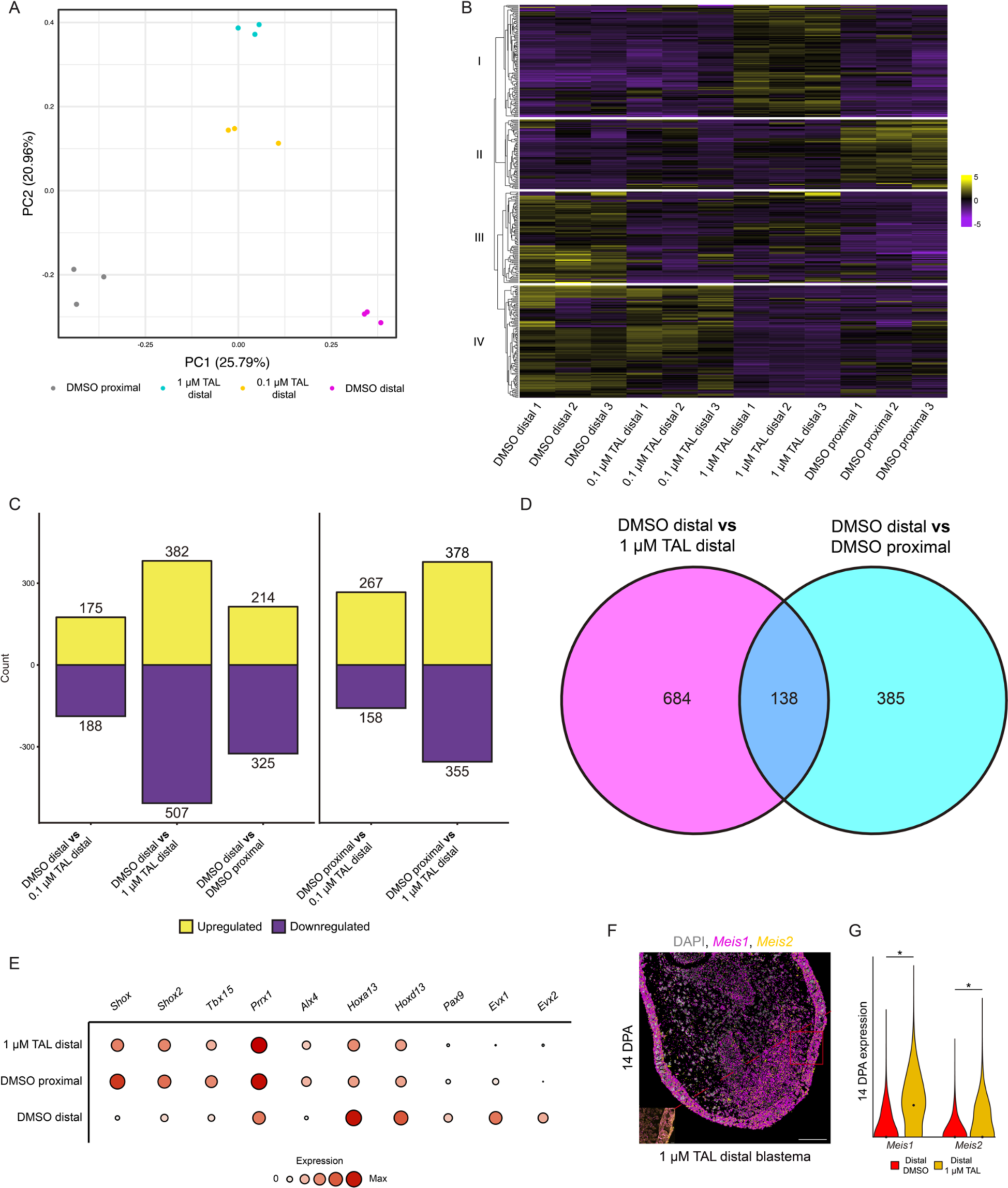
CYP26 inhibition reprograms DBs into a more PB-like identity. (A) PCA of bulk transcriptomes from DBs treated with DMSO, 0.1, or 1 µM TAL and PBs treated with DMSO. (B) Heatmap of the top 371 (padj < 0.01, FC = 1.5) genes expressed in each sample type. Cluster numbers are next to the dendrogram. (C) Bar graphs of significantly upregulated and downregulated genes (padj < 0.1) within each comparison. (D) Venn diagram of overlapping DEGs (padj < 0.1) from DMSO treated DBs vs DMSO treated PBs and DMSO treated DBs vs 1 µM TAL treated DBs. Full gene lists are in Table S6. (E) Selected shared DEGs from (D). (F) HCR-FISH for *Meis1* and *Meis2* in DBs administered 1 µM TAL at 14 DPA. Dashed line indicates amputation plane. Scale bars = 200 µm or 20 µm (inset). (F) HCR-FISH dot quantification for mesenchymal *Meis1* and *Meis2* in DBs treated with DMSO or 1 µM TAL at 14 DPA (n = 3-6, 3.5 cm (HT) animals aged 2.5 months). Axes and analyses as in Fig. 1E. * = p < 0.05.

We classified these transcriptional differences using hierarchical clustering of the top 371 differentially expressed genes (DEGs) (padj < 0.01, FC > 1.5) and observed four clusters (Fig. 4B). In cluster one, we observed RA-responsive genes upregulated by 1 µM TAL, including *Cyp26a1*, *Krt15*, and *Acan* which are known to be upregulated in response to RA (Fig. 4B) (Nguyen et al., 2017; Polvadore and Maden, 2021). Considering these genes were not differentially expressed in PBs and DBs, it is unlikely that they play a role in PD positional identity. Cluster two consisted of genes highly expressed in DMSO-treated PBs, some of which increased with higher TAL concentrations, while cluster three included genes highly expressed in DMSO-treated DBs. TAL treatment generally decreased cluster three gene expression, with the lowest levels in DMSO-treated PBs (Fig. 4B). Cluster four represents a transitional phase between DBs treated with 0.1 µM and 1 µM TAL, with genes highly expressed in DBs treated with DMSO and 0.1 µM TAL, and lowly expressed in DBs treated with 1 µM TAL and PBs treated with DMSO (Fig. 4B). The number of DEGs relative to DMSO treated DBs increases from 363 to 889 (padj < 0.1) as TAL concentration rises from 0.1 or 1 µM, respectively (Fig. 4C). In comparison, DMSO treated PBs contained 539 DEGs (padj < 0.1) relative to DMSO treated DBs (Fig. 4C). The number of DEGs between DMSO treated DBs and 1 µM TAL treated DBs was higher than between DMSO treated DBs and PBs, likely reflecting RA-responsive genes from cluster one not involved in PD patterning. DMSO treated PBs had 425 and 733 DEGs (padj < 0.1) compared to 0.1 or 1 µM TAL treated DBs, showing that while TAL treated DBs adopted a more PB-like identity, differences remained (Fig. 4C).

Comparisons between DMSO treated DBs and DMSO treated PBs or 1 µM TAL treated DBs enabled identification of RA-responsive genes associated with proximal or distal limb identity. We found that 138 DEGs (FDR <0.1) were shared between both comparisons (Fig. 4D, Table S6). Among these genes, we identified several that have known roles in patterning the developing or regenerating limb, including *Pax9* (McGlinn et al., 2005), *Evx1* (Niswander et al., 1994), and *Alx4* (te Welscher et al., 2002) (Fig. 4E). Notably absent were *Meis1* and *Meis2* despite their known roles as RA-responsive genes involved in PD patterning (Bryant et al., 2017; Nguyen et al., 2017; Polvadore and Maden, 2021). Both genes were significantly more highly expressed in the mesenchyme of DBs treated with 1 µM TAL compared to DMSO treated DBs at 14 DPA (Fig. 1D-E, Fig. 4F-G). This indicates that high levels of epithelial expression in blastemas obscured detection by RNA-seq. High epithelial expression also prevented *Cyp26b1* detection by RNA-seq (Fig. 2C-D, Fig. 3E-F), indicating that *Meis1*, *Meis2*, and *Cyp26b1* are involved in patterning and change in response to TAL treatment. Together, our results show that TAL treatment induces DBs to adopt a more proximal positional identity, likely due to RA-responsive patterning genes differentially expressed between PBs and DBs.

### *Shox* is a downstream target of RA involved in stylopod and zeugopod patterning

Among RA-responsive genes that were differentially expressed between PBs and DBs were *Shox* and *Shox2* (Fig. 4E). Shortened limbs are frequently observed in humans with *Shox* haploinsufficiency, which is commonly linked to idiopathic short stature, Turner syndrome, and Leri-Weill dyschondrosteosis (Rao et al., 1997; Shears et al., 1998). While mice lack a functional *Shox* ortholog, *Shox2* mutant mice develop with shortened humeri (Yu et al., 2007). In the axolotl, SHOX and SHOX2 share 73.98% sequence similarity and contain a 100% identical homeodomain (Fig. S11A). *Shox* and *Shox2* have previously been noted for their potential role in proximal positional identity during axolotl limb regeneration (Bryant et al., 2017), and were more epigenetically accessible in uninjured connective tissue (CT) cells of the stylopod compared to those of the autopod (Kawaguchi et al., 2024). These findings suggest that *Shox* and *Shox2* are involved in maintaining and reestablishing proximal limb positional information during limb regeneration.

To explore the role of *Shox* and *Shox2* in patterning the PD axis, we visualized *Shox* and *Shox2* expression in developing limb buds from stage 44-47 (Fig. 5A). During limb development, we observed high levels of *Shox* throughout the mesenchyme of stage 44 limb buds with *Shox2* localized to the posterior mesenchyme. At stage 47, *Shox2* expression remained in the posterior limb mesenchyme but was proximally biased (Fig. 5A). *Shox* was exclusively expressed in the proximal mesenchyme of the stage 47 limb bud, leaving a *Hoxa13*^+^ zone of distal mesenchymal cells devoid of *Shox* expression (Fig. 5A). These results suggest that *Shox* is involved in establishing proximal limb identity during limb development.

**Figure 5:**
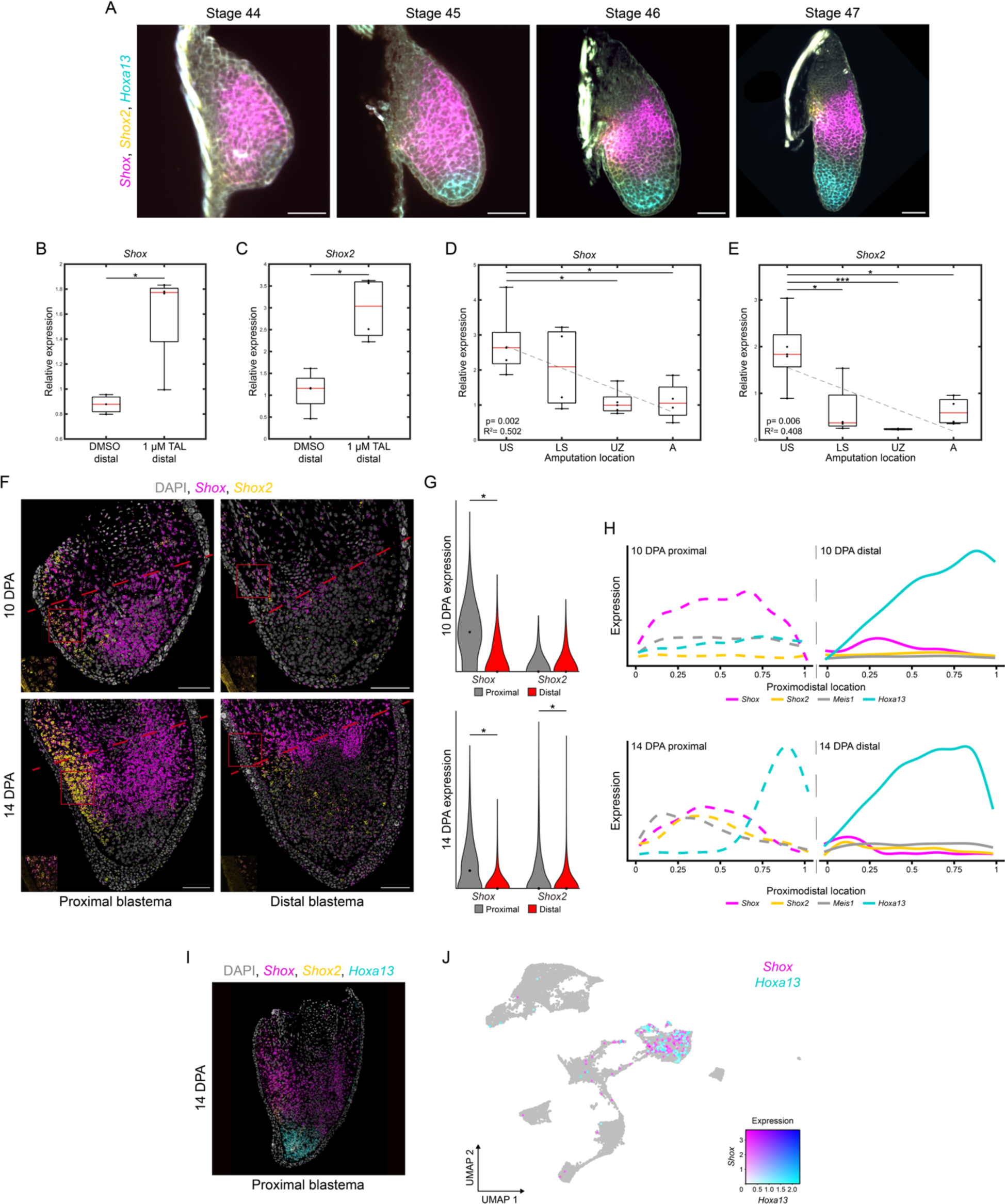
*Shox* and *Shox2* mark proximal and posterior positional identity. (A) Whole mount HCR-FISH for *Shox*, *Shox2*, and *Hoxa13* in stage 44-47 developing limb buds. Scale bars = 100 µm. (B-C) qRT-PCR of *Shox* and *Shox2* in DMSO or 1 µM TAL treated DBs (n = 4, 4 blastemas per sample, 3.5 cm (HT) animals aged 2.5 months, 10 DPA). Each gene was normalized to *Ef1a* and the groups were analyzed using a two-tailed t-test. * = p < 0.05. (D-E) qRT-PCR of *Shox* (D) and *Shox2* (E) at different PD amputation locations (n = 3-6, 4-5 blastemas per sample, 3.5 cm (HT) animals aged 2.5 months, 10 DPA). Analyses as in Fig. 1B-C. * = p < 0.05, *** = p < 0.001. (F) HCR-FISH for *Shox* and *Shox2* in PBs and DBs at 10 and 14 DPA. Dashed lines indicate amputation plane. Scale bars = 200 µm or 20 µm (inset). (G) HCR-FISH dot quantification for mesenchymal *Shox* and *Shox2* in PBs and DBs at 10 and 14 DPA (n = 3-6, 3.5 cm (HT) animals aged 2.5 months). Axes and analyses as in Fig. 1E. * = p < 0.05. (H) PD intensity plots for mesenchymal *Shox*, *Shox2*, *Meis1*, and *Hoxa13* in PBs and DBs at 10 and 14 DPA. Axes and analyses as in Fig. 1F. (I) HCR-FISH for *Shox*, *Shox2*, and *Hoxa13* in PBs at 14 DPA. Dashed line indicates amputation plane. Scale bar = 200 µm or 20 µm (inset). (J) UMAP of *Shox*^+^ and *Hoxa13*^+^ cells in DBs from reanalyzed scRNA-seq dataset (Li et al., 2021).

We next examined *Shox* and *Shox2* expression following limb amputation. In agreement with our RNA-seq results, *Shox* and *Shox2* expression increased in DBs upon TAL treatment and were more highly expressed in US amputations compared to autopod amputations (Fig. 5B-E). *Shox* expression appeared to decrease incrementally in more distal amputations whereas *Shox2* expression in LS blastemas had similar expression as autopod blastemas (Fig. 5D-E). Furthermore, *Shox* and *Shox2* were primarily expressed in mesenchymal cells with little expression in any other cell type (Fig. S11B-C). As in limb development, *Shox2* expression in PBs and DBs at each time point was localized proximally and posteriorly in mesenchymal cells (Fig. 5F-H, Fig. S11D). *Shox2* expression levels in PBs and DBs at 10 DPA were similar, but by 14 DPA *Shox2* was significantly more highly expressed in the proximal and posterior mesenchyme of PBs (Fig. 5F-H). This may indicate that *Shox2* has a role in patterning both the PD and AP axis during limb development and regeneration. *Shox* was lowly expressed in the mesenchyme of PBs and DBs at 7 DPA (Fig. S11D). By 10 DPA, *Shox* expression spread throughout the mesenchyme of PBs and was significantly more highly expressed than in DBs (Fig. 5F-H). The limited areas of *Shox* expression in DBs at 10 and 14 DPA seemed to be associated with the uninjured skeletal elements (Fig. 5F-H). At 14 DPA, *Shox* remained significantly more highly expressed in PBs but was restricted to the proximal mesenchyme, leaving a distal subset of *Hoxa13*^+^ cells devoid of *Shox* expression (Fig. 5F-I). *Shox*^+^ and *Hoxa13*^+^ cells are mutually exclusive (Fig. 5I-J), suggesting that *Shox* is not involved in autopod formation.

Given the spatiotemporal expression patterns of *Shox* (Fig. 5F-H), *Meis1*, and *Hoxa13* (Fig. 1), it is possible that *Shox* is activated by *Meis1* and repressed by *Hoxa13*. Given that *Meis1* is RA-responsive (Fig. 4F-G), this overlap in expression may indicate that mesenchymal *Shox* is activated by RA via *Meis1*. Conversely, *Hoxa13* is repressed by RA (Fig. 3G, Fig S8C-D), suggesting that *Hoxa13* expression creates the distal limit for *Shox*.

### *Shox* is required for stylopodial and zeugopodial endochondral ossification

Our gene expression results led us to hypothesize that *Shox* has a role in establishing stylopod and zeugopod, not autopod, positional identity during both limb development and regeneration. To test this hypothesis, we utilized CRISPR/Cas9 to genetically inactivate *Shox*. We targeted *Shox* with two sgRNAs specifically designed against exons 1 and 2 and simultaneously injected these sgRNAs into axolotl embryos to create mosaic F0 *Shox* knockout animals (*Shox* crispants) (Fig. 6A). NGS genotyping analyses of 10 *Shox* crispants indicated that both targeted loci were highly mutated, ranging from 75.15-97.62% of all sequenced alleles being mutated across each animal (Fig. S12). Furthermore, these animals developed to adulthood, enabling us to examine the role of *Shox* during limb development and regeneration. *Shox* crispants developed significantly smaller limbs than controls with significantly smaller stylopods and zeugopods (Fig. 6B-C). Interestingly, autopod length was unaffected in *Shox* crispants compared to controls (Fig. 6C), showing that *Shox* is critical for stylopod and zeugopod development but dispensable for autopod development. A similar phenotype was observed in the limbs of *Shox2* knockout mice, where chondrocytes in shortened stylopods and zeugopods failed to proliferate and mature, preventing endochondral ossification (Yu et al., 2007). We observed that while skeletal elements in control stylopods and zeugopods were partially calcified, those from *Shox* crispants appeared to lack calcification (Fig. 6D). Additionally, chondrocytes from control stylopods demonstrated clear progression from proliferation to hypertrophy before calcifying (Fig. 6E). In contrast, chondrocytes from *Shox* crispants failed to proliferate, appearing to remain as reserve cartilage through adulthood (Fig. 6E). Considering autopod size was not impacted in *Shox* crispant limbs, we next examined the digits of *Shox* crispants limbs to determine if endochondral ossification in autopodial elements was disrupted. Chondrocytes from both control and *Shox* crispant digits underwent phenotypically normal endochondral ossification, suggesting that while essential for more proximal limb segments, SHOX is not required for autopodial skeletal maturation (Fig. 6E). This indicates that proximal and distal skeletal elements have disparate transcriptional programs responsible for endochondral ossification. Collectively, our data indicate that *Shox* is required for chondrocyte maturation within proximal skeletal elements. During limb development, however, many *Shox*^+^ cells are *Sox9*^-^, suggesting that *Shox* may also be involved in patterning non-chondrogenic mesenchymal cells (Fig. 6F).

**Figure 6:**
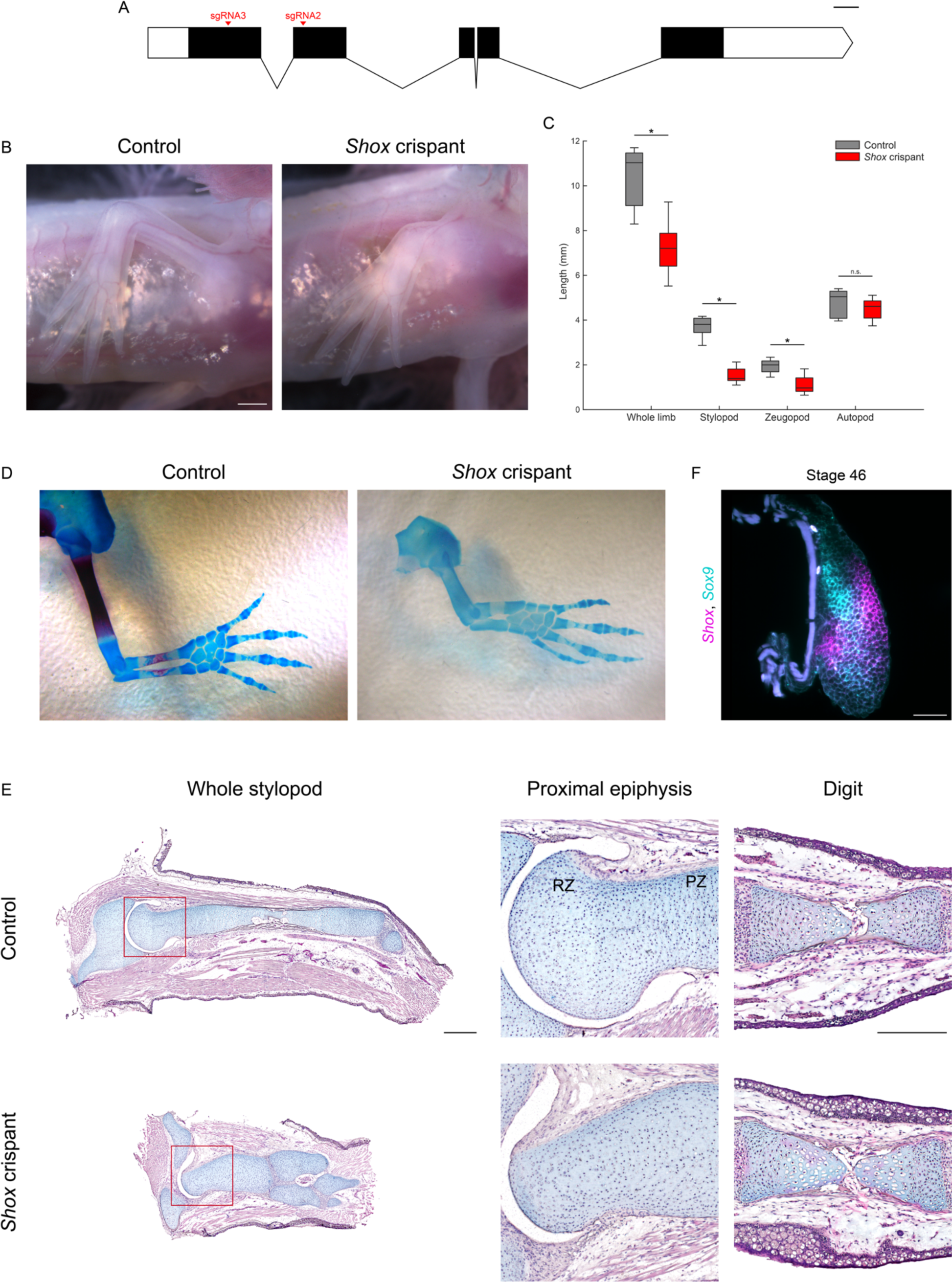
*Shox* crispants show defects in endochondral ossification of proximal limb skeletal elements. (A) Schematic of the *Shox* genomic landscape. Introns reduced 50X for visibility. Scale bar = 100 bp. (B) Brightfield images of control and *Shox* crispant limbs (3.5 cm (HT) animals aged 2.5 months). Scale bar = 1 mm. (C) Skeletal element quantification in control and *Shox* crispant limbs (n = 8 per group, 7.5 cm (HT) animals aged 6 months). n.s. = no statistical difference, * = p < 0.05. (D) Alcian blue and alizarin red stain of adult control and *Shox* crispant limbs (12 cm (HT) animals aged 10 months). (E) H&E&A of whole stylopods, proximal epiphyses, and digits from controls and *Shox* crispants (8 cm (HT) animals aged 7 months). RZ = resting zone, PZ = proliferative zone. Stylopod scale bar = 1 mm, Digit scale bar = 0.5 mm. (F) Whole mount HCR-FISH for *Shox* and *Sox9* in a stage 46 developing limb. Scale bar = 100 µm.

### *Shox* is not required for limb regeneration but is essential for proximal limb patterning

Finally, we found that *Shox* was dispensable for limb regeneration, as *Shox* crispants successfully regenerated limbs and progressed though typical stages, including blastema and palate formation, without obvious abnormalities (Fig. 7A). *Shox* crispant limbs remained shortened after fully regenerating, suggesting that *Shox* is dispensable for limb regeneration but crucial for patterning the regenerating stylopodial and zeugopodial elements (Fig. 7A). However, like in limb development, not all *Shox*^+^ cells were *Sox9*^+^, suggesting that *Shox* has an additional role in patterning mesenchymal cells outside the chondrocyte lineage during limb regeneration (Fig. 7B-C). We then investigated if *Shox* crispant blastemas show abnormalities in PD patterning gene expression and found no change in the spatial expression of *Meis1* and *Hoxa13* (Fig. 1D, J, Fig. 7D), consistent with results in *Shox2* KO mice (Yu et al., 2007). This observation, along with *Meis1* and *Shox* colocalization (Fig. 5H), suggests that *Shox* acts downstream of *Meis1* to establish proximal limb positional identity. Considering *Meis1* is RA-responsive and limb-specific *Meis* KO mice lack proximal skeletal elements (Delgado et al., 2020), RA likely directs proximal endochondral ossification through *Shox* via *Meis1*. Consistent with this, administering 1 µM TAL to *Shox* crispant DBs lead to duplication of shortened proximal elements (Fig. 7E), showing that *Shox* crispant limbs respond to RA but cannot properly pattern proximal elements.

**Figure 7:**
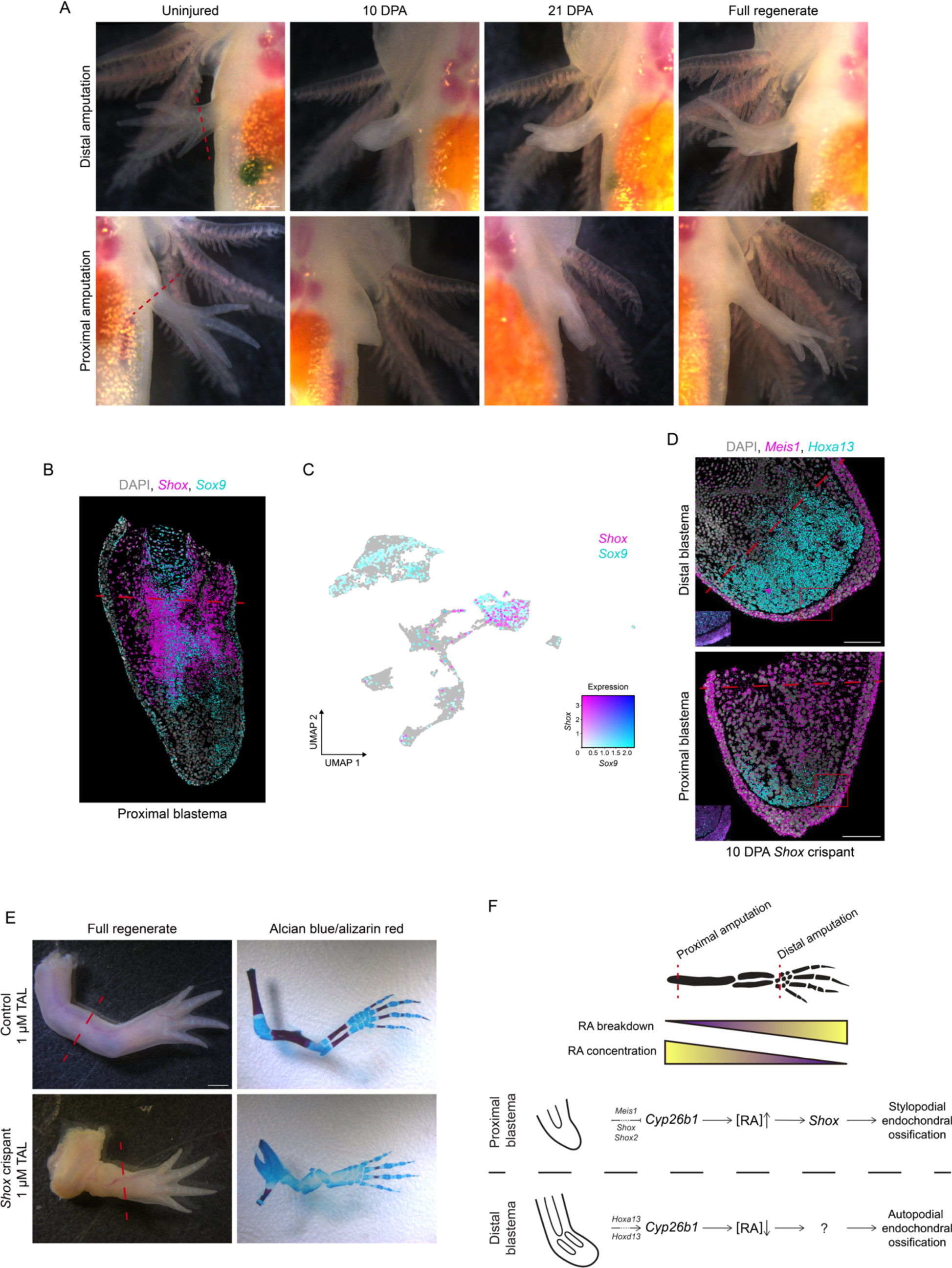
*Shox* is dispensable for limb regeneration but required for PD patterning. (A) Regeneration time course of PBs and DBS in *Shox* crispants. Scale bar = 1 mm. (B) HCR-FISH for *Shox* and *Sox9* in PBs at 21 DPA. Dashed line indicates amputation plane. Scale bars = 200 µm or 20 µm (inset). (C) UMAP of *Shox*^+^ and *Sox9*^+^ cells in DBs from reanalyzed scRNA-seq dataset (Li et al., 2021). (D) HCR-FISH for *Meis1* and *Hoxa13* in *Shox* crispant PBs and DBs at 10 DPA. Dashed lines indicate amputation plane. Scale bars = 200 µm or 20 µm (inset). (E) Brightfield images of regenerates and skeletal structures of control or *Shox* crispant limbs treated with 1 µm TAL. Scale bar = 2 mm. (F) Model for PD patterning during limb regeneration.

## Discussion

Our study shows that endogenous RA is required for PD limb patterning during regeneration. We propose that *Cyp26b1-*mediated RA breakdown, not RA synthesis or RAR expression, determines PD positional identity by setting the RA signaling levels in the blastema, activating or repressing RA signaling (Fig. 7F). Moreover, CT cells in the uninjured limb have inherent, epigenetically encoded positional identity (Kawaguchi et al., 2024). Upon amputation, these cells dedifferentiate into a limb bud-like state while retaining their PD positional memory (Gerber et al., 2018). We propose that dedifferentiated blastema cells modify *Cyp26b1* expression depending on their positional memory, which adjusts RA signaling levels to the appropriate PD location and regulates genes that convey PD positional identity. Elevated RA levels in PBs activate *Shox*, promoting endochondral ossification in proximal skeletal elements. In contrast, reduced RA levels in DBs leads a *Shox*-independent mechanism for endochondral ossification of distal skeletal elements. Our results show that RA signaling levels exert segment-specific effects on during skeleton regeneration (Fig. 7F).

The model that we propose explains how positional identity is determined among cells at two spatial scales: the entire limb PD axis and PD axes within different limb segments. *Cyp26b1* expression was graded across the limb PD axis, with similar levels of expression observed between blastemas that formed from different, but spatially juxtaposed limb segments (e.g. LS and UZ blastemas). However, within limb segments, *Cyp26b1* expression differed considerably between blastemas that formed from spatially disparate locations (e.g. US and LS blastemas). Our model suggests that positional identity is conveyed in a segment specific manner, as *Shox* expression is mutually exclusive from *Hoxa13* and *Shox* perturbation affects only stylopodial and zeugopodial skeletal elements. Further evidence for this comes from *Hoxa13* knockout newts that fail to regenerate autopods (Takeuchi et al., 2022). This may imply that each limb segment requires a specific threshold of positional values determined by RA signaling levels, as posited by the French Flag model (Wolpert, 1969). Once this threshold is met, blastema cells can create intra-segmental positional values based on RA signaling levels, enabling fine-tuning of limb segment morphology. It may also explain why half-segment duplications are observed following treatment with lower concentrations of TAL or RA during limb regeneration (Maden, 1982; Thoms and Stocum, 1984).

Further studies are needed to identify upstream activators of *Cyp26b1* during amphibian limb regeneration. We observed that *Cyp26b1* exhibited a similar spatiotemporal expression pattern as *Hoxa11* and *Hoxa13*, providing circumstantial evidence that 5’ Hox genes regulate *Cyp26b1* expression. Previous studies on mouse limb development have suggested that AER-derived FGFs activate *Cyp26b1* instead of 5’ Hox genes to create a domain of distal identity while simultaneously interacting with SHH to promote distal outgrowth (Probst et al., 2011). Indeed, *Fgf8* expression in *Cyp26b1*^-/-^ limbs does not appear to be impacted by the loss of CYP26B1 function (Yashiro et al., 2004). However, this may differ during limb regeneration as the SHH-FGF feedback loop is required for blastema distal outgrowth and proliferation (Nacu et al., 2016), and neither *Fgf8* nor *Shh* were differentially expressed between PBs and DBs in our RNA-seq. For these reasons, it seems more likely that *Cyp26b1* is regulated by 5’ Hox genes including *Hoxa11* and *Hoxa13* during limb regeneration.

An outstanding question is why RA would be synthesized in the regenerating limb only to be degraded depending on amputation location. While our work does not address this complexity directly, we speculate that RA has several roles during limb regeneration outside of providing proximal limb positional identity. RA is important for directing nerves to their targets, including epithelial and neuromuscular junctions (Dmetrichuk et al., 2005). Furthermore, RA plays an essential role in replenishing epithelial cells during physiological growth (Zasada and Budzisz, 2019), which may contribute to the scarless wound healing following injuries in salamanders (Seifert et al., 2012).

How a cell conveys its PD positional identity to neighboring cells in response to RA to coordinate patterning in the blastema remains an unanswered question in the field. Several lines of evidence suggest that differences in adhesivity and cell adhesion molecules (CAMs) differentiate PBs from DBs (Nardi and Stocum, 1984). These differences in adhesivity can be modified by RA, suggesting that RA signaling levels are important for generating a gradient of adhesive properties in PBs and DBs (Crawford and Stocum, 1988; Johnson and Scadding, 1992). In agreement with these studies, two CAMs, TIG1 and PROD1, have been identified that are RA-responsive and modify adhesivity during limb regeneration (da Silva et al., 2002; Oliveira et al., 2022). Neither of these CAMs appeared to be RA-responsive or differentially expressed in PBs and DBs, despite their importance in directing cell adhesivity during limb regeneration. Nonetheless, our RNA-seq dataset should serve as a helpful resource for identifying other RA-responsive CAMs that are differentially expressed in PBs and DBs.

## Limitations of the study

While TAL is often used to study endogenous RA levels, chemical inhibitors are not tissue specific. TAL also inhibits all CYP26 paralogs, not just CYP26B1. Our results have shown that *Cyp26a1* and *Cyp26c1* are lowly expressed or not expressed and as such, CYP26B1 should be the primary paralog affected. However, future studies would benefit from developing a mesenchyme specific *Cyp26b1* KO to ensure that it is the primary driver of RA breakdown during limb regeneration.

## Acknowledgements

The authors thank Guoxin Rong for his imaging expertise and assistance with microscopy. Additionally, we thank Prayag Murawala for providing transgenic animals and the Ambystoma Genetic Stock Center for non-transgenic animals. We thank Malcolm Maden for his critical analysis of the manuscript. We finally thank the Institute for Chemical Imaging of Living Systems at Northeastern University for consultation and imaging support.

## Funding

The work from this paper was funded by NIH grant R01HD099174 and by NSF grants 1558017 and 1656429. Non-transgenic animals were obtained from the Ambystoma Genetic Stock Center funded through NIH grant P40-OD019794.

## Declaration of generative AI and AI-assisted technologies in the writing process

During the preparation of this work the authors used ChatGPT to aid in sentence structure and proofread for grammatical errors. After using this tool, the authors reviewed and edited the content as needed and take full responsibility for the content of the publication.

---

**Figure S1:**
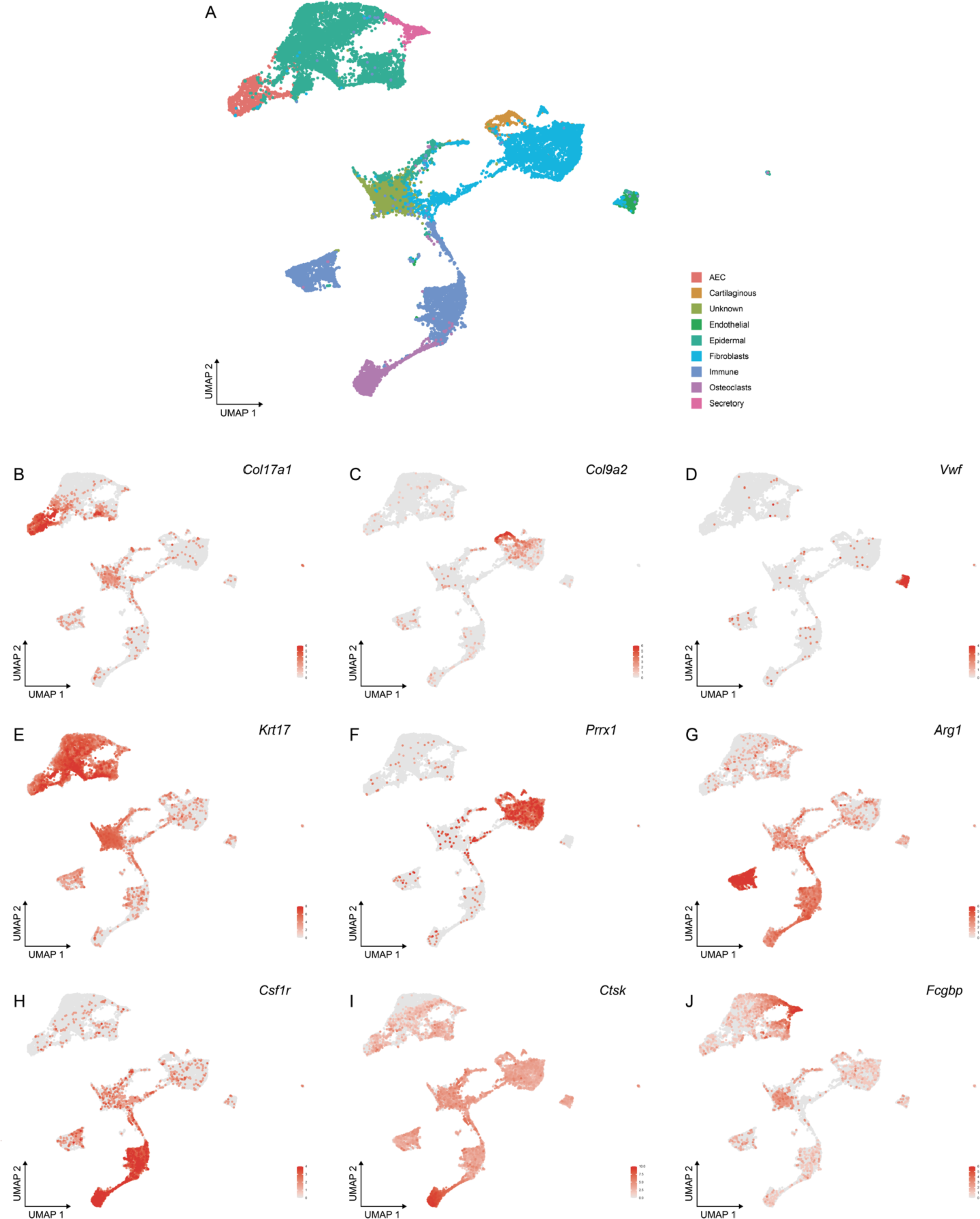
Cluster identification in reanalysis of DBs from Li et al. 2021. (A) Clustering of 7, 14, and 22 DPA blastemas from Li et al. 2021 with 9 clusters marked by designated genes. (B) *Col17a1* marks the apical epithelial cap (AEC). (C) *Col9a2* marks chondrocytes. (D) *Vwf* marks endothelial cells. (E) *Krt17* marks an unknown cell population. (F) *Prrx1* marks dedifferentiated limb CT cells. (G-H) *Arg1* and *Csf1r* mark immune cells. (I) *Ctsk* marks osteoclasts. (J) *Fcgbp* marks secretory cells.

**Figure S2:**
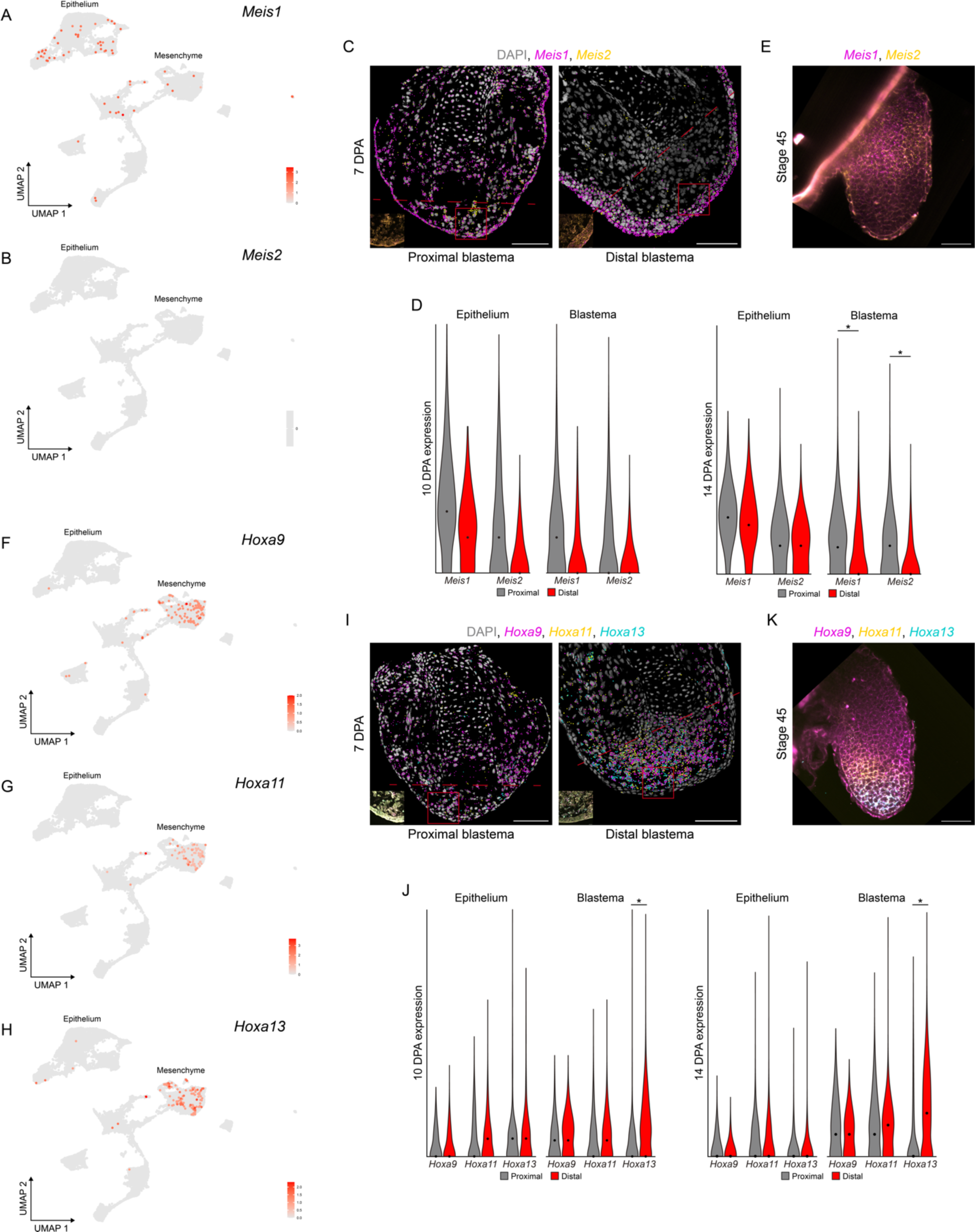
Additional characterization of PD patterning gene expression. (A) UMAP showing *Meis1* in DBs at 7, 14, and 22 DPA. (B) UMAP showing *Meis2* expression was undetected in DBs at 7, 14, or 22 DPA. (C) HCR-FISH for *Meis1* and *Meis2* in PBs and DBs at 7 DPA. Dashed lines indicate amputation plane. Scale bars = 200 µm or 20 µm (inset). (D) HCR-FISH dot quantification for mesenchymal or epithelial *Meis1* and *Meis2* in PBs and DBs at 10 and 14 DPA (n = 3-6, 3.5 cm (HT) animals aged 2.5 months). Axes and analyses as in Fig. 1E. * = p < 0.05. (E) Whole mount HCR-FISH for *Meis1* and *Meis2* expression in whole mount developing limb buds at stage 45. The image represents a single, 2D z-plane within a 3D image stack. Scale bar = 100 µm. (F-H) UMAP showing *Hoxa9* (F), *Hoxa11* (G), and *Hoxa13* (H) in DBs at 7, 14, and 22 DPA. (I) HCR-FISH for *Hoxa9*, *Hoxa11*, and *Hoxa13* in PBs and DBs at 7 DPA. Dashed lines indicate amputation plane. Scale bars = 200 µm or 20 µm (inset). (J) HCR-FISH dot quantification in the epithelium and whole blastema for *Hoxa9*, *Hoxa11*, and *Hoxa13* in PBs and DBs at 10 and 14 DPA (n = 3-6, 3.5 cm (HT) animals aged 2.5 months). Axes and analyses as in Fig. 1E. * = p < 0.05. (K) Whole mount HCR-FISH for *Hoxa9*, *Hoxa11*, and *Hoxa13* in whole mount developing limb buds at stage 45. The image represents a single, 2D z-plane within a 3D image stack. Scale bar = 100 µm. ScRNA-seq data were reanalyzed from a previously published dataset (Li et al., 2021).

**Figure S3:**
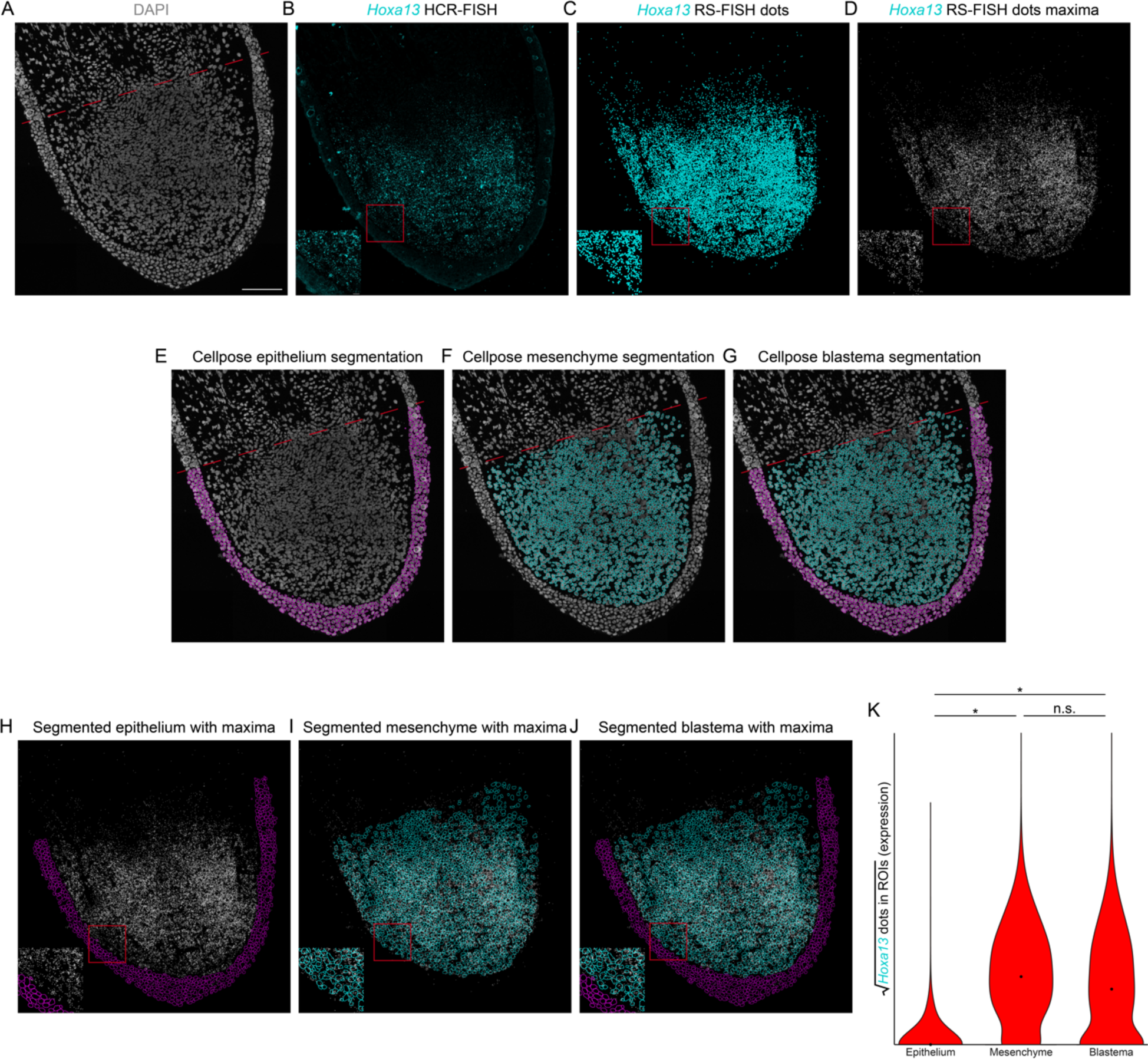
HCR-FISH dot quantification workflow. (A) DAPI channel for a representative DB replicate at 14 DPA. Scale bar = 200 µm. (B) Raw imaging data for *Hoxa13* expression. Inset scale bar = 20 µm. (C) Pseudodots obtained by RS-FISH (Bahry et al., 2022). (D) Maxima-converted pseudodots, each containing a pixel value of 255. (E-G) Cellpose-generated segmentation of epithelium (E), mesenchyme (F), and whole blastema (G) from DAPI channel in panel A (Stringer et al., 2021). (H-J) Segmentation for epithelium (H), mesenchyme (I), and whole blastema (J) overlayed on atop maxima from panel D. (K) HCR-FISH dot quantification in each tissue type for *Hoxa13* expression in DBs at 14 DPA (n = 3-6, 3.5 cm (HT) animals aged 2.5 months). The sum of the pixel values within each cell was divided by 255 to obtain the total number of HCR-FISH dots within a cell. These values were then square root normalized for visualization in violin plots. Groups were analyzed using a clustered Wilcoxon rank sum test according to the Datta-Satten method. n.s. = no statistical difference, * = p < 0.05.

**Figure S4:**
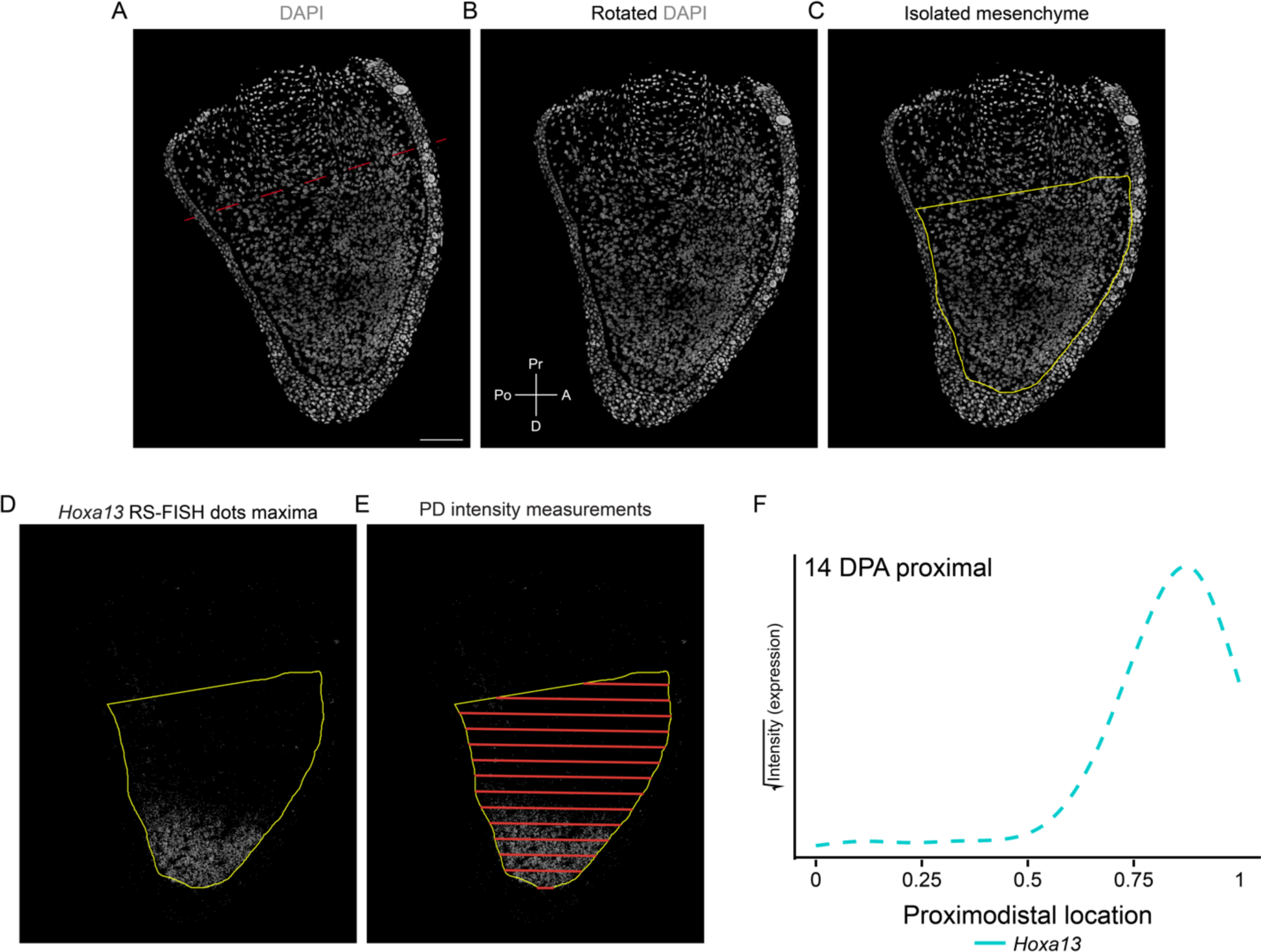
Workflow for generating HCR-FISH PD intensity plots. (A) DAPI channel for a representative PB replicate at 14 DPA. Scale bar = 200 µm. (B) Rotated DAPI channel from panel A. Pr = proximal, A = anterior, D = distal, Po = posterior. (C) Rotated DAPI image with mesenchyme outlined. (D) Mesenchyme outline with maxima-converted pseudodots. Pseudodots obtained via workflow outlined in Figure S3. (E) Intensity measurements obtained continuously along the proximodistal axis. (F) Intensity plots for *Hoxa13* expression along the PD axis within the mesenchyme of PBs at 14 DPA. Lines represent average signal intensity (expression) along a normalized PD axis across each sample.

**Figure S5:**
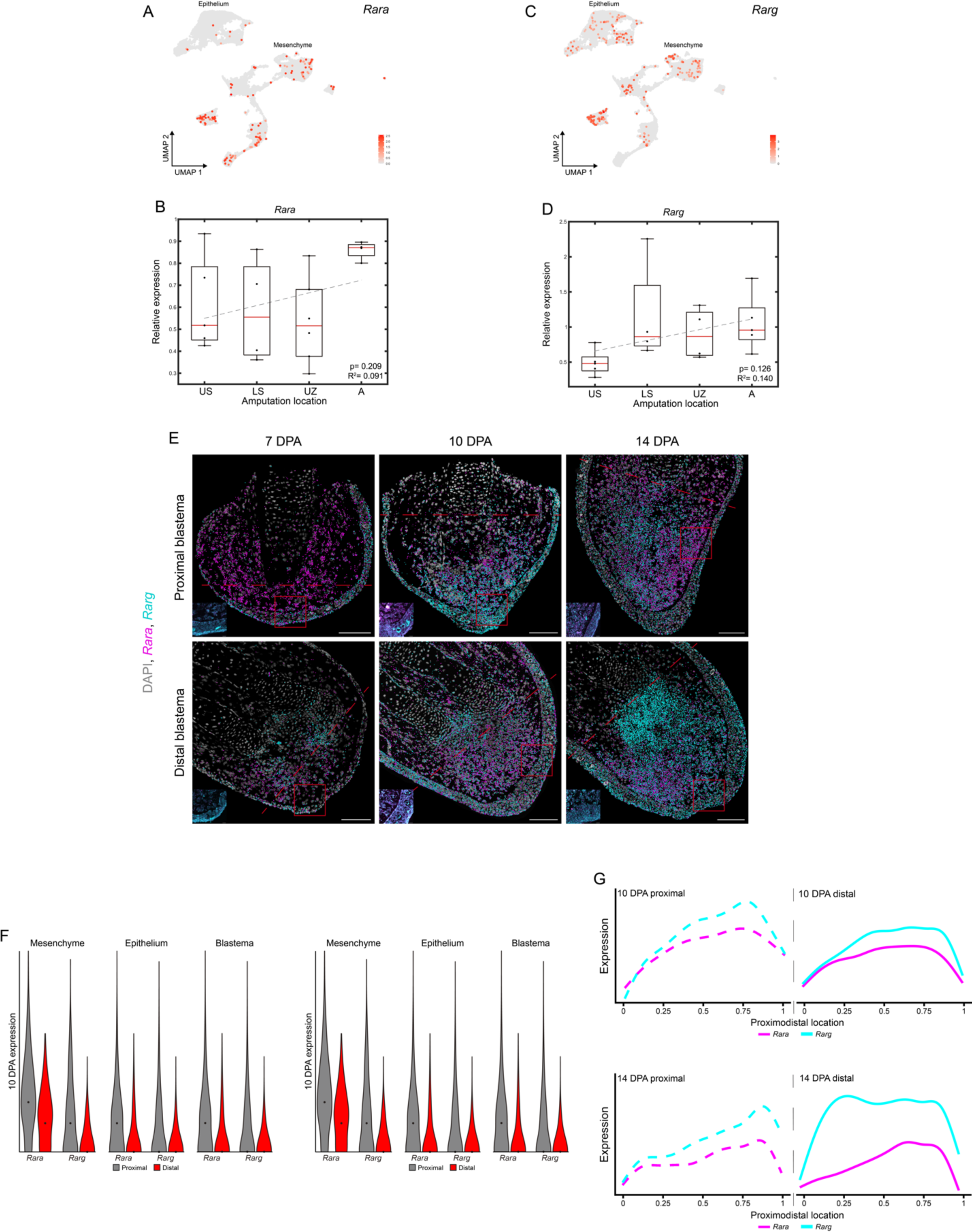
*Rar* expression along the regenerating PD axis. (A) UMAP showing *Rara* in DBs at 7, 14, and 22 DPA. (B) qRT-PCR quantification of *Rara* at different amputation locations along the PD axis (n = 3-6, 4-5 blastemas pooled per sample, 3.5 cm (HT) animals aged 2.5 months, blastemas collected at 10 DPA). Analyses as in Fig. 1B-C. (C) UMAP showing *Rarg* in DBs at 7, 14, and 22 DPA. (D) qRT-PCR quantification of *Rarg* at different amputation locations along the PD axis (n = 3-6, 4-5 blastemas pooled per sample, 3.5 cm (HT) animals aged 2.5 months, blastemas collected at 10 DPA). Analyses as in Fig. 1B-C. (E) HCR-FISH for *Rara* and *Rarg* in PBs and DBs at 7, 10, and 14 DPA. Dashed lines indicate amputation plane. Scale bars = 200 µm or 20 µm (inset). (F) HCR-FISH dot quantification for *Rara* and *Rarg* expression in the mesenchyme, epithelium, and whole blastema of PBs and DBs at 10 DPA (n = 3-6, 3.5 cm (HT) animals aged 2.5 months). Axes and analyses as in Fig. 1E. (L) PD Intensity plots for mesenchymal *Rara* and *Rarg* in PBs and DBs at 10 and 14 DPA (n = 3-6, 3.5 cm (HT) animals aged 2.5 months). Axes and analyses as in Fig. 1F.

**Figure S6:**
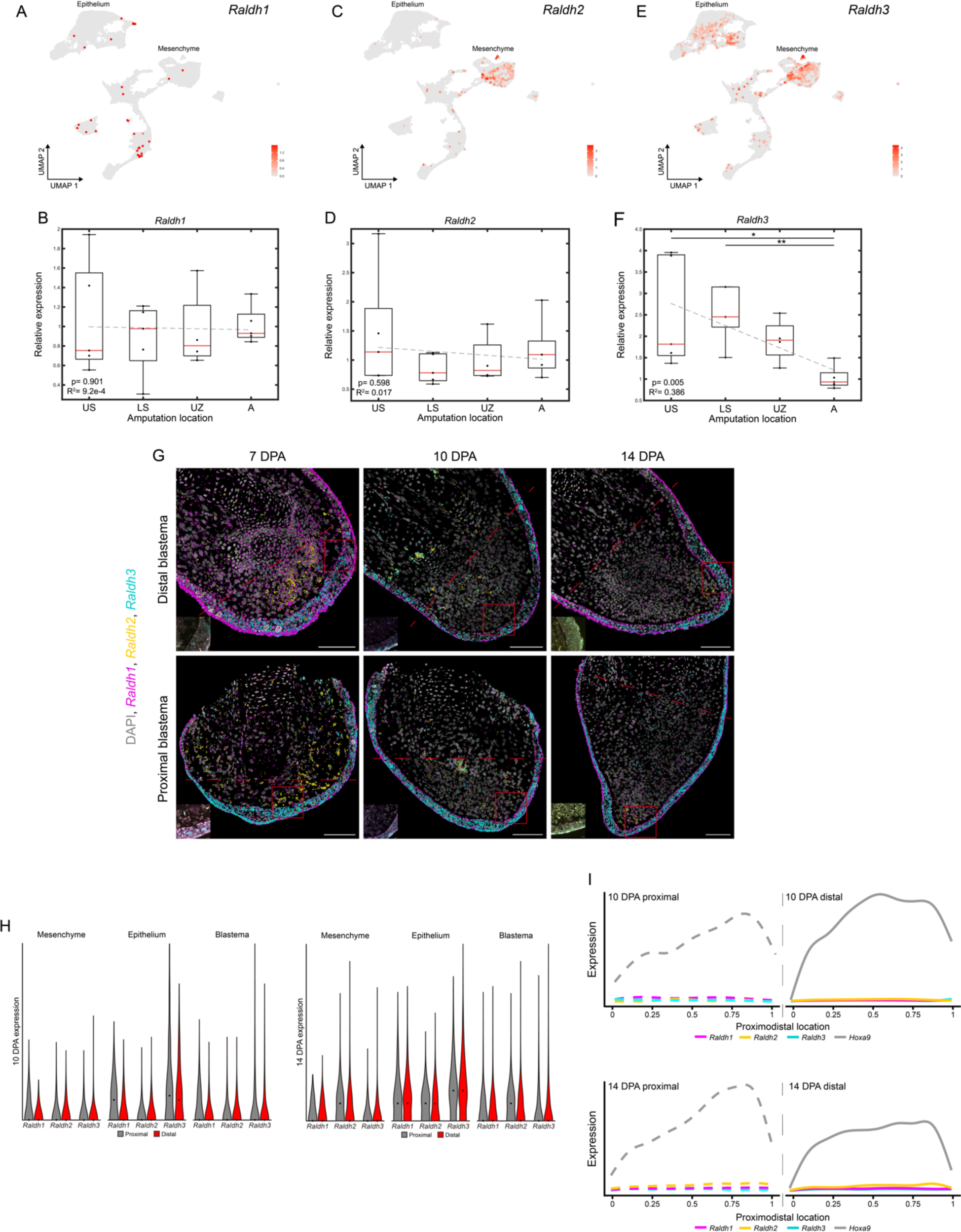
*Raldh* expression along the regenerating PD axis. (A) UMAP showing *Raldh1* in DBs at 7, 14, and 22 DPA. (B) qRT-PCR quantification of *Raldh1* at different amputation locations along the PD axis (n = 3-6, 4-5 blastemas pooled per sample, 3.5 cm (HT) animals aged 2.5 months, blastemas collected at 10 DPA). Analyses as in Fig. 1B-C. (C) UMAP showing *Raldh2* in DBs at 7, 14, and 22 DPA. (D) qRT-PCR quantification of *Raldh2* at different amputation locations along the PD axis (n = 3-6, 4-5 blastemas pooled per sample, 3.5 cm (HT) animals aged 2.5 months, blastemas collected at 10 DPA). Analyses as in Fig. 1B-C. * = p < 0.05, ** = p < 0.01. (E) UMAP showing *Raldh3* in DBs at 7, 14, and 22 DPA. (F) qRT-PCR quantification of *Raldh3* at different amputation locations along the PD axis (n = 3-6, 4-5 blastemas pooled per sample, 3.5 cm (HT) animals aged 2.5 months, blastemas collected at 10 DPA). Analyses as in Fig. 1B-C. (G) HCR-FISH for *Raldh1*, *Raldh2*, and *Raldh3* in PBs and DBs at 7, 10, and 14 DPA. Dashed lines indicate amputation plane. Scale bars = 200 µm or 20 µm (inset). (H) HCR-FISH dot quantification for *Raldh1*, *Raldh2*, and *Raldh*3 in the mesenchyme, epithelium, and whole blastema of PBs and DBs at 10 and 14 DPA (n = 3-6, 3.5 cm (HT) animals aged 2.5 months). Axes and analyses as in Fig. 1E. (I) PD intensity plots for mesenchymal *Raldh1*, *Raldh2*, *Raldh3*, and *Hoxa9* in PBs and DBs at 10 and 14 DPA (n = 3-6, 3.5 cm (HT) animals aged 2.5 months). *Hoxa9* was included as a reference for a more highly expressed gene in the mesenchyme. Axes and analyses as in Fig. 1F.

**Figure S7:**
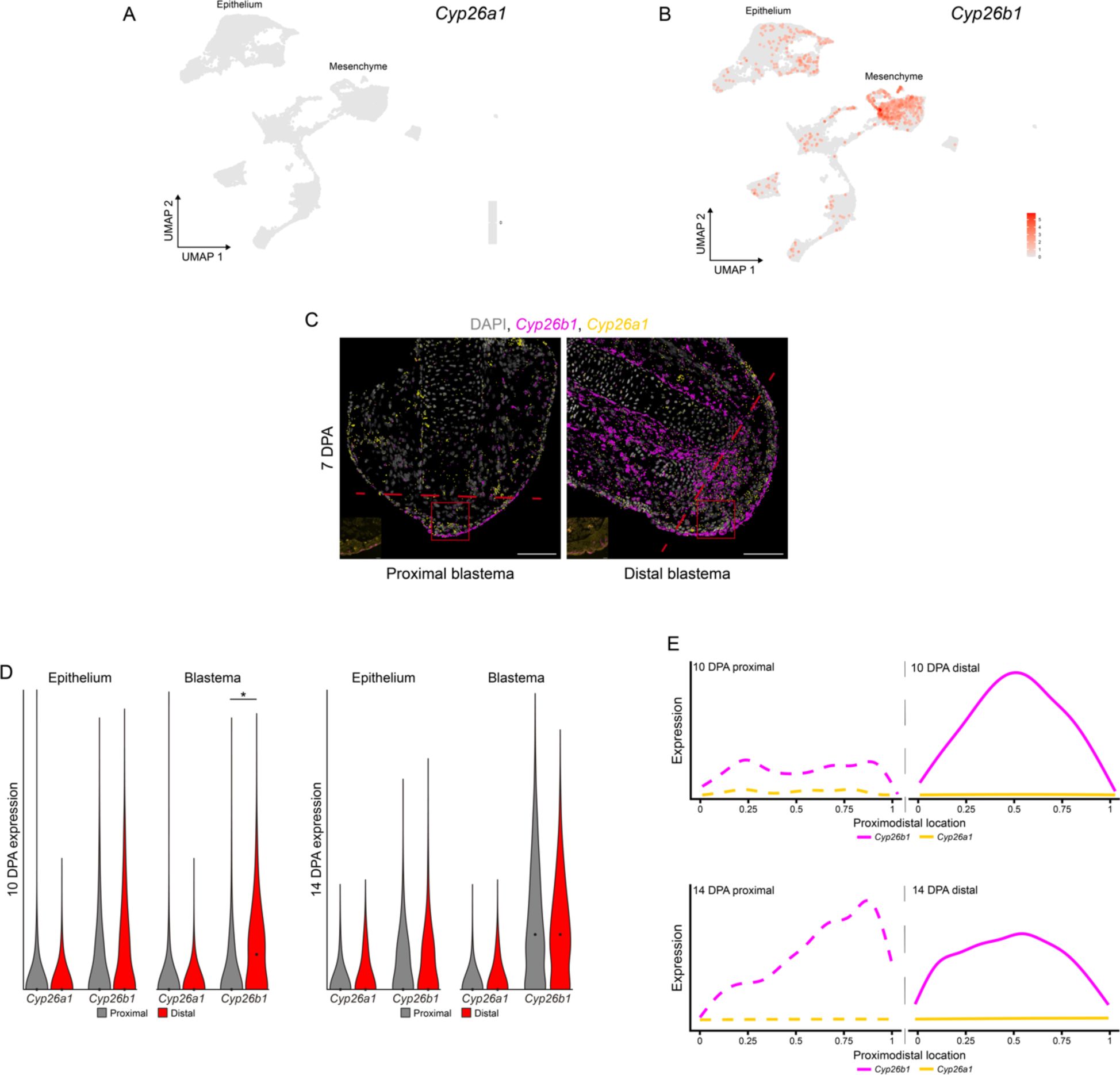
Additional characterization of *Cyp26a1* and *Cyp26b1* expression. (A) UMAP showing *Cyp26a1* expression was undetected in DBs at 7, 14, or 22 DPA. (B) UMAP showing *Cyp26b1* within DBs at 7, 14, and 22 DPA. (C) HCR-FISH for *Cyp26a1* and *Cyp26b1* in PBs and DBs at 7 DPA. Dashed lines indicate amputation plane. Scale bars = 200 µm or 20 µm (inset). (D) HCR-FISH dot quantification in the epithelium and whole blastema for *Cyp26a1* and *Cyp26b1* expression in PBs and DBs at 10 and 14 DPA (n = 3-6, 3.5 cm (HT) animals aged 2.5 months). Axes and analyses as in Fig. 1E. * = p < 0.05. (E) PD intensity plots for *Cyp26a1* and *Cyp26b1* in PBs and DBs at 10 and 14 DPA (n = 3-6, 3.5 cm (HT) animals aged 2.5 months). Axes and analyses as in Fig. 1F.

**Figure S8:**
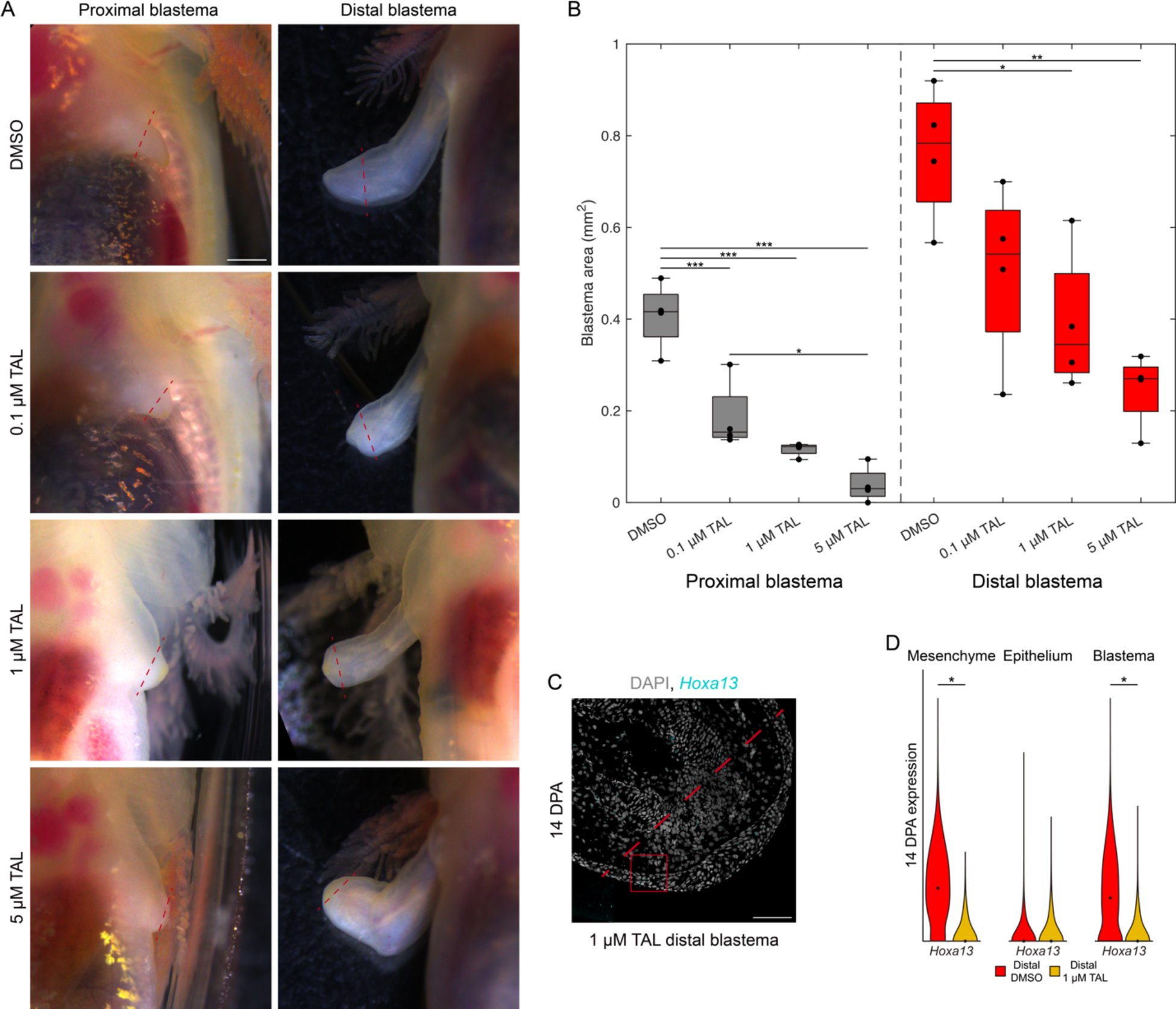
Additional characterization of the impact of TAL during limb regeneration. (A) Brightfield images of PBs and DBs treated with DMSO or 0.1, 1, or 5 µM TAL at 14 DPA. Dashed lines indicate amputation plane. Scale bar = 1 mm. (B) Quantification of blastema area for PBs and DBs treated with DMSO or 0.1, 1, or 5 µM TAL at 14 DPA. Groups were analyzed using a one-way ANOVA using a Tukey-Kramer multiple comparison test. * = p < 0.05, ** = p < 0.01, *** = p < 0.001. (C) HCR-FISH for *Hoxa13* in DBs treated with 1 µM TAL at 14 DPA. Dashed line indicates amputation plane. Scale bars = 200 µm or 20 µm (inset). (D) HCR-FISH dot quantification in the epithelium and whole blastema for *Cyp26a1* and *Cyp26b1* in PBs and DBs at 10 and 14 DPA (n = 3-6, 3.5 cm (HT) animals aged 2.5 months). Axes and analyses as in Fig. 1E. * = p < 0.05.

**Figure S9:**
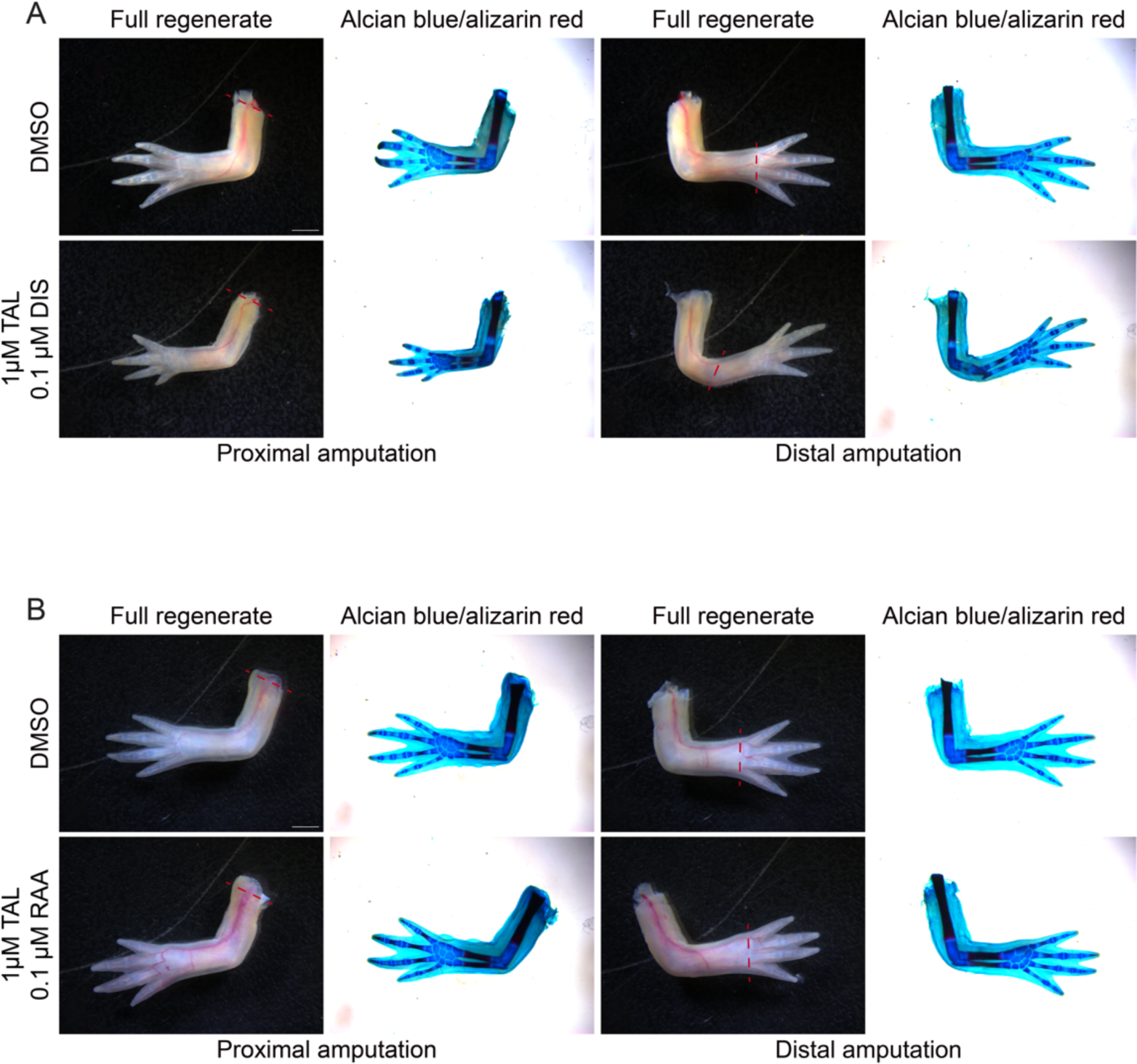
Cotreating TAL with DIS or RAA prevents proximalization. (A) Brightfield images of regenerates and skeletal structures of PBs and DBs treated with DMSO or 1µM TAL/0.1µM DIS. Dashed lines indicate amputation plane. Scale bar = 2 mm. (B) Brightfield images of regenerates and skeletal structures of PBs and DBs treated with DMSO or 1µM TAL/0.1µM RAA. Dashed lines indicate amputation plane. Scale bar = 2 mm.

**Figure S10:**
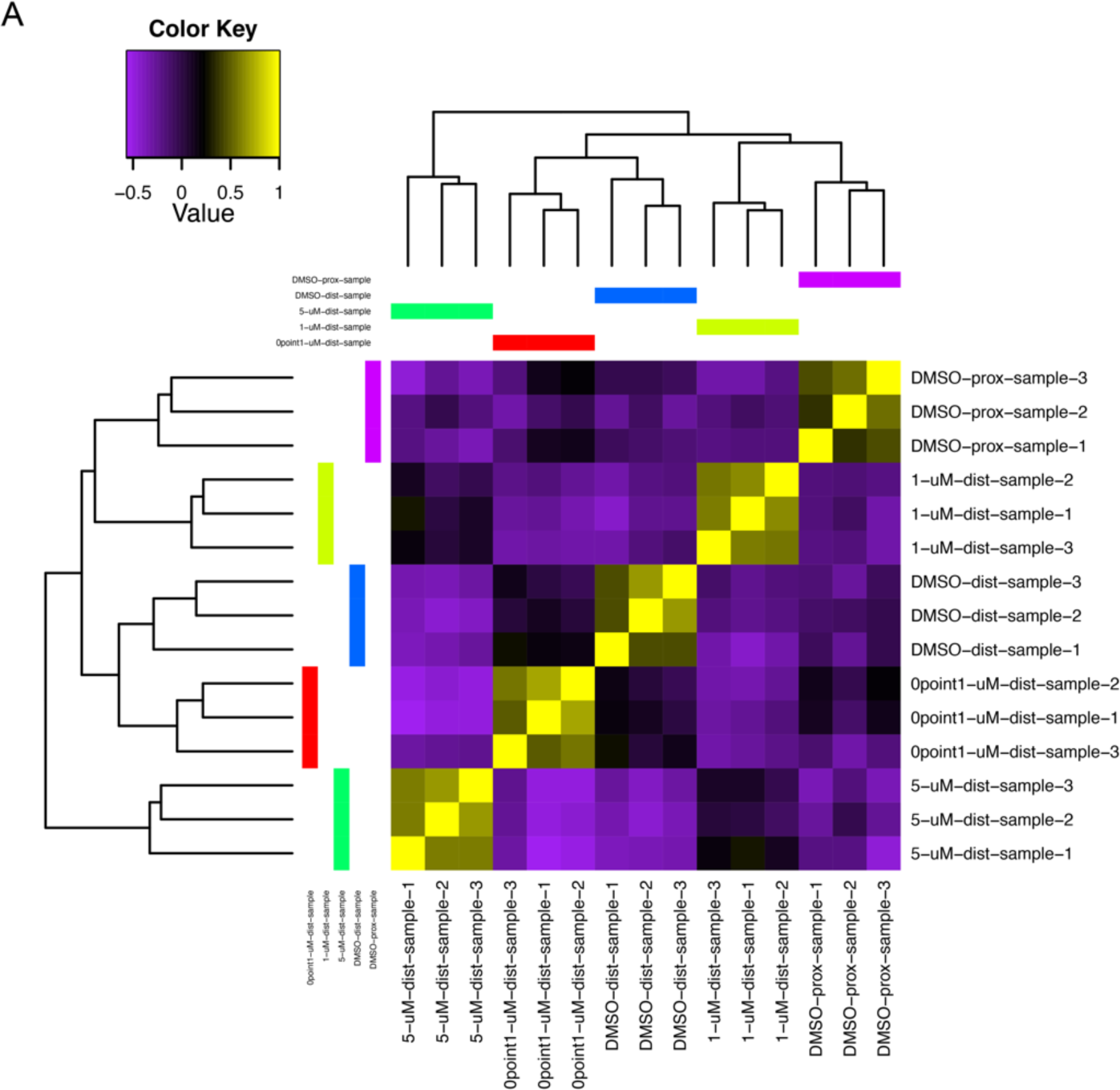
Additional results for bulk RNAseq results. (A) Sample correlation matrix showing relatedness between each sample

**Figure S11:**
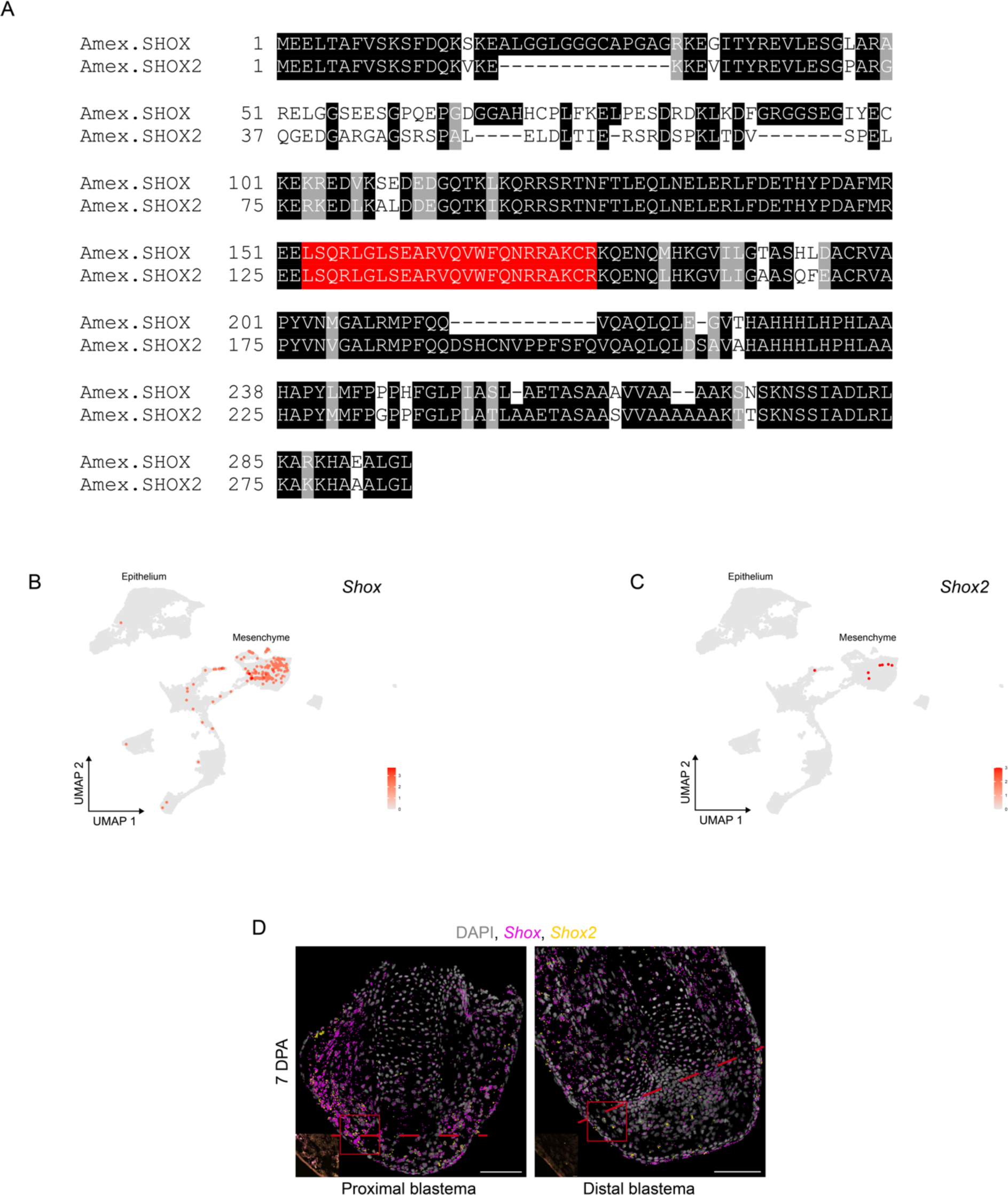
Additional characterization of *Shox* and *Shox2*. (A) Amino acid alignment of SHOX and SHOX2. Highlighted in red is the 100% conserved homeobox domain. (B) UMAP showing *Shox* in DBs at 7, 14, and 22 DPA. (C) UMAP showing *Shox2* in DBs at 7, 14, and 22 DPA. (D) HCR-FISH for *Shox* and *Shox2* in PBs and DBs at 7 DPA. Dashed lines indicate amputation plane. Scale bars = 200 µm or 20 µm (inset).

**Figure S12:**
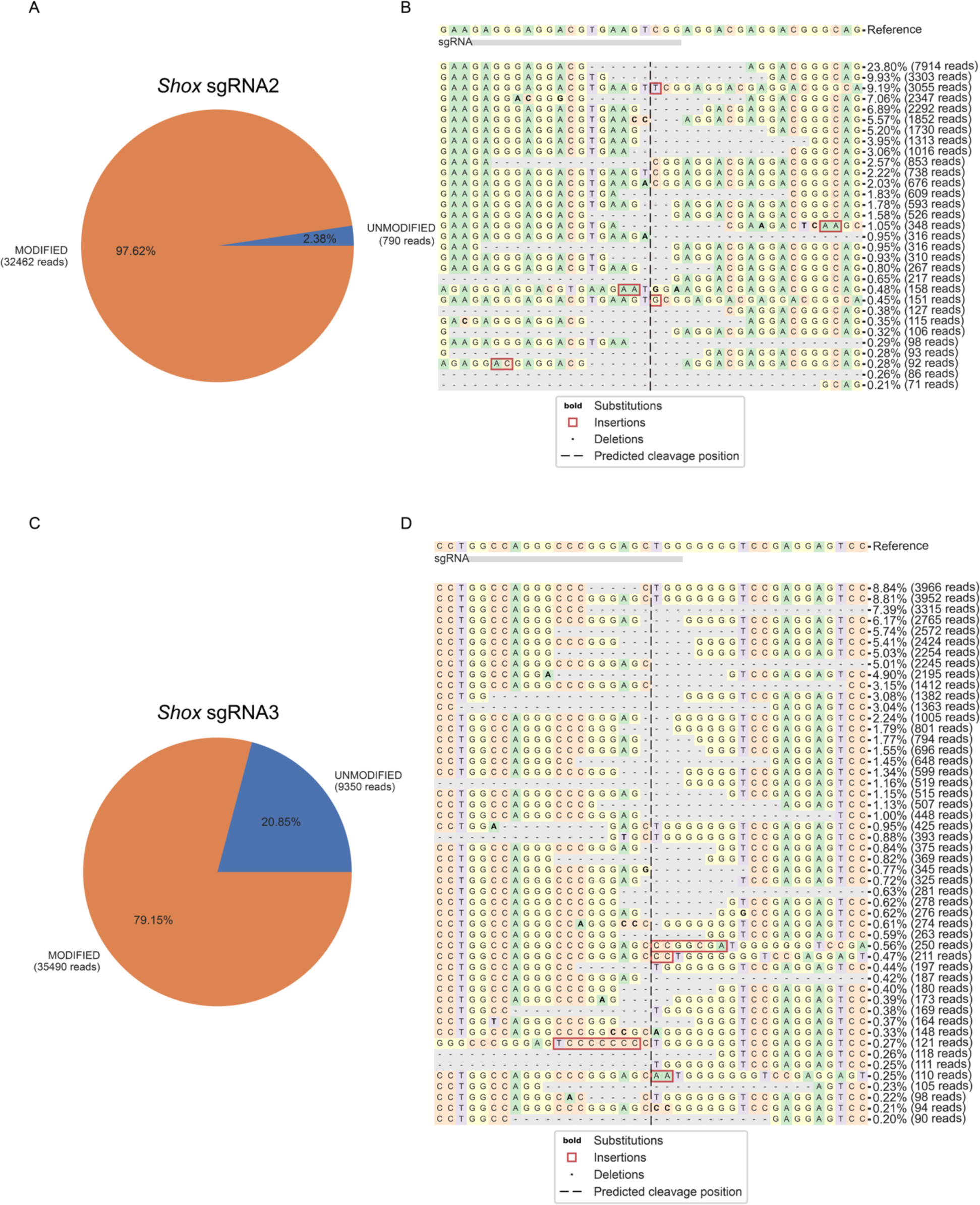
Genotyping for *Shox* crispants. (A) Pie chart for *Shox* sgRNA2 showing that 97.62% of alleles sequenced from 10 pooled tail tips were modified. (B) Sequence alignment for *Shox* sgRNA2 depicting the frequency of mutation type in each *Shox* crispants. (C) Pie chart for *Shox* sgRNA3 showing that 79.15% of alleles sequenced from 10 pooled tail tips were modified. (B) Sequence alignment for *Shox* sgRNA3 depicting the frequency of mutation type in each *Shox* crispants. Data generated from CRISPResso2 (Clement et al., 2019).

**Table S1: Primers used for qRT-PCR**

**Table S2: Probe sequences for HCR-FISH**

**Table S3: Quantification of PD limb duplications after TAL treatment**

**Table S4: Quantification of PD limb duplications after DIS or TAL/DIS treatment**

**Table S5: Quantification of PD limb duplications after RAA or TAL/RAA treatment**

**Table S6: Gene identities from each Venn diagram category in Figure 4D**

## Notes

### Competing Interest Statement

The authors have declared no competing interest.

## References

Bahry, E., Breimann, L., Zouinkhi, M., Epstein, L., Kolyvanov, K., Mamrak, N., King, B., Long, X., Harrington, K. I. S., Lionnet, T., et al. (2022). RS-FISH: precise, interactive, fast, and scalable FISH spot detection. Nature Methods 19, 1563–1567.

Bolger, A. M., Lohse, M. and Usadel, B. (2014). Trimmomatic: a flexible trimmer for Illumina sequence data. Bioinformatics 30, 2114–2120.

Bray, N. L., Pimentel, H., Melsted, P. and Pachter, L. (2016). Near-optimal probabilistic RNA-seq quantification. Nature Biotechnology 34, 525–527.

Brockes, J. P. (1992). Introduction of a retinoid reporter gene into the urodele limb blastema. Proc Natl Acad Sci U S A 89, 11386–11390.

Bryant, D. M., Johnson, K., DiTommaso, T., Tickle, T., Couger, M. B., Payzin-Dogru, D., Lee, T. J., Leigh, N. D., Kuo, T. H., Davis, F. G., et al. (2017). A Tissue-Mapped Axolotl De Novo Transcriptome Enables Identification of Limb Regeneration Factors. Cell Rep 18, 762–776.

Choudhary, S. and Satija, R. (2022). Comparison and evaluation of statistical error models for scRNA-seq. Genome Biology 23, 27.

Clement, K., Rees, H., Canver, M. C., Gehrke, J. M., Farouni, R., Hsu, J. Y., Cole, M. A., Liu, D. R., Joung, J. K., Bauer, D. E., et al. (2019). CRISPResso2 provides accurate and rapid genome editing sequence analysis. Nat Biotechnol 37, 224–226.

Cooper, K. L., Hu, J. K.-H., Berge, D. t., Fernandez-Teran, M., Ros, M. A. and Tabin, C. J. (2011). Initiation of Proximal-Distal Patterning in the Vertebrate Limb by Signals and Growth. Science 332, 1083-1086.

Crawford, K. and Stocum, D. L. (1988). Retinoic acid coordinately proximalizes regenerate pattern and blastema differential affinity in axolotl limbs. Development 102, 687–698.

Cuervo, R. and Chimal-Monroy, J. (2013). Chemical activation of RARβ induces post-embryonically bilateral limb duplication during Xenopus limb regeneration. Sci Rep 3, 1886.

Currie, J. D., Kawaguchi, A., Traspas, R. M., Schuez, M., Chara, O. and Tanaka, E. M. (2016). Live Imaging of Axolotl Digit Regeneration Reveals Spatiotemporal Choreography of Diverse Connective Tissue Progenitor Pools. Dev Cell 39, 411–423.

da Silva, S. M., Gates, P. B. and Brockes, J. P. (2002). The newt ortholog of CD59 is implicated in proximodistal identity during amphibian limb regeneration. Dev Cell 3, 547–555.

Datta, S. and Satten, G. A. (2005). Rank-Sum Tests for Clustered Data. Journal of the American Statistical Association 100, 908–915.

Delgado, I., López-Delgado, A. C., Roselló-Díez, A., Giovinazzo, G., Cadenas, V., Fernández-de-Manuel, L., Sánchez-Cabo, F., Anderson, M. J., Lewandoski, M. and Torres, M. (2020). Proximo-distal positional information encoded by an Fgf-regulated gradient of homeodomain transcription factors in the vertebrate limb. Science Advances 6, eaaz0742.

Dmetrichuk, J. M., Spencer, G. E. and Carlone, R. L. (2005). Retinoic acid-dependent attraction of adult spinal cord axons towards regenerating newt limb blastemas in vitro. Dev Biol 281, 112–120.

Fei, J. F., Lou, W. P., Knapp, D., Murawala, P., Gerber, T., Taniguchi, Y., Nowoshilow, S., Khattak, S. and Tanaka, E. M. (2018). Application and optimization of CRISPR-Cas9-mediated genome engineering in axolotl (Ambystoma mexicanum). Nat Protoc 13, 2908–2943.

Gardiner, D. M., Blumberg, B., Komine, Y. and Bryant, S. V. (1995). Regulation of HoxA expression in developing and regenerating axolotl limbs. Development 121, 1731–1741.

Gerber, T., Murawala, P., Knapp, D., Masselink, W., Schuez, M., Hermann, S., Gac-Santel, M., Nowoshilow, S., Kageyama, J., Khattak, S., et al. (2018). Single-cell analysis uncovers convergence of cell identities during axolotl limb regeneration. Science 362.

Grabherr, M. G., Haas, B. J., Yassour, M., Levin, J. Z., Thompson, D. A., Amit, I., Adiconis, X., Fan, L., Raychowdhury, R., Zeng, Q., et al. (2011). Full-length transcriptome assembly from RNA-Seq data without a reference genome. Nature Biotechnology 29, 644–652.

Gu, Z., Eils, R. and Schlesner, M. (2016). Complex heatmaps reveal patterns and correlations in multidimensional genomic data. Bioinformatics 32, 2847–2849.

Hao, Y., Stuart, T., Kowalski, M. H., Choudhary, S., Hoffman, P., Hartman, A., Srivastava, A., Molla, G., Madad, S., Fernandez-Granda, C., et al. (2024). Dictionary learning for integrative, multimodal and scalable single-cell analysis. Nat Biotechnol 42, 293–304.

Jiang, Y., Lee, M.-L. T., He, X., Rosner, B. and Yan, J. (2020). Wilcoxon Rank-Based Tests for Clustered Data with R Package clusrank. Journal of Statistical Software 96, 1–26.

Johnson, K. J. and Scadding, S. R. (1992). Effects of tunicamycin on retinoic acid induced respecification of positional values in regenerating limbs of the larval axolotl, Ambystoma mexicanum. Dev Dyn 193, 185–192.

Kawaguchi, A., Wang, J., Knapp, D., Murawala, P., Nowoshilow, S., Masselink, W., Taniguchi-Sugiura, Y., Fei, J. F. and Tanaka, E. M. (2024). A chromatin code for limb segment identity in axolotl limb regeneration. Dev Cell.

Kragl, M., Knapp, D., Nacu, E., Khattak, S., Maden, M., Epperlein, H. H. and Tanaka, E. M. (2009). Cells keep a memory of their tissue origin during axolotl limb regeneration. Nature 460, 60–65.

Lee, E., Ju, B.-G. and Kim, W.-S. (2012). Endogenous retinoic acid mediates the early events in salamander limb regeneration. Animal Cells and Systems 16, 462–468.

Li, H., Wei, X., Zhou, L., Zhang, W., Wang, C., Guo, Y., Li, D., Chen, J., Liu, T., Zhang, Y., et al. (2021). Dynamic cell transition and immune response landscapes of axolotl limb regeneration revealed by single-cell analysis. Protein Cell 12, 57–66.

Lin, T. Y., Gerber, T., Taniguchi-Sugiura, Y., Murawala, P., Hermann, S., Grosser, L., Shibata, E., Treutlein, B. and Tanaka, E. M. (2021). Fibroblast dedifferentiation as a determinant of successful regeneration. Dev Cell 56, 1541–1551.e1546.

Livak, K. J. and Schmittgen, T. D. (2001). Analysis of relative gene expression data using real-time quantitative PCR and the 2(-Delta Delta C(T)) Method. Methods 25, 402–408.

Love, M. I., Huber, W. and Anders, S. (2014). Moderated estimation of fold change and dispersion for RNA-seq data with DESeq2. Genome Biol 15, 550.

Lovely, A. M., Duerr, T. J., Stein, D. F., Mun, E. T. and Monaghan, J. R. (2023). Hybridization Chain Reaction Fluorescence In Situ Hybridization (HCR-FISH) in Ambystoma mexicanum Tissue. Methods Mol Biol 2562, 109–122.

Maden, M. (1982). Vitamin A and pattern formation in the regenerating limb. Nature 295, 672–675.

Maden, M. (1983). The effect of vitamin A on the regenerating axolotl limb. J Embryol Exp Morphol 77, 273–295.

Maden, M. (1998). Retinoids as endogenous components of the regenerating limb and tail. Wound Repair Regen 6, 358–365.

McGlinn, E., van Bueren, K. L., Fiorenza, S., Mo, R., Poh, A. M., Forrest, A., Soares, M. B., Bonaldo, M. d. F., Grimmond, S., Hui, C.-c., et al. (2005). Pax9 and Jagged1 act downstream of Gli3 in vertebrate limb development. Mechanisms of Development 122, 1218–1233.

Melsted, P., Ntranos, V. and Pachter, L. (2019). The barcode, UMI, set format and BUStools. Bioinformatics 35, 4472–4473.

Mercader, N., Leonardo, E., Piedra, M. E., Martinez, A. C., Ros, M. A. and Torres, M. (2000). Opposing RA and FGF signals control proximodistal vertebrate limb development through regulation of Meis genes. Development 127, 3961–3970.

Mercader, N., Selleri, L., Criado, L. M., Pallares, P., Parras, C., Cleary, M. and Torres, M. (2009). Ectopic *Meis1* expression in the mouse limb bud alters P-D patterning in a Pbx1-independent manner. Int. J. Dev. Biol. 53, 1483–1494.

Mercader, N., Tanaka, E. M. and Torres, M. (2005). Proximodistal identity during vertebrate limb regeneration is regulated by Meis homeodomain proteins. Development 132, 4131–4142.

Monaghan, J. R. and Maden, M. (2012). Visualization of retinoic acid signaling in transgenic axolotls during limb development and regeneration. Developmental biology 368, 63–75.

Nacu, E., Glausch, M., Le, H. Q., Damanik, F. F. R., Schuez, M., Knapp, D., Khattak, S., Richter, T. and Tanaka, E. M. (2013). Connective tissue cells, but not muscle cells, are involved in establishing the proximo-distal outcome of limb regeneration in the axolotl. Development 140, 513.

Nacu, E., Gromberg, E., Oliveira, C. R., Drechsel, D. and Tanaka, E. M. (2016). FGF8 and SHH substitute for anterior-posterior tissue interactions to induce limb regeneration. Nature 533, 407–410.

Nardi, J. B. and Stocum, D. L. (1984). Surface properties of regenerating limb cells: Evidence for gradation along the proximodistal axis. Differentiation 25, 27–31.

Nguyen, M., Singhal, P., Piet, J. W., Shefelbine, S. J., Maden, M., Voss, S. R. and Monaghan, J. R. (2017). Retinoic acid receptor regulation of epimorphic and homeostatic regeneration in the axolotl. Development 144, 601–611.

Niederreither, K., Subbarayan, V., Dolle, P. and Chambon, P. (1999). Embryonic retinoic acid synthesis is essential for early mouse post-implantation development. Nat Genet 21, 444–448.

Niederreither, K., Vermot, J., Schuhbaur, B., Chambon, P. and Dolle, P. (2002). Embryonic retinoic acid synthesis is required for forelimb growth and anteroposterior patterning in the mouse. Development 129, 3563–3574.

Niswander, L., Jeffrey, S., Martin, G. R. and Tickle, C. (1994). A positive feedback loop coordinates growth and patterning in the vertebrate limb. Nature 371, 609–612.

Nowoshilow, S., Schloissnig, S., Fei, J. F., Dahl, A., Pang, A. W. C., Pippel, M., Winkler, S., Hastie, A. R., Young, G., Roscito, J. G., et al. (2018). The axolotl genome and the evolution of key tissue formation regulators. Nature 554, 50–55.

Oliveira, C. R., Knapp, D., Elewa, A., Gerber, T., Gonzalez Malagon, S. G., Gates, P. B., Walters, H. E., Petzold, A., Arce, H., Cordoba, R. C., et al. (2022). Tig1 regulates proximo-distal identity during salamander limb regeneration. Nature Communications 13, 1141.

Pachitariu, M. and Stringer, C. (2022). Cellpose 2.0: how to train your own model. Nature Methods 19, 1634–1641.

Patro, R., Duggal, G., Love, M. I., Irizarry, R. A. and Kingsford, C. (2017). Salmon provides fast and bias-aware quantification of transcript expression. Nat Methods 14, 417–419.

Pescitelli, M. J., Jr. and Stocum, D. L. (1981). Nonsegmental organization of positional information in regenerating Ambystoma limbs. Dev Biol 82, 69–85.

Polvadore, T. and Maden, M. (2021). Retinoic Acid Receptors and the Control of Positional Information in the Regenerating Axolotl Limb. Cells 10.

Probst, S., Kraemer, C., Demougin, P., Sheth, R., Martin, G. R., Shiratori, H., Hamada, H., Iber, D., Zeller, R. and Zuniga, A. (2011). SHH propagates distal limb bud development by enhancing CYP26B1-mediated retinoic acid clearance via AER-FGF signalling. Development 138, 1913–1923.

Rao, E., Weiss, B., Fukami, M., Rump, A., Niesler, B., Mertz, A., Muroya, K., Binder, G., Kirsch, S., Winkelmann, M., et al. (1997). Pseudoautosomal deletions encompassing a novel homeobox gene cause growth failure in idiopathic short stature and Turner syndrome. Nat Genet 16, 54–63.

Riquelme-Guzmán, C. and Sandoval-Guzmán, T. (2023). Methods for Studying Appendicular Skeletal Biology in Axolotls. Methods Mol Biol 2562, 155–163.

Roensch, K., Tazaki, A., Chara, O. and Tanaka, E. M. (2013). Progressive specification rather than intercalation of segments during limb regeneration. Science 342, 1375–1379.

Roselló-Díez, A., Arques, C. G., Delgado, I., Giovinazzo, G. and Torres, M. (2014). Diffusible signals and epigenetic timing cooperate in late proximo-distal limb patterning. Development 141, 1534–1543.

Roselló-Díez, A., Ros, M. A. and Torres, M. (2011). Diffusible Signals, Not Autonomous Mechanisms, Determine the Main Proximodistal Limb Subdivision. Science 332, 1086–1088.

Saiz-Lopez, P., Chinnaiya, K., Campa, V. M., Delgado, I., Ros, M. A. and Towers, M. (2015). An intrinsic timer specifies distal structures of the vertebrate limb. Nature Communications 6, 8108.

Scadding, S. R. (1999). Citral, an inhibitor of retinoic acid synthesis, modifies pattern formation during limb regeneration in the axolotl Ambystoma mexicanum. Canadian Journal of Zoology 77, 1835–1837.

Scadding, S. R. and Maden, M. (1986). Comparison of the effects of vitamin A on limb development and regeneration in the axolotl, Ambystoma mexicanum. J Embryol Exp Morphol 91, 19–34.

Scadding, S. R. and Maden, M. (1994). Retinoic Acid Gradients during Limb Regeneration. Developmental Biology 162, 608–617.

Schindelin, J., Arganda-Carreras, I., Frise, E., Kaynig, V., Longair, M., Pietzsch, T., Preibisch, S., Rueden, C., Saalfeld, S., Schmid, B., et al. (2012). Fiji: an open-source platform for biological-image analysis. Nature Methods 9, 676-682.

Schloissnig, S., Kawaguchi, A., Nowoshilow, S., Falcon, F., Otsuki, L., Tardivo, P., Timoshevskaya, N., Keinath, M. C., Smith, J. J., Voss, S. R., et al. (2021). The giant axolotl genome uncovers the evolution, scaling, and transcriptional control of complex gene loci. Proceedings of the National Academy of Sciences 118, e2017176118.

Seifert, A. W., Monaghan, J. R., Voss, S. R. and Maden, M. (2012). Skin regeneration in adult axolotls: a blueprint for scar-free healing in vertebrates. PLoS One 7, e32875.

Shears, D. J., Vassal, H. J., Goodman, F. R., Palmer, R. W., Reardon, W., Superti-Furga, A., Scambler, P. J. and Winter, R. M. (1998). Mutation and deletion of the pseudoautosomal gene SHOX cause Leri-Weill dyschondrosteosis. Nat Genet 19, 70–73.

Stringer, C., Wang, T., Michaelos, M. and Pachitariu, M. (2021). Cellpose: a generalist algorithm for cellular segmentation. Nature Methods 18, 100–106.

Takeuchi, T., Matsubara, H., Minamitani, F., Satoh, Y., Tozawa, S., Moriyama, T., Maruyama, K., Suzuki, K. T., Shigenobu, S., Inoue, T., et al. (2022). Newt Hoxa13 has an essential and predominant role in digit formation during development and regeneration. Development 149.

te Welscher, P., Fernandez-Teran, M., Ros, M. A. and Zeller, R. (2002). Mutual genetic antagonism involving GLI3 and dHAND prepatterns the vertebrate limb bud mesenchyme prior to SHH signaling. Genes Dev 16, 421–426.

Thoms, S. D. and Stocum, D. L. (1984). Retinoic acid-induced pattern duplication in regenerating urodele limbs. Dev Biol 103, 319–328.

Trofka, A., Huang, B. L., Zhu, J., Heinz, W. F., Magidson, V., Shibata, Y., Shi, Y. B., Tarchini, B., Stadler, H. S., Kabangu, M., et al. (2021). Genetic basis for an evolutionary shift from ancestral preaxial to postaxial limb polarity in non-urodele vertebrates. Curr Biol 31, 4923–4934.e4925.

Uzkudun, M., Marcon, L. and Sharpe, J. (2015). Data-driven modelling of a gene regulatory network for cell fate decisions in the growing limb bud. Mol Syst Biol 11, 815.

Vieira, W. A. and McCusker, C. D. (2019). Hierarchical pattern formation during amphibian limb regeneration. Biosystems 183, 103989.

Viviano, C. M., Horton, C. E., Maden, M. and Brockes, J. P. (1995). Synthesis and release of 9-cis retinoic acid by the urodele wound epidermis. Development 121, 3753–3762.

Wickham, H. (2016). ggplot2: Elegant Graphics for Data Analysis: Springer International Publishing.

Wolpert, L. (1969). Positional information and the spatial pattern of cellular differentiation. Journal of Theoretical Biology 25, 1–47.

Yashiro, K., Zhao, X., Uehara, M., Yamashita, K., Nishijima, M., Nishino, J., Saijoh, Y., Sakai, Y. and Hamada, H. (2004). Regulation of retinoic acid distribution is required for proximodistal patterning and outgrowth of the developing mouse limb. Dev Cell 6, 411–422.

Yu, L., Liu, H., Yan, M., Yang, J., Long, F., Muneoka, K. and Chen, Y. (2007). Shox2 is required for chondrocyte proliferation and maturation in proximal limb skeleton. Developmental Biology 306, 549–559.

Zasada, M. and Budzisz, E. (2019). Retinoids: active molecules influencing skin structure formation in cosmetic and dermatological treatments. Postepy Dermatol Alergol 36, 392–397.

