## Supplementary material for "Retinoic acid breakdown is required for proximodistal positional identity during amphibian limb regeneration": Table S6

| **DEGs shared between DMSO DBs vs DMSO PBs and DMSO DBs vs 1μM TAL DBs** | **DEGs unique to DMSO DBs vs 1μM TAL DBs** | **DEGs unique to DMSO DBs vs DMSO PBs** |
| --- | --- | --- |
| SHOX2 | CLDN5 | ANKLE2 |
| AMEX60DDU001035434 | MMP11 | LOC107385925 |
| SHOX | CCDC74A | CCDC92 |
| CHRM4 | ALDH2 | HPD |
| TBX15 | SLC2A9 | AMEX60DD000664 |
| AMEX60DD005167 | OASL | USP30 |
| AMEX60DDU001013699 | LOC115073459 | AMEX60DD000878 |
| AMEX60DD003537 | EMID1 | DR999_PMT12787 |
| LOC114593837 | AMEX60DD001052 | AMEX60DD001061 |
| ALX4 | PEBP1 | XRN1 |
| PARP4 | IL1B | PAK2 |
| AMEX60DDU001017832 | LOC112551351 | COPS9 |
| AMEX60DDU001026790 | CYP2J2 | SST |
| ZBTB16 | LOC115082225 | AMEX60DD003455 |
| ENC1 | B3GNT5 | CLTC |
| LEP | VWA5A | MEX3B |
| AMEX60DDU001012288 | AMEX60DD001987 | XELAEV_18038498MG |
| PRRX1 | ROHU_001730 | AMEX60DD004769 |
| KRT19 | EPHA4 | TMEM41B |
| ADGRL2 | HDLBP | GAS2 |
| POL | AMEX60DD002380 | AMEX60DD005171 |
| TGFB2 | PLSCR1 | AMEX60DD005415 |
| CRABP2 | HARBI1 | FCN1 |
| AMEX60DD009711 | ZNF703 | AMEX60DD005894 |
| AMEX60DD046442 | RBP1 | N/A |
| AMEX60DD052103 | LOC114092512 | PHYH |
| CYP2J2 | AMEX60DD002997 | LDHB |
| ABTB2 | AMEX60DD003027 | CCND2 |
| AMEX60DD029540 | AMEX60DD003031 | FAM180A |
| AMEX60DD048926 | PTPRH | AMEX60DD007717 |
| AMEX60DD018451 | LOC108718065 | AMEX60DD008223 |
| APELA | RTP3 | ITPR3 |
| CXCL14 | CLDN1 | PIM1 |
| AMEX60DDU001010337 | OSTN | TGM1 |
| HOXA13 | CIB1 | COL1A1 |
| AMEX60DDU001037429 | IGDCC3 | AMEX60DD010032 |
| KRT17 | CILP | KRT12 |
| KII | ALDH1A3 | TRIM39 |
| LOC108696921 | TCF12 | AMEX60DD010547 |
| COCH | THSD4 | LOC115082302 |
| EPHA3 | SUGT1P4-STRA6LP | AMEX60DD010651 |
| AMEX60DDU001029479 | LOC115075475 | TAP1 |
| AMEX60DDU001036305 | LOC115075287 | AMEX60DD010885 |
| AMEX60DD051976 | RHCG | AMEX60DD010887 |
| AMEX60DDU001018078 | ITGA11 | LOC107986004 |
| EVX1 | N/A | AMEX60DD011241 |
| VSTM2A | PHLDA2 | AMEX60DD011243 |
| HOXD13 | LOC101949100 | SIX1 |
| AMEX60DDU001004023 | AMEX60DD004344 | RDH12 |
| AMEX60DDU001017189 | IRF7 | AMEX60DD012311 |
| TENM4 | MUC5AC | AMEX60DD013447 |
| TNN | AMEX60DD004714 | B3GNT3 |
| AMEX60DDU001025455 | LMO1 | AMEX60DD014023 |
| FLRT3 | PAX6 | MYO1F |
| AMEX60DDU001028376 | FSH | AMEX60DD014706 |
| AMEX60DDU001036575 | ELF5 | AMEX60DD015653 |
| AMEX60DDU001023145 | PAMR1 | KIRREL1 |
| AMEX60DD017703 | CD82 | NTN1 |
| AMEX60DDU001004497 | LOC115468841 | AMEX60DD017008 |
| AMEX60DDU001039043 | SMPDL3B | AMEX60DD017446 |
| LOC115098183 | UMOD | AMEX60DD017457 |
| AMEX60DDU001029836 | AMEX60DD005691 | AMEX60DD017687 |
| AMEX60DDU001023767 | GP2 | AMEX60DD017969 |
| NDNF | AMEX60DD005785 | PLAUR |
| AMEX60DD021955 | PNKD | VWA7 |
| AMEX60DD032374 | COL9A2 | FMO3 |
| AMEX60DD012676 | FUCA1 | RRBP1 |
| AMEX60DDU001003360 | HGF | AMEX60DD018915 |
| AMEX60DDU001028515 | ELAPOR2 | CCN1 |
| AMEX60DDU001040316 | UPK3A | JUN |
| AMEX60DDU001038586 | NYNRIN | AMEX60DD019701 |
| EMP1 | USP18 | AMEX60DD019911 |
| GSONMT00055939001 | LOC115478063 | AMEX60DD020038 |
| LOC115085683 | TIMP3 | LOC102946426 |
| AMEX60DDU001035919 | ACAN | UMOD |
| AMEX60DDU001011650 | MAS1 | AMEX60DD021009 |
| LOC112543511 | CNTF | HBG1 |
| AMEX60DDU001037422 | TMEM176B | HB-AM |
| KRT6A | RHOF | HBZ |
| AMEX60DD054010 | CLCF1 | CPVL |
| AMEX60DDU001011461 | GSTP1 | CCK |
| GSONMT00075035001 | LOC102567599 | AMEX60DD022241 |
| AMEX60DDU001039088 | SLC38A2 | MEOX2 |
| MSLN | HSP90B1 | AMEX60DD022790 |
| EVX2 | LOC115073812 | TMLHE |
| PAX9 | LOC115481879 | FGL2 |
| AMEX60DD027120 | SCUBE3 | AMEX60DD024041 |
| LOC105079761 | BCL2L15 | CHGA |
| TIMP3 | PPM1H | OTX2 |
| NR2F2 | MOV10 | AMEX60DD024621 |
| ENV | SMYD3 | AMEX60DD024789 |
| LGALS3BP | FMOD | ZNF208 |
| DSG4 | GSTM1 | AMEX60DD024977 |
| LPO | AMEX60DD009552 | LRRN4CL |
| GSONMT00022338001 | IRF8 | AMEX60DD025503 |
| AMEX60DDU001031440 | MYO1D | ALAS2 |
| WFDC8 | NPY | AMEX60DD026005 |
| AMEX60DDU001039758 | SOST | HBD |
| KRT12 | ARL4D | GLNC1 |
| AMEX60DDU001022902 | DHX58 | PARPI_0020006 |
| AMEX60DDU001026793 | JUP | C1S |
| LOC115078387 | KRT15 | AMEX60DD027084 |
| AMEX60DD024721 | KRT12 | AMEX60DD027603 |
| LOC115074559 | TNS4 | IL17B |
| TFAP2B | TOP2A | HBG2 |
| ROHU_035623 | HOXB13 | HBE1 |
| GJB6 | MIEN1 | AMEX60DD028126 |
| AMEX60DDU001037682 | TRIM39-RPP21 | AMEX60DD028209 |
| LOC112977635 | AMEX60DD010432 | GDF5 |
| ENDOU | TRIM39 | DDX46 |
| UMOD | PSMB7 | HOXC4 |
| PLK2 | XB22063241[PROVISIONAL:LY6A2] | KRT5 |
| AMEX60DD027541 | AMEX60DD010633 | PRPH |
| AMEX60DD050350 | LOC115085991 | HAND2 |
| AMEX60DD016059 | DDAH2 | AMEX60DD031109 |
| CRYM.L | DXO | AMEX60DD031141 |
| LOC112545447 | HLA-DPB1 | APOH |
| TRH | LOC112123664 | RNF213 |
| SEMA3F | LBH | AMEX60DD032141 |
| AQP3 | PKDCC | RHAG |
| A2M | SLC25A29 | MYCN |
| LOC115476892 | AMEX60DD011527 | LOC114552212 |
| VWA1 | ZFP36L1 | TBX18 |
| TRIM15 | STXBP6 | AMEX60DD034235 |
| GAD1 | AMEX60DD012267 | CCN2 |
| COL2A1 | AB205_0062080 | AMEX60DD034356 |
| LOC111565595 | ATP1B2 | AMEX60DD035221 |
| AMEX60DD032872 | AMEX60DD012939 | AMEX60DD035740 |
| AMEX60DD039602 | AMEX60DD013106 | MRPS10 |
| CST14B.1 | AMEX60DD013530 | KCNK5 |
| AMEX60DDU001018245 | TM6SF2 | GNMT |
| DUOX1 | STAP2 | AMEX60DD037245 |
| DR999_PMT21178 | GRAMD2B | DACH2 |
| SLITRK6 | AMEX60DD014562 | AMEX60DD038299 |
| NAMPT.S | ANXA9 | FOXC1 |
| AMEX60DD010634 | AMEX60DD014765 | NDUFB5 |
| CRLF1 | IFIH1 | CCDC102B |
|  | LOC115458134 | RNF125 |
|  | THBS3 | MSC |
|  | LOC115077151 | LOC115089817 |
|  | RERGL | HEY1 |
|  | LOC112547415 | VMD2 |
|  | AMEX60DD015399 | AMEX60DD040981 |
|  | XAF1 | LOC115481447 |
|  | LOC108433851 | AMEX60DD041114 |
|  | AMEX60DD015940 | LOC114069761 |
|  | AMEX60DD015941 | FCHO2 |
|  | AMEX60DD015942 | AMEX60DD042814 |
|  | D9C73_028124 | MAMDC2 |
|  | FCGBP | VPS13A |
|  | B4GALT3 | AMEX60DD043648 |
|  | CLEC11A | AMEX60DD043685 |
|  | IRF3 | LOC108706699 |
|  | PRMT1 | MAB21L2 |
|  | AMEX60DD016376 | AMEX60DD045170 |
|  | LOC115080300 | AMEX60DD045299 |
|  | LOC108697796 | LOC101935553 |
|  | IL11 | LOC112059317 |
|  | RHBG | CYP26B1 |
|  | AMEX60DD017007 | ZNF300 |
|  | CA5B | SPAG17 |
|  | WFDC1 | TSPAN7 |
|  | SOCS3 | AMEX60DD047878 |
|  | ATF5 | AMEX60DD047904 |
|  | EOD39_21475 | ST6GAL2 |
|  | AMEX60DD017702 | LOC109141055 |
|  | CMKLR1 | DCLK1 |
|  | AMEX60DD017947 | MAB21L1 |
|  | AMEX60DD018092 | AMEX60DD049414 |
|  | OLFML2B | AMEX60DD050166 |
|  | PRRX1 | ASS1 |
|  | AMEX60DD018606 | AMEX60DD050446 |
|  | RNASEL | AMEX60DD050994 |
|  | AMEX60DD018687 | MXRA8 |
|  | LHX9 | ALPL |
|  | CLCA2 | AMEX60DD051774 |
|  | JAK1 | LOC102366391 |
|  | ROR1 | AMEX60DD052686 |
|  | CYP4B1 | AMEX60DD052922 |
|  | KLF4 | TCF7L2 |
|  | LOC115183147 | AMEX60DD053231 |
|  | NTN1 | AMEX60DD053324 |
|  | TMC7 | APLP2 |
|  | IL4R | LOC115073865 |
|  | FAM20C | TTC36 |
|  | LOC112543511 | TMPRSS13 |
|  | MSLN | AMEX60DD054004 |
|  | AMEX60DD021073 | AMEX60DD054465 |
|  | PARPI_0026573 | NOS2 |
|  | AMEX60DD021717 | UBC |
|  | AMEX60DD021718 | LOC115081983 |
|  | GSDME | HOXD10 |
|  | AMEX60DD021925 | AMEX60DD056123 |
|  | HOXA3 | NNMT |
|  | AMEX60DD022304 | LOC105357518 |
|  | AGR2 | LOC112545447 |
|  | CITED1 | AMEX60DDU001000438 |
|  | AMEX60DD022620 | LOC105379356 |
|  | AMEX60DD022623 | AMEX60DDU001000909 |
|  | TRIM71 | AMEX60DDU001000989 |
|  | HCG_2039481 | AMEX60DDU001001010 |
|  | LOC102564405 | ALKBH3.L |
|  | SEC61A1 | AMEX60DDU001001321 |
|  | BHLHE40 | AMEX60DDU001002018 |
|  | WNT5A | AMEX60DDU001002251 |
|  | AMEX60DD023735 | AMEX60DDU001002259 |
|  | TREX2 | AMEX60DDU001002319 |
|  | PPARG | AMEX60DDU001002383 |
|  | INKA1 | LOC102361187 |
|  | AMEX60DD024219 | AMEX60DDU001002719 |
|  | UBA1 | AMEX60DDU001002720 |
|  | ACTN4 | EAF1 |
|  | NCCRP1 | AMEX60DDU001003433 |
|  | LOC115641468 | AMEX60DDU001003497 |
|  | RNASET2 | AMEX60DDU001003810 |
|  | AMEX60DD024642 | E1301_TTI023775 |
|  | AMEX60DD024693 | PARPI_0025344 |
|  | LOC106737871 | AMEX60DDU001004238 |
|  | TGFB2 | AMEX60DDU001004278 |
|  | PLD3 | AMEX60DDU001004291 |
|  | PFKFB1 | AMEX60DDU001004633 |
|  | SSR4 | AMEX60DDU001005147 |
|  | NT5DC2 | AMEX60DDU001005594 |
|  | FLNA | AMEX60DDU001005702 |
|  | PLXNA3 | AMEX60DDU001005842 |
|  | LITAF | AMEX60DDU001006151 |
|  | PPL | AMEX60DDU001006284 |
|  | LOC102473117 | AMEX60DDU001006372 |
|  | LOC115077990 | LOC112544220 |
|  | AMEX60DD026386 | AMEX60DDU001006790 |
|  | AMEX60DD026427 | MARCKS |
|  | AMEX60DD026447 | LOC115088534 |
|  | LOC103261025 | AMEX60DDU001007159 |
|  | AMEX60DD026561 | AMEX60DDU001007283 |
|  | LOC102568417 | AMEX60DDU001007451 |
|  | YBX3 | AMEX60DDU001007499 |
|  | COL9A3 | AMEX60DDU001007551 |
|  | OGFR | AMEX60DDU001007600 |
|  | HELZ2 | AMEX60DDU001007633 |
|  | BMP7 | AMEX60DDU001007769 |
|  | CYP24A1 | LOC115481199 |
|  | SALL4 | AMEX60DDU001008246 |
|  | ZNFX1 | AMEX60DDU001008316 |
|  | PLTP | AMEX60DDU001008317 |
|  | TGM6 | AMEX60DDU001008532 |
|  | LOC114842160 | AMEX60DDU001008796 |
|  | AMEX60DD027662 | AMEX60DDU001009352 |
|  | ELOB | AMEX60DDU001009716 |
|  | LOC115094691 | HARBI1 |
|  | PRSS27 | AMEX60DDU001010124 |
|  | AMEX60DD028191 | AMEX60DDU001010266 |
|  | TMPRSS5 | AMEX60DDU001010341 |
|  | CRK | AMEX60DDU001010485 |
|  | AMEX60DD028455 | WDR3 |
|  | MATN4.S | PARPI_0022647 |
|  | AMEX60DD028734 | AMEX60DDU001011897 |
|  | SPRY4 | AMEX60DDU001011982 |
|  | AMEX60DD028869 | AMEX60DDU001012072 |
|  | TMEM238 | AMEX60DDU001012460 |
|  | AMEX60DD029259 | AMEX60DDU001012529 |
|  | RAC1 | AMEX60DDU001012569 |
|  | HOXC13 | AMEX60DDU001012833 |
|  | RARG | AMEX60DDU001012975 |
|  | CYP27B1 | LOC106733090 |
|  | KRT5 | LOC112547415 |
|  | OJAV_G00236990 | AMEX60DDU001013614 |
|  | AMEX60DD030332 | AMEX60DDU001013820 |
|  | GPRC5A | AMEX60DDU001014129 |
|  | AMEX60DD030636 | AMEX60DDU001014181 |
|  | PMP22 | AMEX60DDU001014398 |
|  | KIAA1618 | AMEX60DDU001014523 |
|  | F11 | AMEX60DDU001014634 |
|  | JUNB | N300_12895 |
|  | PRDX2 | AMEX60DDU001015402 |
|  | CALR | AMEX60DDU001015455 |
|  | AMEX60DD031992 | AMEX60DDU001015843 |
|  | LOC110166736 | AMEX60DDU001016534 |
|  | UCKL1 | AMEX60DDU001017112 |
|  | AMEX60DD032597 | AMEX60DDU001017373 |
|  | LOC108804010 | AMEX60DDU001017448 |
|  | AMEX60DD033140 | AMEX60DDU001017460 |
|  | AMEX60DD033423 | AMEX60DDU001017751 |
|  | AMEX60DD033498 | AMEX60DDU001017898 |
|  | COL9A1 | AMEX60DDU001018587 |
|  | PRDM1 | AMEX60DDU001018605 |
|  | LOC102371746 | AMEX60DDU001018776 |
|  | AMEX60DD034207 | AMEX60DDU001018784 |
|  | APMAP | AMEX60DDU001018907 |
|  | DLL1 | AMEX60DDU001018998 |
|  | RSPO3 | LOC102461666 |
|  | NCOA7 | AMEX60DDU001019384 |
|  | AMEX60DD035121 | LOC107982656 |
|  | SRD5A2 | AMEX60DDU001019802 |
|  | LTBP1 | AMEX60DDU001020421 |
|  | MTR | OGDH |
|  | AMEX60DD035546 | AMEX60DDU001020680 |
|  | JAG1 | AMEX60DDU001020989 |
|  | SPTLC3 | AMEX60DDU001021301 |
|  | EPAS1 | AMEX60DDU001021623 |
|  | AMEX60DD035959 | AMEX60DDU001021749 |
|  | AMEX60DD036840 | AMEX60DDU001021769 |
|  | P2RY4 | AMEX60DDU001021807 |
|  | AMEX60DD037000 | AMEX60DDU001022240 |
|  | DUI87_25150 | AMEX60DDU001022445 |
|  | ZIC3 | AMEX60DDU001023118 |
|  | IL13RA1 | AMEX60DDU001023235 |
|  | TNMD | E1301_TTI006344 |
|  | DSP | AMEX60DDU001023577 |
|  | AMEX60DD038389 | AMEX60DDU001023953 |
|  | SERPINB2 | AMEX60DDU001024489 |
|  | AMEX60DD038466 | AMEX60DDU001024606 |
|  | TPMT | LOC112543511 |
|  | SERPINB5 | AMEX60DDU001025722 |
|  | HSBP1L1 | AMEX60DDU001025835 |
|  | SLC66A2 | AMEX60DDU001025977 |
|  | AMEX60DD039227 | AMEX60DDU001026207 |
|  | SATL1 | AMEX60DDU001026537 |
|  | E1301_TTI013099 | AMEX60DDU001026694 |
|  | SULF1 | AMEX60DDU001026728 |
|  | YWHAZ | AMEX60DDU001026828 |
|  | PKHD1L1 | AMEX60DDU001026933 |
|  | AMEX60DD040339 | AMEX60DDU001027642 |
|  | TRPS1 | AMEX60DDU001028088 |
|  | CCN3 | AMEX60DDU001028320 |
|  | FAM83A | AMEX60DDU001028341 |
|  | AMEX60DD040550 | AMEX60DDU001028831 |
|  | AMEX60DD040575 | AMEX60DDU001029159 |
|  | AMEX60DD040656 | AMEX60DDU001029485 |
|  | AMEX60DD040756 | AMEX60DDU001029529 |
|  | DST | AMEX60DDU001029531 |
|  | SCX | AMEX60DDU001030638 |
|  | RPS13 | AMEX60DDU001030674 |
|  | LOC114789697 | AMEX60DDU001030709 |
|  | AMEX60DD041013 | AMEX60DDU001030716 |
|  | LOC116061033 | AMEX60DDU001030879 |
|  | AMEX60DD041062 | AMEX60DDU001030988 |
|  | LOC115480847 | AMEX60DDU001031353 |
|  | LOC115572840 | PARPI_0014898 |
|  | LOC112551687 | AMEX60DDU001031829 |
|  | AMEX60DD041183 | AMEX60DDU001032048 |
|  | LOC115073646 | Y1Q_0014614 |
|  | LOC115079342 | AMEX60DDU001032541 |
|  | TRIM27 | LOC115095888 |
|  | SLC14A2 | AMEX60DDU001032971 |
|  | ROHU_001488 | AMEX60DDU001033176 |
|  | AMEX60DD042484 | AMEX60DDU001033447 |
|  | LOC114595149 | AMEX60DDU001033522 |
|  | AMEX60DD042658 | AMEX60DDU001033785 |
|  | CDO1 | PRRC2A |
|  | NFIL3 | AMEX60DDU001034247 |
|  | AOPEP | AMEX60DDU001034289 |
|  | GAS1 | AMEX60DDU001034719 |
|  | KLF9 | AMEX60DDU001035063 |
|  | TMEM215 | AMEX60DDU001035284 |
|  | AMEX60DD043448 | AMEX60DDU001035416 |
|  | AMEX60DD043517 | AMEX60DDU001035657 |
|  | AMEX60DD043525 | LOC106705246 |
|  | LOC115080730 | AMEX60DDU001036731 |
|  | CORO2A | AMEX60DDU001036769 |
|  | AMEX60DD043826 | LOC115460730 |
|  | STAP1 | AMEX60DDU001036829 |
|  | LOC115089273 | AMEX60DDU001036940 |
|  | FABP2 | AMEX60DDU001037306 |
|  | ELOVL6 | AMEX60DDU001037542 |
|  | AMEX60DD044318 | AMEX60DDU001037591 |
|  | LOC115478516 | AMEX60DDU001038105 |
|  | DMRTA1 | AMEX60DDU001038354 |
|  | CASP6 | AMEX60DDU001038476 |
|  | DR999_PMT00179 | AMEX60DDU001038725 |
|  | AMEX60DD044971 | AMEX60DDU001039046 |
|  | MAB21L2 | AMEX60DDU001039294 |
|  | TLR2 | AMEX60DDU001039320 |
|  | HAND2 | AMEX60DDU001039922 |
|  | CRMP1 | AMEX60DDU001040550 |
|  | KAZALD1 | AMEX60DDU001040833 |
|  | AMEX60DD046395 | AMEX60DDU001040883 |
|  | CPXM1 | AMEX60DDU001040910 |
|  | CPXM2 | AMEX60DDU001041233 |
|  | LOC115087755 | AMEX60DDU001041464 |
|  | TTC31 | AMEX60DDU001041593 |
|  | PRDM9 | POL |
|  | D9C73_028471 | TMEM161B |
|  | ATP1A1 | AMEX60DDU001042089 |
|  | AMEX60DD046915 | LOC113050139 |
|  | AMEX60DD047120 | |
|  | RIPPLY3 |  |
|  | RUNX1 |  |
|  | AMEX60DD047256 | |
|  | AMEX60DD047683 | |
|  | CRACDL |  |
|  | ENV |  |
|  | ZIC2 |  |
|  | ZIC5 |  |
|  | EGFL6 |  |
|  | LOC113406022 | |
|  | AMEX60DD048744 | |
|  | FBXL3 |  |
|  | AMEX60DD049360 | |
|  | MMP13 |  |
|  | YAP1 |  |
|  | PRSS23 |  |
|  | TCAF2 |  |
|  | PTGES |  |
|  | RALGDS |  |
|  | HSPA5 |  |
|  | NR6A1 |  |
|  | LOC115088534 | |
|  | LOC115097977 | |
|  | LOC104149496 | |
|  | LOC112432775 | |
|  | BRINP1 |  |
|  | GLUL |  |
|  | LOC107728464 | |
|  | PTGS1 |  |
|  | ENO1 |  |
|  | UNC5B |  |
|  | AMEX60DD051967 | |
|  | AMEX60DD051969 | |
|  | AMEX60DD051970 | |
|  | AMEX60DD051971 | |
|  | AMEX60DD051972 | |
|  | AMEX60DD051980 | |
|  | GLUD1 |  |
|  | CAPNS1 |  |
|  | DHRS3 |  |
|  | LOC102939808 | |
|  | AMEX60DD052390 | |
|  | ARHGAP22 |  |
|  | IFIT5 |  |
|  | CYP26A1 |  |
|  | AMEX60DD053035 | |
|  | CASP7 |  |
|  | PLEKHS1 |  |
|  | DMBT1 |  |
|  | AMEX60DD053453 | |
|  | ATP12A |  |
|  | LOC113036195 | |
|  | BARX2 |  |
|  | LOC115074560 | |
|  | LOC114651557 | |
|  | LOC108696475 | |
|  | AMEX60DD053844 | |
|  | UPK2.S |  |
|  | NXPE4 |  |
|  | LOC115481896 | |
|  | AMEX60DD054157 | |
|  | ASL |  |
|  | SLFN13 |  |
|  | AMEX60DD054597 | |
|  | SERPINF2 |  |
|  | MYO1B |  |
|  | TMEFF2 |  |
|  | FN1 |  |
|  | ATIC |  |
|  | ACKR3 |  |
|  | AMEX60DD055537 | |
|  | NFE2L2 |  |
|  | SLC25A12 |  |
|  | DPP4 |  |
|  | TNFAIP6 |  |
|  | MBD5 |  |
|  | AMEX60DD055902 | |
|  | AMEX60DD055929 | |
|  | AMEX60DD056045 | |
|  | PARP14 |  |
|  | MYLK |  |
|  | TFCP2L1 |  |
|  | NNMT |  |
|  | IGFBP2 |  |
|  | ANKZF1 |  |
|  | AMEX60DD056389 | |
|  | AMEX60DDU001000165 | |
|  | AMEX60DDU001000241 | |
|  | AMEX60DDU001000583 | |
|  | AMEX60DDU001001062 | |
|  | DUOX2 |  |
|  | AMEX60DDU001001245 | |
|  | AMEX60DDU001001334 | |
|  | AMEX60DDU001001537 | |
|  | AMEX60DDU001001978 | |
|  | AMEX60DDU001002112 | |
|  | AMEX60DDU001002158 | |
|  | AMEX60DDU001002237 | |
|  | AMEX60DDU001002641 | |
|  | ZBP1 |  |
|  | AMEX60DDU001003020 | |
|  | AMEX60DDU001003157 | |
|  | AMEX60DDU001003789 | |
|  | AMEX60DDU001003864 | |
|  | AMEX60DDU001004308 | |
|  | AMEX60DDU001004496 | |
|  | LOC112842973 | |
|  | AMEX60DDU001004798 | |
|  | AMEX60DDU001004834 | |
|  | AMEX60DDU001004999 | |
|  | GSONMT00027580001 | |
|  | AMEX60DDU001005272 | |
|  | AMEX60DDU001005561 | |
|  | AMEX60DDU001005779 | |
|  | AMEX60DDU001005989 | |
|  | ABCB5 |  |
|  | DR999_PMT09825 | |
|  | AMEX60DDU001006168 | |
|  | AMEX60DDU001006253 | |
|  | AMEX60DDU001006423 | |
|  | AMEX60DDU001006662 | |
|  | VMD2 |  |
|  | AMEX60DDU001006905 | |
|  | PARPI_0014716 | |
|  | AMEX60DDU001007392 | |
|  | AMEX60DDU001007407 | |
|  | AMEX60DDU001007563 | |
|  | AMEX60DDU001007697 | |
|  | AMEX60DDU001007772 | |
|  | DHX36.S |  |
|  | AMEX60DDU001008344 | |
|  | AMEX60DDU001008366 | |
|  | AMEX60DDU001008460 | |
|  | AMEX60DDU001008556 | |
|  | AMEX60DDU001008652 | |
|  | AMEX60DDU001008799 | |
|  | GASK1A |  |
|  | AMEX60DDU001008956 | |
|  | AMEX60DDU001008972 | |
|  | LOC115388170 | |
|  | AMEX60DDU001009366 | |
|  | AMEX60DDU001009438 | |
|  | AMEX60DDU001009444 | |
|  | E1301_TTI005140 | |
|  | AMEX60DDU001009763 | |
|  | AMEX60DDU001010115 | |
|  | AMEX60DDU001010124 | |
|  | AMEX60DDU001010311 | |
|  | LOC102351696 | |
|  | AMEX60DDU001011029 | |
|  | LOC112060006 | |
|  | PARPI_0022647 | |
|  | AMEX60DDU001011364 | |
|  | AMEX60DDU001011663 | |
|  | AMEX60DDU001011922 | |
|  | LOC109975224 | |
|  | AMEX60DDU001012469 | |
|  | AMEX60DDU001012479 | |
|  | AMEX60DDU001012621 | |
|  | AMEX60DDU001012632 | |
|  | AMEX60DDU001013804 | |
|  | AMEX60DDU001014139 | |
|  | AMEX60DDU001014270 | |
|  | AMEX60DDU001014509 | |
|  | AMEX60DDU001014524 | |
|  | AMEX60DDU001014766 | |
|  | AMEX60DDU001014971 | |
|  | AMEX60DDU001015076 | |
|  | AMEX60DDU001015110 | |
|  | LOC114427641 | |
|  | E1301_TTI023775 | |
|  | AMEX60DDU001015582 | |
|  | AMEX60DDU001015706 | |
|  | AMEX60DDU001015844 | |
|  | IP6K3 |  |
|  | AMEX60DDU001016491 | |
|  | AMEX60DDU001016577 | |
|  | AMEX60DDU001016610 | |
|  | AMEX60DDU001016878 | |
|  | LOC113138288 | |
|  | AMEX60DDU001017209 | |
|  | AMEX60DDU001017224 | |
|  | AMEX60DDU001017241 | |
|  | AMEX60DDU001017427 | |
|  | AMEX60DDU001018296 | |
|  | AMEX60DDU001018531 | |
|  | AMEX60DDU001018695 | |
|  | AMEX60DDU001019186 | |
|  | AMEX60DDU001019274 | |
|  | AMEX60DDU001019312 | |
|  | AMEX60DDU001019363 | |
|  | AMEX60DDU001019783 | |
|  | AMEX60DDU001019849 | |
|  | AMEX60DDU001019964 | |
|  | AMEX60DDU001020056 | |
|  | LOC102461666 | |
|  | AMEX60DDU001020224 | |
|  | AMEX60DDU001020341 | |
|  | AMEX60DDU001020540 | |
|  | A2M |  |
|  | AMEX60DDU001021334 | |
|  | AMEX60DDU001021408 | |
|  | AMEX60DDU001021885 | |
|  | LOC112546943 | |
|  | AMEX60DDU001022427 | |
|  | AMEX60DDU001022517 | |
|  | PARPI_0024169 | |
|  | AMEX60DDU001023175 | |
|  | AMEX60DDU001023711 | |
|  | AMEX60DDU001023988 | |
|  | AMEX60DDU001024180 | |
|  | AMEX60DDU001024336 | |
|  | AMEX60DDU001024484 | |
|  | MPEG1 |  |
|  | LOC104942223 | |
|  | LOC107982642 | |
|  | AMEX60DDU001025724 | |
|  | AMEX60DDU001026051 | |
|  | AMEX60DDU001026353 | |
|  | AMEX60DDU001026509 | |
|  | AMEX60DDU001026672 | |
|  | AMEX60DDU001027931 | |
|  | AMEX60DDU001027950 | |
|  | LOC112545447 | |
|  | AMEX60DDU001028679 | |
|  | AMEX60DDU001028779 | |
|  | AMEX60DDU001028801 | |
|  | AMEX60DDU001028916 | |
|  | AMEX60DDU001029224 | |
|  | APELA |  |
|  | AMEX60DDU001029317 | |
|  | AMEX60DDU001029461 | |
|  | POL |  |
|  | AMEX60DDU001029731 | |
|  | AMEX60DDU001029766 | |
|  | AMEX60DDU001029845 | |
|  | AMEX60DDU001029970 | |
|  | PARPI_0024411 | |
|  | AMEX60DDU001030328 | |
|  | AMEX60DDU001030652 | |
|  | AMEX60DDU001030736 | |
|  | AMEX60DDU001031094 | |
|  | AMEX60DDU001031110 | |
|  | AMEX60DDU001031122 | |
|  | LOC103305703 | |
|  | LOC106930792 | |
|  | PARPI_0007370 | |
|  | TMEM161B |  |
|  | AMEX60DDU001032305 | |
|  | GNMT |  |
|  | AMEX60DDU001032976 | |
|  | AMEX60DDU001033149 | |
|  | AMEX60DDU001033367 | |
|  | AMEX60DDU001033505 | |
|  | AMEX60DDU001034126 | |
|  | AMEX60DDU001034387 | |
|  | AMEX60DDU001034456 | |
|  | LOC115381268 | |
|  | AMEX60DDU001034628 | |
|  | AMEX60DDU001035335 | |
|  | AMEX60DDU001035475 | |
|  | AMEX60DDU001035584 | |
|  | AMEX60DDU001035636 | |
|  | SACS |  |
|  | AMEX60DDU001036000 | |
|  | AMEX60DDU001036067 | |
|  | AMEX60DDU001036341 | |
|  | AMEX60DDU001036438 | |
|  | AMEX60DDU001036548 | |
|  | AMEX60DDU001036596 | |
|  | AMEX60DDU001036717 | |
|  | AMEX60DDU001036843 | |
|  | AMEX60DDU001037272 | |
|  | LOC101952591 | |
|  | AMEX60DDU001037379 | |
|  | AMEX60DDU001037386 | |
|  | AMEX60DDU001037438 | |
|  | AMEX60DDU001037479 | |
|  | AMEX60DDU001037741 | |
|  | AMEX60DDU001037955 | |
|  | AMEX60DDU001038078 | |
|  | AMEX60DDU001038085 | |
|  | AMEX60DDU001038165 | |
|  | LOC115075634 | |
|  | AMEX60DDU001038657 | |
|  | AMEX60DDU001038918 | |
|  | AMEX60DDU001038928 | |
|  | AMEX60DDU001039055 | |
|  | AMEX60DDU001039810 | |
|  | AMEX60DDU001040368 | |
|  | AMEX60DDU001040700 | |
|  | KRT8 |  |
|  | LOC105373926 | |
|  | AMEX60DDU001040947 | |
|  | AMEX60DDU001041209 | |
|  | AMEX60DDU001041406 | |
|  | AMEX60DDU001041472 | |
|  | AMEX60DDU001041801 | |
