## Supplementary material for "Retinoic acid breakdown is required for proximodistal positional identity during amphibian limb regeneration": Table S4

|  |  |  |  |  | PD duplication | | | | |
| --- | --- | --- | --- | --- | --- | --- | --- | --- | --- |
| Dose | Amputation plane | Total animals | No duplication | No regeneration | Half radius/ulna | Full radius/ulna | Half humerus | Full humerus | Shoulder |
| DMSO | Proximal | 6 | 6 | ― | ― | ― | ― | ― | ― |
| DMSO | Distal | 6 | 6 | ― | ― | ― | ― | ― | ― |
| 0.1uM DIS | Proximal | 7 | 7 | ― | ― | ― | ― | ― | ― |
| 0.1uM DIS | Distal | 7 | 7 | ― | ― | ― | ― | ― | ― |
| 1uM DIS | Proximal | 6 | 5 | 1 | ― | ― | ― | ― | ― |
| 1uM DIS | Distal | 6 | 6 | ― | ― | ― | ― | ― | ― |
| 5uM DIS | Proximal | 5 | 5 | ― | ― | ― | ― | ― | ― |
| 5uM DIS | Distal | 5 | 5 | ― | ― | ― | ― | ― | ― |
| 0.1uM DIS/1 uM TAL | Proximal | 4 | 4 | ― | ― | ― | ― | ― | ― |
| 0.1uM DIS/1 uM TAL | Distal | 4 | ― | ― | ― | 4 | ― | ― | ― |
| 1uM DIS/1 uM TAL | Proximal | 5 | 5 | ― | ― | ― | ― | ― | ― |
| 1uM DIS/1 uM TAL | Distal | 5 | 2 | 1 | 2 | ― | ― | ― | ― |
