## Supplementary material for "Retinoic acid breakdown is required for proximodistal positional identity during amphibian limb regeneration": Table S3

|  |  |  |  |  | PD duplication | | | | |
| --- | --- | --- | --- | --- | --- | --- | --- | --- | --- |
| Dose | Amputation plane | Total animals | No duplication | No regeneration | Half radius/ulna | Full radius/ulna | Half humerus | Full humerus | Shoulder |
| DMSO | Proximal | 14 | 14 | ― | ― | ― | ― | ― | ― |
| DMSO | Distal | 14 | 14 | ― | ― | ― | ― | ― | ― |
| 0.1 µM TAL | Proximal | 13 | 13 | ― | ― | ― | ― | ― | ― |
| 0.1 µM TAL | Distal | 13 | 1 | ― | 4 | 8 | ― | ― | ― |
| 1 µM TAL | Proximal | 12 | 12 | ― | ― | ― | ― | ― | ― |
| 1 µM TAL | Distal | 12 | ― | ― | ― | ― | 4 | 8 | ― |
| 5 µM TAL | Proximal | 13 | 2 | 11 | ― | ― | ― | ― | ― |
| 5 µM TAL | Distal | 13 | ― | 12 | ― | ― | ― | 1 | ― |
