## Supplementary material for "Retinoic acid breakdown is required for proximodistal positional identity during amphibian limb regeneration": Table S2

| **Meis1** | |
| --- | --- |
| Meis1_B1 | gAggAgggCAgCAAACggAAATCCCACCGTAGTGGGGCAAGTCGT |
| Meis1_B1 | ATGGTGCTGGGGATACCCACTCCGTTAgAAgAgTCTTCCTTTACg |
| Meis1_B1 | gAggAgggCAgCAAACggAAGCTGCATGGACCTGGCCGCGTGGGG |
| Meis1_B1 | GAGGGCCGTGGTTCAGGTGGTGGACTAgAAgAgTCTTCCTTTACg |
| Meis1_B1 | gAggAgggCAgCAAACggAATGTGCGGGTACTGGTGCGAGTGCAG |
| Meis1_B1 | CGGGCGGCATGGCGTTCGTGTGGCCTAgAAgAgTCTTCCTTTACg |
| Meis1_B1 | gAggAgggCAgCAAACggAAGGGCGTCGTTGACCGAGGAGCCCAT |
| Meis1_B1 | CGTAGATGGAGTCCTTGTCCCGCTTTAgAAgAgTCTTCCTTTACg |
| Meis1_B1 | gAggAgggCAgCAAACggAAAACACACGTCCCCGCCGGCTACCCC |
| Meis1_B1 | CGATGTCTTCATTGAAGGACTCGGATAgAAgAgTCTTCCTTTACg |
| Meis1_B1 | gAggAgggCAgCAAACggAACTGCTCGGATCTGTTTAGCGAACAC |
| Meis1_B1 | CTGGGTTCGACGAAAATAAAGGCTTTAgAAgAgTCTTCCTTTACg |
| Meis1_B1 | gAggAgggCAgCAAACggAATCGCCTGGATCATCAAGTTATCCAG |
| Meis1_B1 | CCAGCAGATGAAACCGTAACACTTGTAgAAgAgTCTTCCTTTACg |
| Meis1_B1 | gAggAgggCAgCAAACggAACGCACAGCTCGTGTACCTTTTCTAA |
| Meis1_B1 | AGCTGATGTACCGGTGGCAGAAATTTAgAAgAgTCTTCCTTTACg |
| Meis1_B1 | gAggAgggCAgCAAACggAAGGTCGATGGGCATCTTCCCTTTCAA |
| Meis1_B1 | ATCCGCCTTCCCGGTCGTCGATCACTAgAAgAgTCTTCCTTTACg |
| Meis1_B1 | gAggAgggCAgCAAACggAAGGCCCATGTCTTCGCTGTCCGACTT |
| Meis1_B1 | AAGGCTGGTCTGTGAGGTTCGCCGGTAgAAgAgTCTTCCTTTACg |
| Meis1_B1 | gAggAgggCAgCAAACggAACCGCTGGAGGGTCCTGGCGTTCCCC |
| Meis1_B1 | TTGTCCCCACTGTGTGACGTGTGCCTAgAAgAgTCTTCCTTTACg |
| Meis1_B1 | gAggAgggCAgCAAACggAATCCAATCCATCACCTTGCTCACTAC |
| Meis1_B1 | CCTGTGCTGGGAGAAGCTACACTGTTAgAAgAgTCTTCCTTTACg |
| Meis1_B1 | gAggAgggCAgCAAACggAAACTTTAGGGAAGATGCCCCGCTTTT |
| Meis1_B1 | AGCCACGCTCGCATAATATTTGTAGTAgAAgAgTCTTCCTTTACg |
| Meis1_B1 | gAggAgggCAgCAAACggAACCCGTGTCCTGTGCCAGCTGCTTTT |
| Meis1_B1 | CAATTGTTCACTTGAAGGATAGTGATAgAAgAgTCTTCCTTTACg |
| Meis1_B1 | gAggAgggCAgCAAACggAAACTATTCTTCTCCTGGCGTTGATAA |
| Meis1_B1 | CGGTTGGACTGGTCTATCATGGGCTTAgAAgAgTCTTCCTTTACg |
| Meis1_B1 | gAggAgggCAgCAAACggAACTGTATGGTGTACCTTGACTTACTG |
| Meis1_B1 | AATCCTCCCATGGGCTGCCCGTCTTTAgAAgAgTCTTCCTTTACg |
| Meis1_B1 | gAggAgggCAgCAAACggAACCCATGTGCTGCTGCCCATCCATCA |
| Meis1_B1 | ATACTTTGCAGTCCTGGCGCTCTGATAgAAgAgTCTTCCTTTACg |
| Meis1_B1 | gAggAgggCAgCAAACggAACCACCCTGCGAGACGTAATCCCCTG |
| Meis1_B1 | GGCTGTCCCATACACATACCCATTGTAgAAgAgTCTTCCTTTACg |
| Meis1_B1 | gAggAgggCAgCAAACggAATGGCCCATCTGTGGTGGGGTGTAAC |
| Meis1_B1 | GGCCCATGACGCAGCTGAGCAGGATTAgAAgAgTCTTCCTTTACg |
| Meis1_B1 | gAggAgggCAgCAAACggAATGTCCAGGAATGTAGGTATGCATTG |
| Meis1_B1 | TGCATCATCATTGCTGGGTGGTGAGTAgAAgAgTCTTCCTTTACg |
| Meis1_B1 | gAggAgggCAgCAAACggAAGACATTCCAGGGTGGGGTGGTCCTC |
| Meis1_B1 | AGCATTGCGGGGCTAGTCGCAGACATAgAAgAgTCTTCCTTTACg |
| Meis1_B1 | gAggAgggCAgCAAACggAACCGCCCATGGTTGGGTCTCCTGTGT |
| Meis1_B1 | TACTGAGCATGGATATCCATGACTTTAgAAgAgTCTTCCTTTACg |
| Meis1_B1 | gAggAgggCAgCAAACggAACATAGCCATTGCAGAAAGTTATTTT |
| Meis1_B1 | CCAGAGTAGATGCTGAGCACGAATCTAgAAgAgTCTTCCTTTACg |
| Meis1_B1 | gAggAgggCAgCAAACggAATATGAAGAAATTGGACTACCTCTTG |
| Meis1_B1 | TTGGTGTTTCCTGCTTTTGTAAGTCTAgAAgAgTCTTCCTTTACg |
| Meis1_B1 | gAggAgggCAgCAAACggAATTATTTAGTGTGTCCCCAAATTTCC |
| Meis1_B1 | TCTTTGTTCTCTTAATGTCTTATATTAgAAgAgTCTTCCTTTACg |
| Meis1_B1 | gAggAgggCAgCAAACggAAGAAACAACACACATAGTGTGGAAAA |
| Meis1_B1 | GGCTTCTGGAGGAAGTGAAGAGCTATAgAAgAgTCTTCCTTTACg |
| Meis1_B1 | gAggAgggCAgCAAACggAATCTGCAGGATAACGGTAAGGCTTTT |
| Meis1_B1 | ACTCCTGCAAATAGGACCTTTATGTTAgAAgAgTCTTCCTTTACg |
| Meis1_B1 | gAggAgggCAgCAAACggAACTGACGTAGTTACTTGAATGCTCTA |
| Meis1_B1 | CCTAGGGAAGAAGAACAATCCTTCTTAgAAgAgTCTTCCTTTACg |
| Meis1_B1 | gAggAgggCAgCAAACggAAAGGGGAGACAGAGTGCATGGCTGAA |
| Meis1_B1 | CAAGCCCCAGGCAAGAGAGGAGAGCTAgAAgAgTCTTCCTTTACg |
| Meis1_B1 | gAggAgggCAgCAAACggAACGAGTAAATGAATTAGGCATGCAAA |
| Meis1_B1 | CATCTGTTCAGGCCAATTCAAATATTAgAAgAgTCTTCCTTTACg |
| Meis1_B1 | gAggAgggCAgCAAACggAAAAATCATTGTTGGCGACGGCGTTTT |
| Meis1_B1 | CTTAAGACATTGCTTGCAACAGCTGTAgAAgAgTCTTCCTTTACg |
| Meis1_B1 | gAggAgggCAgCAAACggAAGTTACCATGCCTCCTCCTACCGGTT |
| Meis1_B1 | GGCTCCAGGCACGCTTGCTGCCTCTTAgAAgAgTCTTCCTTTACg |
| Meis1_B1 | gAggAgggCAgCAAACggAAAACCTTGATCCGCTGATGGGACATT |
| Meis1_B1 | GCGTACGGTTGATGCTACCGCATGGTAgAAgAgTCTTCCTTTACg |
| Meis1_B1 | gAggAgggCAgCAAACggAATCGAAATAAAGTCCACATATTAAAA |
| Meis1_B1 | AAATGCACACATGACTGATGCTTTGTAgAAgAgTCTTCCTTTACg |
| Meis1_B1 | gAggAgggCAgCAAACggAACATAATGCTCCATGGTGGACTGTGG |
| Meis1_B1 | TTGCACAAAATTTATTTACAATGTATAgAAgAgTCTTCCTTTACg |
| Meis1_B1 | gAggAgggCAgCAAACggAAGAATTGCAATGATACAGTTCATTTT |
| Meis1_B1 | TCATACCAAACTGCTACTTTTACAATAgAAgAgTCTTCCTTTACg |
| Meis1_B1 | gAggAgggCAgCAAACggAAAGCTCCTTGTTTTCTTGATAGAAAA |
| Meis1_B1 | GGTGTGCAATAATTACTAATCATGATAgAAgAgTCTTCCTTTACg |
| **Meis2** | |
| Meis2_B2 | CCTCgTAAATCCTCATCAAACCACTCCGTCCATCCCGTAGTGGGG |
| Meis2_B2 | GGTCCCCGTACATCGACGTGGGCACAAATCATCCAgTAAACCgCC |
| Meis2_B2 | CCTCgTAAATCCTCATCAAATGCTGGGCATGACATTGGGGTGGGG |
| Meis2_B2 | GGGCGTCGTTAACAGCCGACCCCATAAATCATCCAgTAAACCgCC |
| Meis2_B2 | CCTCgTAAATCCTCATCAAATCTCAAAGACCAGGGCGAGCAGGGG |
| Meis2_B2 | GAGGGGTGCAGGTGGCCAGCTCGCAAAATCATCCAgTAAACCgCC |
| Meis2_B2 | CCTCgTAAATCCTCATCAAACATCCCCGCCCGCCACACCGGGCTC |
| Meis2_B2 | CCTCGTTGAAGGAGTCGGAGGAGCAAAATCATCCAgTAAACCgCC |
| Meis2_B2 | CCTCgTAAATCCTCATCAAACTTTCAGGCAACTTATATAGCGGTG |
| Meis2_B2 | CGATGACCAGGTCGATGGGCATCTTAAATCATCCAgTAAACCgCC |
| Meis2_B2 | CCTCgTAAATCCTCATCAAACCGACTTGGAGCTGCCGTCCCTCTC |
| Meis2_B2 | TGGAGGAGCCGGAGAGCTCCTCGTGAAATCATCCAgTAAACCgCC |
| Meis2_B2 | CCTCgTAAATCCTCATCAAAGGCCCGGCGTGCCGGCCGAATGCGT |
| Meis2_B2 | TCTGAGAAGCGTGCCCTCCACTGGAAAATCATCCAgTAAACCgCC |
| Meis2_B2 | CCTCgTAAATCCTCATCAAAGGTCGTCGTCGTCGCCCGTGCCAGG |
| Meis2_B2 | TCTTCTGGCGCTTCTTGTCTTTGTCAAATCATCCAgTAAACCgCC |
| Meis2_B2 | CCTCgTAAATCCTCATCAAATGGCTACTTTGGGGAAAATGCCTCG |
| Meis2_B2 | GGAAGAGCCAGGCCCTCATGATATTAAATCATCCAgTAAACCgCC |
| Meis2_B2 | CCTCgTAAATCCTCATCAAATGTACGCTGCTCCTTGACTCACTGA |
| Meis2_B2 | AGCTTCCCATTGGCTGGCCCTCGGGAAATCATCCAgTAAACCgCC |
| Meis2_B2 | CCTCgTAAATCCTCATCAAACTCCCGGAGGTCCGTAGTCCCCAGA |
| Meis2_B2 | ACTGTGCCATACTCATGCCCATCGGAAATCATCCAgTAAACCgCC |
| Meis2_B2 | CCTCgTAAATCCTCATCAAAGGGTAAACTGGGGAGCAGTGTAACT |
| Meis2_B2 | GTCCGTGTCTCAACTGAGAAAGGTGAAATCATCCAgTAAACCgCC |
| Meis2_B2 | CCTCgTAAATCCTCATCAAAGGCTTGGCAAATAGGGATGCATTGG |
| Meis2_B2 | GGACCATGGCTGGCCGATGATGAGGAAATCATCCAgTAAACCgCC |
| Meis2_B2 | CCTCgTAAATCCTCATCAAAATAGTCATTCCAGAGTGGGCAGGGG |
| Meis2_B2 | TTGCGCATGGAAGGGCCCTGTGCTGAAATCATCCAgTAAACCgCC |
| Meis2_B2 | CCTCgTAAATCCTCATCAAATGTCCGCCAAAGCCGGGATCTGCAG |
| Meis2_B2 | TACAGTTAGGCATGGATGTCCATCAAAATCATCCAgTAAACCgCC |
| Meis2_B2 | CCTCgTAAATCCTCATCAAATGTTCTGTTTTCCCTTGAGTTCCCT |
| Meis2_B2 | TGGTCCAAGCTCAAATGTCTGAGAAAAATCATCCAgTAAACCgCC |
| Meis2_B2 | CCTCgTAAATCCTCATCAAAGTTGGTCCTCTGTTGTTATTGAAAA |
| Meis2_B2 | AATTCACATTTGTGTTCTTGGTTTAAAATCATCCAgTAAACCgCC |
| Meis2_B2 | CCTCgTAAATCCTCATCAAATCTCGCGCTCTCTTTCTGGTGTTTT |
| Meis2_B2 | AGTAGTTTTAAACTCTTTGAGTCTTAAATCATCCAgTAAACCgCC |
| Meis2_B2 | CCTCgTAAATCCTCATCAAAAACTTAGTTCCTATGCTTATATACT |
| Meis2_B2 | TCGCGTGATGAAAGAAATGTACAAGAAATCATCCAgTAAACCgCC |
| Meis2_B2 | CCTCgTAAATCCTCATCAAAGGAACAACACACATAGTGTGGAAAA |
| Meis2_B2 | CTGTACCAAAACACAGTAATTCTTTAAATCATCCAgTAAACCgCC |
| Meis2_B2 | CCTCgTAAATCCTCATCAAAGTCTTTCAAAGATGGAGACCTTAGC |
| Meis2_B2 | TCCTGCTCTCTGGTGAAGCAGGCTGAAATCATCCAgTAAACCgCC |
| Meis2_B2 | CCTCgTAAATCCTCATCAAACACTGGTGCGACCGTCACTGTCGGC |
| Meis2_B2 | ACAGGAGGCAGTGCAAGGAGAGATGAAATCATCCAgTAAACCgCC |
| Meis2_B2 | CCTCgTAAATCCTCATCAAAGGCTTGAAACTGAAAGGGGACTTTT |
| Meis2_B2 | AGAAGGGCCTCCGTGTCCTCTCTGAAAATCATCCAgTAAACCgCC |
| Meis2_B2 | CCTCgTAAATCCTCATCAAAAAGAAGAGTCCAGCCAGAAAGCAAA |
| Meis2_B2 | CTGGGTCTTCACCTGATAGCACAGAAAATCATCCAgTAAACCgCC |
| **Hoxa9** | |
| Hoxa9_B1 | gAggAgggCAgCAAACggAACCAGCTCCTCGCTCTCGTGGATGAG |
| Hoxa9_B1 | CCGCGGCGGCGGCATAGCGCGACTGTAgAAgAgTCTTCCTTTACg |
| Hoxa9_B1 | gAggAgggCAgCAAACggAAACGGCGTGAACTCGGGGTGCTCCCC |
| Hoxa9_B1 | ACACCGGGCTCTTGGACTGGAAGCTTAgAAgAgTCTTCCTTTACg |
| Hoxa9_B1 | gAggAgggCAgCAAACggAATAGACGGCGGGCACGCTGCCGCCGG |
| Hoxa9_B1 | TGGTGCACGTAGGGATGGTGGTGGTTAgAAgAgTCTTCCTTTACg |
| Hoxa9_B1 | gAggAgggCAgCAAACggAAACCTGCTGGCCTCGGGCGCCCCTGG |
| Hoxa9_B1 | GCATGGGCTCCAGCCAGGAGCGCATTAgAAgAgTCTTCCTTTACg |
| Hoxa9_B1 | gAggAgggCAgCAAACggAAGCAGCCCGGGGAAGGAGAGCGCGCC |
| Hoxa9_B1 | GCTTGATGCCGTAGTGCCTGGCAGCTAgAAgAgTCTTCCTTTACg |
| Hoxa9_B1 | gAggAgggCAgCAAACggAATGCTGTCGAAGGTGGTGCAGTCCCC |
| Hoxa9_B1 | CGTAGTCGGACAGGGAGAGCGTGTGTAgAAgAgTCTTCCTTTACg |
| Hoxa9_B1 | gAggAgggCAgCAAACggAAAGATCGAGCGGCCGGCCAGCCGCCC |
| Hoxa9_B1 | GGGCACACGGGTGGGAACGGGTCCGTAgAAgAgTCTTCCTTTACg |
| Hoxa9_B1 | gAggAgggCAgCAAACggAATGGGCGGCACTGGGCGGCCCGAGGT |
| Hoxa9_B1 | CATGTGTGAAAGGATGACAGTACCTTAgAAgAgTCTTCCTTTACg |
| Hoxa9_B1 | gAggAgggCAgCAAACggAACCAGAGCGCCTGCTTTTGCTTTCCT |
| Hoxa9_B1 | TATGTACAGGCCAGCACGCCCCAGGTAgAAgAgTCTTCCTTTACg |
| Hoxa9_B1 | gAggAgggCAgCAAACggAAGTGTGCATTCTGAAAAGGTAGTTTT |
| Hoxa9_B1 | TAAATTAACTATACAGTCCGTTCGGTAgAAgAgTCTTCCTTTACg |
| Hoxa9_B1 | gAggAgggCAgCAAACggAAGAGCTACATATATACATTTATATAT |
| Hoxa9_B1 | TTCCTGCCGACAATTAGCAAATAAATAgAAgAgTCTTCCTTTACg |
| Hoxa9_B1 | gAggAgggCAgCAAACggAACAAGTTTCAAGTATCTTACAGCAGG |
| Hoxa9_B1 | CACAAACACATTTACCACGACAAAATAgAAgAgTCTTCCTTTACg |
| Hoxa9_B1 | gAggAgggCAgCAAACggAATACAGCACCCCAGCACGCACGTTGA |
| Hoxa9_B1 | ATGTCTAGCTTACACAACCACAAAGTAgAAgAgTCTTCCTTTACg |
| Hoxa9_B1 | gAggAgggCAgCAAACggAACTTATTAAACAACCTGAGTTCAAAG |
| Hoxa9_B1 | AACACTACCGAGCTATACCCTATGCTAgAAgAgTCTTCCTTTACg |
| Hoxa9_B1 | gAggAgggCAgCAAACggAACACGCCATCTAATCACAGAAAACAC |
| Hoxa9_B1 | CAAACCCGTGATTGTTGCAGGCCAATAgAAgAgTCTTCCTTTACg |
| Hoxa9_B1 | gAggAgggCAgCAAACggAAAGCCACTGTCCAGCACACAATCCAC |
| Hoxa9_B1 | AGCCACAAGGGACGCAGGCTGCGTGTAgAAgAgTCTTCCTTTACg |
| Hoxa9_B1 | gAggAgggCAgCAAACggAAAAAGTCCTTCTTCCTCTCCCCAATG |
| Hoxa9_B1 | GGGCGAAAACAAAAGTCCGAGCCAATAgAAgAgTCTTCCTTTACg |
| Hoxa9_B1 | gAggAgggCAgCAAACggAAGCTCCCAAACCGAATGTCTGTCCAG |
| Hoxa9_B1 | GTCAACCTTAGTTTTCACAAACGTATAgAAgAgTCTTCCTTTACg |
| Hoxa9_B1 | gAggAgggCAgCAAACggAAGCATGAAAGGAGGTGAATTTCTTTT |
| Hoxa9_B1 | TCAATAGATTTAACTTCATAACAACTAgAAgAgTCTTCCTTTACg |
| Hoxa9_B1 | gAggAgggCAgCAAACggAACTGCATCGCATACATTCAAGAGGGG |
| Hoxa9_B1 | ACTATCTGTACGTATACAGCGTGTGTAgAAgAgTCTTCCTTTACg |
| Hoxa9_B1 | gAggAgggCAgCAAACggAAGGCACCGGGTGGGAAGGCCTTCGCT |
| Hoxa9_B1 | GGCGGCCGTGGGGTGTCCTATAAGTTAgAAgAgTCTTCCTTTACg |
| Hoxa9_B1 | gAggAgggCAgCAAACggAAGGGTATTTCAGAAGGAGTTCTTGTG |
| Hoxa9_B1 | GCTGTACTGTATTTAAATATATGTGTAgAAgAgTCTTCCTTTACg |
| Hoxa9_B1 | gAggAgggCAgCAAACggAATTATGCAATCATTTGAGCCATAAAA |
| Hoxa9_B1 | CCTGCGATCGGAGACCTCCACATAATAgAAgAgTCTTCCTTTACg |
| Hoxa9_B1 | gAggAgggCAgCAAACggAACACCGGGCCCGCGTTACAGGCGAAA |
| Hoxa9_B1 | CTTTCCTATCATAATTAATGACAGCTAgAAgAgTCTTCCTTTACg |
| **Hoxa11** | |
| Hoxa11_B2 | CCTCgTAAATCCTCATCAAAACGGGACACGCTCATCAAAATCCAT |
| Hoxa11_B2 | AACTTGGCAAGTACATGTTAGAGGAAAATCATCCAgTAAACCgCC |
| Hoxa11_B2 | CCTCgTAAATCCTCATCAAAAATCGGGGCCGGAGACGTAGTAAGT |
| Hoxa11_B2 | GGGGCAGGAAGGAGGGCAGGCTGGAAAATCATCCAgTAAACCgCC |
| Hoxa11_B2 | CCTCgTAAATCCTCATCAAAAAGGCATGGGGCGAGAAGCCGGGTT |
| Hoxa11_B2 | CCTGGGGCAGGTTGGACGAGTAGGAAAATCATCCAgTAAACCgCC |
| Hoxa11_B2 | CCTCgTAAATCCTCATCAAAGGAAGGTGACCTCCCGCACGGGCTG |
| Hoxa11_B2 | TGCTGGCGGGGTCAATGGCGTACTCAAATCATCCAgTAAACCgCC |
| Hoxa11_B2 | CCTCgTAAATCCTCATCAAAGGGCCAGGTTGCTCCGCGGGTGCCA |
| Hoxa11_B2 | GCATCAGCTCCTCGGCGGAGTAGCAAAATCATCCAgTAAACCgCC |
| Hoxa11_B2 | CCTCgTAAATCCTCATCAAAAGCCGGGTGGTGGTAGGGGCTGGGG |
| Hoxa11_B2 | GCTGTAAAAGTTGGAGGAGGCGCCCAAATCATCCAgTAAACCgCC |
| Hoxa11_B2 | CCTCgTAAATCCTCATCAAACGGCAGGACCCCGTTCCTGCCCACG |
| Hoxa11_B2 | CGTCTCGAAGAACTGGTCGAAAGCCAAATCATCCAgTAAACCgCC |
| Hoxa11_B2 | CCTCgTAAATCCTCATCAAAGGGTCCGCTCTCGGGGCCACCGTAG |
| Hoxa11_B2 | GCAGCCCTTGTCCCCAGCGTAGTCCAAATCATCCAgTAAACCgCC |
| Hoxa11_B2 | CCTCgTAAATCCTCATCAAAGCCCGCAACCGCCGGGGATCCCTTC |
| Hoxa11_B2 | GCCTCGACAGGCCTCGGCGCTGGGAAAATCATCCAgTAAACCgCC |
| Hoxa11_B2 | CCTCgTAAATCCTCATCAAAACTCTCGGCTCCCCGCCGGTCCGCC |
| Hoxa11_B2 | GCTGGAGCTGCTGCCGCCGCCGCCAAAATCATCCAgTAAACCgCC |
| Hoxa11_B2 | CCTCgTAAATCCTCATCAAACTCGTTGTTGCCGGAAGAGGACTCC |
| Hoxa11_B2 | ATTGGGAGCGCTGCCGGACGCCTTCAAATCATCCAgTAAACCgCC |
| Hoxa11_B2 | CCTCgTAAATCCTCATCAAAGCACCTCTTCTTTCGGGTGCGCTGC |
| Hoxa11_B2 | TTCGCGGATCTGGTACTTGGTGTACAAATCATCCAgTAAACCgCC |
| Hoxa11_B2 | CCTCgTAAATCCTCATCAAATCCTTGTTGATGTAGACGCTGAAAA |
| Hoxa11_B2 | AGCATCCGCGACAGCTGCAGCCGCTAAATCATCCAgTAAACCgCC |
| Hoxa11_B2 | CCTCgTAAATCCTCATCAAAATTGCAAGCGGTCCCGGTTGATTTT |
| Hoxa11_B2 | CTCAGAGCAGCGGGTTCGCGGAGTAAAATCATCCAgTAAACCgCC |
| Hoxa11_B2 | CCTCgTAAATCCTCATCAAATCCAATTGATAGTTAACTATTAAAA |
| Hoxa11_B2 | AACACATATGTGCATTTAGCCATCGAAATCATCCAgTAAACCgCC |
| Hoxa11_B2 | CCTCgTAAATCCTCATCAAAGGGAATCTCCGCGGCCAAGGCTGGA |
| Hoxa11_B2 | TCGGCGAGTAGGACGTCCGCGGGGAAAATCATCCAgTAAACCgCC |
| Hoxa11_B2 | CCTCgTAAATCCTCATCAAACCCGTGCACCGTTGCAATCCTAAAA |
| Hoxa11_B2 | CTTTAAAACCAAGCCTGTTTGGCAAAAATCATCCAgTAAACCgCC |
| Hoxa11_B2 | CCTCgTAAATCCTCATCAAAACGGAAGTGCGCCTGCAATTCTGGT |
| Hoxa11_B2 | CCTGCACGAAGGTCAAAGCTGATTTAAATCATCCAgTAAACCgCC |
| Hoxa11_B2 | CCTCgTAAATCCTCATCAAAGGACGAACCAGCATTCGCCGATTTT |
| Hoxa11_B2 | GCTTCCGGCGATTGTTTAGTCGCGTAAATCATCCAgTAAACCgCC |
| Hoxa11_B2 | CCTCgTAAATCCTCATCAAAATCTCCAGGCGGCGCTCCCGCGTCA |
| Hoxa11_B2 | CGGTCGGAGAGGTGCACGCTGCGACAAATCATCCAgTAAACCgCC |
| Hoxa11_B2 | CCTCgTAAATCCTCATCAAACCGTGAGTTCGCGGACTCGGCTCTC |
| Hoxa11_B2 | CGGGGATCAGGAGAAGCCGAAGTTGAAATCATCCAgTAAACCgCC |
| Hoxa11_B2 | CCTCgTAAATCCTCATCAAACTGGCTTTCCCGGGACTCCTGGCGC |
| Hoxa11_B2 | AAGCGTGCCAGGCAACAGTGTGCAAAAATCATCCAgTAAACCgCC |
| Hoxa11_B2 | CCTCgTAAATCCTCATCAAATTCTGGAAACAGCGACCATCAAAGC |
| Hoxa11_B2 | ACTCGGCAGTTCTGCAGTTAGCAAGAAATCATCCAgTAAACCgCC |
| Hoxa11_B2 | CCTCgTAAATCCTCATCAAATCGCAATTTCAATCCGTGCTGTTCC |
| Hoxa11_B2 | CCTGTGCCCGCCAGTTCGGGGAAGGAAATCATCCAgTAAACCgCC |
| **Hoxa13** | |
| Hoxa13_B3 | gTCCCTgCCTCTATATCTTTGACGAGCCCTGCGCCGAGCAGGGGC |
| Hoxa13_B3 | AAATAACCGTAGGGCAGCGCGGCCCTTCCACTCAACTTTAACCCg |
| Hoxa13_B3 | gTCCCTgCCTCTATATCTTTACCCTGCACGGGTAGTATCCGCTTC |
| Hoxa13_B3 | CAGGACTTGATGCCGCCGTGGTGGCTTCCACTCAACTTTAACCCg |
| Hoxa13_B3 | gTCCCTgCCTCTATATCTTTTTGTCGGCGAAGGAGGAGGGCTGCG |
| Hoxa13_B3 | CCCGCCGAGCCCGACGTGTCCATGTTTCCACTCAACTTTAACCCg |
| Hoxa13_B3 | gTCCCTgCCTCTATATCTTTTCCTTGGCCCTGGAGGTGAACTCCT |
| Hoxa13_B3 | GCCGCGTAGCCCTGGTAGAAGGCGATTCCACTCAACTTTAACCCg |
| Hoxa13_B3 | gTCCCTgCCTCTATATCTTTTAACCGGGCACCGGCTGGTACGGGC |
| Hoxa13_B3 | GTGGGCACCATGGTGGGCATATCCATTCCACTCAACTTTAACCCg |
| Hoxa13_B3 | gTCCCTgCCTCTATATCTTTAGGGGCTCGTGCCTGGGCTCGCCGG |
| Hoxa13_B3 | CAGGGCTGGTAGGGCTCCATGGGCATTCCACTCAACTTTAACCCg |
| Hoxa13_B3 | gTCCCTgCCTCTATATCTTTTGCCCATTCCAGCCATTGGTGAGGG |
| Hoxa13_B3 | TGCCCTTGCTCCTTGGAACAGTACATTCCACTCAACTTTAACCCg |
| Hoxa13_B3 | gTCCCTgCCTCTATATCTTTTTCCGGCCTCGCCGGTACGAGTTCG |
| Hoxa13_B3 | TGGACCTTGGTGTACGGCACGCGCTTTCCACTCAACTTTAACCCg |
| Hoxa13_B3 | gTCCCTgCCTCTATATCTTTGCGTACTCGCGCTCCAGTTCCTTCA |
| Hoxa13_B3 | TTGTCCTTGGTAATGAACTTATTCGTTCCACTCAACTTTAACCCg |
| Hoxa13_B3 | gTCCCTgCCTCTATATCTTTTTGGTGGTGGCCGATATCCGCCTCC |
| Hoxa13_B3 | CAAATGGTGACCTGGCGCTCGGAGATTCCACTCAACTTTAACCCg |
| Hoxa13_B3 | gTCCCTgCCTCTATATCTTTTTCTCTTTGACCCTCCTGTTTTGGA |
| Hoxa13_B3 | GTGGTCTTGAGTTTGTTGATGACCTTTCCACTCAACTTTAACCCg |
| Hoxa13_B3 | gTCCCTgCCTCTATATCTTTTTCTTTTAGCTTCGGCTTAGGTTTT |
| Hoxa13_B3 | AGCTTCATTTCCTTGTAACTGTCAGTTCCACTCAACTTTAACCCg |
| Hoxa13_B3 | gTCCCTgCCTCTATATCTTTGTTCGCGAATTCTGGTGACCAATGA |
| Hoxa13_B3 | AAACAACATGTCACAAATTCCATAATTCCACTCAACTTTAACCCg |
| Hoxa13_B3 | gTCCCTgCCTCTATATCTTTGGCAATTGTCTTCATGAACCAGTTT |
| Hoxa13_B3 | ACAGTTCCAAAGAGGCACACATTGCTTCCACTCAACTTTAACCCg |
| Hoxa13_B3 | gTCCCTgCCTCTATATCTTTCTGAAATGCGTGTCCGTTGCTCAGT |
| Hoxa13_B3 | CGCTGGCGTGAGAACGCGTACGTAGTTCCACTCAACTTTAACCCg |
| Hoxa13_B3 | gTCCCTgCCTCTATATCTTTGTTTTCACCGTCTTCCCCATTAAAA |
| Hoxa13_B3 | GCACACACCTTGTACAACGGGGTGTTTCCACTCAACTTTAACCCg |
| Hoxa13_B3 | gTCCCTgCCTCTATATCTTTATGCAGTCTGGGTGTAAGTTTGGGC |
| Hoxa13_B3 | TATAAATTACCTGAGCAGGCATTCCTTCCACTCAACTTTAACCCg |
| Hoxa13_B3 | gTCCCTgCCTCTATATCTTTTTTTCAATATCCGCTTAATATAAAA |
| Hoxa13_B3 | TTTCTAAATACTTGCTTTCAAATGATTCCACTCAACTTTAACCCg |
| Hoxa13_B3 | gTCCCTgCCTCTATATCTTTATTCGCACACACCACAAGCTCTACG |
| Hoxa13_B3 | TCCAAATCCCTGCCAGCTCCCAGAGTTCCACTCAACTTTAACCCg |
| Hoxa13_B3 | gTCCCTgCCTCTATATCTTTAAAACAAGGTGTTGGGGCAGAGGGG |
| Hoxa13_B3 | TAATAATGCCGAAATCCCCGCTATTTTCCACTCAACTTTAACCCg |
| Hoxa13_B3 | gTCCCTgCCTCTATATCTTTTTTGTAGGTTTTCTTGTGGAAGTAT |
| Hoxa13_B3 | ATTATATATTTATGAAATGCAGTTCTTCCACTCAACTTTAACCCg |
| Hoxa13_B3 | gTCCCTgCCTCTATATCTTTTTTGCATTAAGGTGGAACTTGTTTT |
| Hoxa13_B3 | ACTTATCTAGACAATTTTGTACCTATTCCACTCAACTTTAACCCg |
| Hoxa13_B3 | gTCCCTgCCTCTATATCTTTTTCCCGGAAACTACCAATTCTACAA |
| Hoxa13_B3 | TCTGTCACTCATTAAACTGAATTTTTTCCACTCAACTTTAACCCg |
| Hoxa13_B3 | gTCCCTgCCTCTATATCTTTCAGCATTGGGACGAGTGGCTCTTTT |
| Hoxa13_B3 | GGAGCACTGAAGCTGTCATAGCCCGTTCCACTCAACTTTAACCCg |
| **Rara** | |
| Rara_B1 | gAggAgggCAgCAAACggAACAGGCCTCCCAGCATGTGCGGAAAA |
| Rara_B1 | AATCCCTGCCAGAGATCCTGGTGGGTAgAAgAgTCTTCCTTTACg |
| Rara_B1 | gAggAgggCAgCAAACggAAGGCTTGTAGATCCGCGGCAAAGGGG |
| Rara_B1 | GAAGACTTGTCCTGGCACACAAAGCTAgAAgAgTCTTCCTTTACg |
| Rara_B1 | gAggAgggCAgCAAACggAAGTGTACACCATGTTCTTCTGAATGC |
| Rara_B1 | ATGATGCACGTCTTGTCGCGGTGGCTAgAAgAgTCTTCCTTTACg |
| Rara_B1 | gAggAgggCAgCAAACggAATGGCAGCGGTTCCGCGTCACCTTGT |
| Rara_B1 | TCGAAGCACTTCTGGAGCCGGCAGTTAgAAgAgTCTTCCTTTACg |
| Rara_B1 | gAggAgggCAgCAAACggAATTTTCCACGAGGTCCTCCACCTCTG |
| Rara_B1 | AAGGTTTCCTGATGGGCCTTGCGCATAgAAgAgTCTTCCTTTACg |
| Rara_B1 | gAggAgggCAgCAAACggAATCCCAGAGATCAATGTCCAGCGACA |
| Rara_B1 | CACTTGGTAGAGAGTTCACTGAACTTAgAAgAgTCTTCCTTTACg |
| Rara_B1 | gAggAgggCAgCAAACggAATTTGCGAACTCCACCGTCTTGATGA |
| Rara_B1 | GTGAGCGTGGTGAAGCCGGGCAGCTTAgAAgAgTCTTCCTTTACg |
| Rara_B1 | gAggAgggCAgCAAACggAATCAGGCGTGTACCTCGTGCAGATTC |
| Rara_B1 | CCGTCTGAGAACGTCATGGTGTCCTTAgAAgAgTCTTCCTTTACg |
| Rara_B1 | gAggAgggCAgCAAACggAATGCATCTGGGTTCGATTCAACGTCA |
| Rara_B1 | TCCGTCAGAGGTCCAAAGCCGGCATTAgAAgAgTCTTCCTTTACg |
| Rara_B1 | gAggAgggCAgCAAACggAAAGCTGATTGGCGAAGGCGAAGACCA |
| Rara_B1 | TCTGCGTCGTCCATCTCCAGCGGCATAgAAgAgTCTTCCTTTACg |
| Rara_B1 | gAggAgggCAgCAAACggAATGCAGCTTGTCCACCTTATCGGGCT |
| Rara_B1 | ATCTTTAACGCTTCGAGTAGGGGCTTAgAAgAgTCTTCCTTTACg |
| Rara_B1 | gAggAgggCAgCAAACggAATTGTTGGGTCTTCTTTTCCGGACGT |
| Rara_B1 | ATTAACATTTTGGGGAACATGTGGGTAgAAgAgTCTTCCTTTACg |
| Rara_B1 | gAggAgggCAgCAAACggAAATGGAGCCCGGGATCTCCATCTTGA |
| Rara_B1 | TCCAGCATTTCTTGGATGAGCGGCGTAgAAgAgTCTTCCTTTACg |
| Rara_B1 | gAggAgggCAgCAAACggAAGTCAGACTGTCCAGCCCCTCAGAGT |
| Rara_B1 | AGGCTGCTGGCGCGAGCAGGCTGTCTAgAAgAgTCTTCCTTTACg |
| Rara_B1 | gAggAgggCAgCAAACggAAAAGCTGGGGCTGCAGCTGCCTGGGG |
| Rara_B1 | GGACTGCTCCTGTTGGAGCTTGGCGTAgAAgAgTCTTCCTTTACg |
| Rara_B1 | gAggAgggCAgCAAACggAAACATGGTGTCAAGGGGAGTGACTGG |
| Rara_B1 | CTGCTGTCCTTGCGGTGGGAACCCCTAgAAgAgTCTTCCTTTACg |
| Rara_B1 | gAggAgggCAgCAAACggAAACACAGTCTAGACCGGTGTGCCCGG |
| Rara_B1 | TGGGAGAGTAGCGGAGGTCCGGCACTAgAAgAgTCTTCCTTTACg |
| Rara_B1 | gAggAgggCAgCAAACggAACCAATACCACCCTTGCCGCCTGTCC |
| Rara_B1 | CCGGACTCTCTCACCTCGCTGCACCTAgAAgAgTCTTCCTTTACg |
| Rara_B1 | gAggAgggCAgCAAACggAATCTGACTCCAGCTCCAGTTCCTTTT |
| Rara_B1 | CAAGTATCTTCTTTTCTGTGCTGCGTAgAAgAgTCTTCCTTTACg |
| Rara_B1 | gAggAgggCAgCAAACggAAAACGTTGCATGTGGTGCATTGAAAA |
| Rara_B1 | AACATTGTATGCATCTCGCAAGTATTAgAAgAgTCTTCCTTTACg |
| Rara_B1 | gAggAgggCAgCAAACggAATAAAGTCCAAAGCTGCACCCTGTAG |
| Rara_B1 | TGCATGTTCCAAGCTGCTGGCTGCCTAgAAgAgTCTTCCTTTACg |
| Rara_B1 | gAggAgggCAgCAAACggAAGCAGCAAATCCTCCTCCACCTCCCC |
| Rara_B1 | AAGCGCTATGCCCGTCCCAGCAACGTAgAAgAgTCTTCCTTTACg |
| Rara_B1 | gAggAgggCAgCAAACggAAGCCAGCAGGAGGCGCACTCATATTG |
| Rara_B1 | TGTCATCCTCTCCGCTCCTCCTGAATAgAAgAgTCTTCCTTTACg |
| Rara_B1 | gAggAgggCAgCAAACggAACTACCGTGGGTTGCAGCGGATTCAA |
| Rara_B1 | CCGGTGGGGCACAGTGGGTTGTCCTTAgAAgAgTCTTCCTTTACg |
| **Rarg** | |
| Rarg_B3 | gTCCCTgCCTCTATATCTTTTGGTGACCTCCTCCTTGCCACCTGG |
| Rarg_B3 | CCTTGCTAGAAGCCATGTTGGCCACTTCCACTCAACTTTAACCCg |
| Rarg_B3 | gTCCCTgCCTCTATATCTTTGCTGTGTGCCAGCGGTGCACATTCG |
| Rarg_B3 | AAGGAGGGAATCCATGCAGCTGGCTTTCCACTCAACTTTAACCCg |
| Rarg_B3 | gTCCCTgCCTCTATATCTTTTGCTGGAGAAGGCAAACGGATACAT |
| Rarg_B3 | GATCAAAAGGTGGGGAGCCACGCATTTCCACTCAACTTTAACCCg |
| Rarg_B3 | gTCCCTgCCTCTATATCTTTTGCGAAAGTAGGCACCGCCATTGGT |
| Rarg_B3 | TCTCCTTGGGTAGGTCTGTAGGGAATTCCACTCAACTTTAACCCg |
| Rarg_B3 | gTCCCTgCCTCTATATCTTTTTGTAGACTCGGGGCGGAGGCGGGG |
| Rarg_B3 | GACTTGTCATTGCACACGAAGCATGTTCCACTCAACTTTAACCCg |
| Rarg_B3 | gTCCCTgCCTCTATATCTTTCGATGGCACGTGTAGACCATGTTTT |
| Rarg_B3 | ACCTTGTTGATTTGGCAATTCTTGTTTCCACTCAACTTTAACCCg |
| Rarg_B3 | gTCCCTgCCTCTATATCTTTCGACAGTACTGGCATCGGTTCCGCG |
| Rarg_B3 | ATGCCAACTTCGAAGCATTTCTGCATTCCACTCAACTTTAACCCg |
| Rarg_B3 | gTCCCTgCCTCTATATCTTTTCCTCCTTGATCTCTTTCTTCTTTT |
| Rarg_B3 | GGCATCTCGTAGCTGTCGGTCACCATTCCACTCAACTTTAACCCg |
| Rarg_B3 | gTCCCTgCCTCTATATCTTTTTTGCGATCAGCGCCTCCATCTCGG |
| Rarg_B3 | AAGGTCTCCTGGTGTGCCTTGCTGATTCCACTCAACTTTAACCCg |
| Rarg_B3 | gTCCCTgCCTCTATATCTTTTCCCAGAGCCCAAGGTCGAGCTGGA |
| Rarg_B3 | CATTTGGTCGCCAGCTCGCTGAATTTTCCACTCAACTTTAACCCg |
| Rarg_B3 | gTCCCTgCCTCTATATCTTTTTGGCAAACTCTACAATCTTAATGA |
| Rarg_B3 | GTCAGAGTGGCAAAGCCGGGCAGCCTTCCACTCAACTTTAACCCg |
| Rarg_B3 | gTCCCTgCCTCTATATCTTTTCCGGCGTGTAACGGGTGCAAATCC |
| Rarg_B3 | CCATCTGAGAAGGTCATGGTGTCCTTTCCACTCAACTTTAACCCg |
| Rarg_B3 | gTCCCTgCCTCTATATCTTTTGCATCTGCGTCCGGTTCAGAGTGA |
| Rarg_B3 | TCCGTCAGGGGCCCAAAGCCTGCGTTTCCACTCAACTTTAACCCg |
| Rarg_B3 | gTCCCTgCCTCTATATCTTTAGCTGCTCAGCAAAGGCAAACACCA |
| Rarg_B3 | TCTGTGTCGTCCATCTCCAGAGGCATTCCACTCAACTTTAACCCg |
| Rarg_B3 | gTCCCTgCCTCTATATCTTTGTCTTCGTCGGCGGGCATAGATTTT |
| Rarg_B3 | TGCGTGGGAACATGTATGGCTTGTTTTCCACTCAACTTTAACCCg |
| Rarg_B3 | gTCCCTgCCTCTATATCTTTCTGGGATCTCCATCTTTAGTGTTAT |
| Rarg_B3 | TCTCACGTATGAGGGGTGGCATCGGTTCCACTCAACTTTAACCCg |
| Rarg_B3 | gTCCCTgCCTCTATATCTTTCTTCGAAGGCCTCTGGATTTTCAAG |
| Rarg_B3 | CGGCTTTGGGAGATGACAAAGCATCTTCCACTCAACTTTAACCCg |
| Rarg_B3 | gTCCCTgCCTCTATATCTTTTTTCTTCGACTTTGATGGGCTTCTC |
| Rarg_B3 | CTTTGGAACTAGGTTTCTCTGCGGGTTCCACTCAACTTTAACCCg |
| Rarg_B3 | gTCCCTgCCTCTATATCTTTCCCGGGATCCCCAGGCCCGCTAGAG |
| Rarg_B3 | GTTGGTGGAGACGGGCAGGAGCACTTTCCACTCAACTTTAACCCg |
| Rarg_B3 | gTCCCTgCCTCTATATCTTTGTGCCGCCACCTCTTGCTCAGAAAA |
| Rarg_B3 | TTGTCTGGGGTTCCCATAGCACCCTTTCCACTCAACTTTAACCCg |
| Rarg_B3 | gTCCCTgCCTCTATATCTTTCGAATCCAGATGACCCAGTATAAAA |
| Rarg_B3 | TTTAAGGCAAGAGAGATATATATACTTCCACTCAACTTTAACCCg |
| Rarg_B3 | gTCCCTgCCTCTATATCTTTTGGCAAAAGCGGCTGGGAGGTGGGG |
| Rarg_B3 | GTACTGCCAGAGGGTCCATCAATACTTCCACTCAACTTTAACCCg |
| Rarg_B3 | gTCCCTgCCTCTATATCTTTTTCCTTTCTCTTCAGATCCATCCCT |
| Rarg_B3 | CAAGTCCACAGGGGCACAGACCTTTTTCCACTCAACTTTAACCCg |
| Rarg_B3 | gTCCCTgCCTCTATATCTTTCTGGGTCGCTGAGGTGGAGAGGGCC |
| Rarg_B3 | CCCCTAAGGCATCCAGGAGCCTCTCTTCCACTCAACTTTAACCCg |
| Rarg_B3 | gTCCCTgCCTCTATATCTTTGGCATTGCACTGGGACAGTTCTGCC |
| Rarg_B3 | AACCTGTAACATTTTCTCCAATCAGTTCCACTCAACTTTAACCCg |
| **Raldh1** | |
| Raldh1_B1 | gAggAgggCAgCAAACggAACACACCTCCTGCTGCAACCCTTCAA |
| Raldh1_B1 | GGGGAAATGAGAGACATCGCTCGTGTAgAAgAgTCTTCCTTTACg |
| Raldh1_B1 | gAggAgggCAgCAAACggAAGATTGGTTGGCAACACCGGCGAGCC |
| Raldh1_B1 | TTAGACCACATTTGCATACACAGACTAgAAgAgTCTTCCTTTACg |
| Raldh1_B1 | gAggAgggCAgCAAACggAATTCCCTTTGCCTTTTACTCGCAGCC |
| Raldh1_B1 | ACTGTTTGGCCTGACTGAGCATGGATAgAAgAgTCTTCCTTTACg |
| Raldh1_B1 | gAggAgggCAgCAAACggAAAAAGCAAGGTGTTAGACAAGCCCTG |
| Raldh1_B1 | CACTGTTCTCTGGAACACAGGTGACTAgAAgAgTCTTCCTTTACg |
| Raldh1_B1 | gAggAgggCAgCAAACggAACCGTTCTGGTCCGGGCTTTGCTTTT |
| Raldh1_B1 | ATGGGTGCGGGCTGGGTTGCGGGAGTAgAAgAgTCTTCCTTTACg |
| Raldh1_B1 | gAggAgggCAgCAAACggAATTGGTGTATTTAATCTTCAGGTCCC |
| Raldh1_B1 | TCATGCCACTCGTTGTTTATAAATATAgAAgAgTCTTCCTTTACg |
| Raldh1_B1 | gAggAgggCAgCAAACggAAACTGAAAATTTCTTGCCGCTGGCAG |
| Raldh1_B1 | ATCTTGTCTCCAGTTGCAGGGTTGTTAgAAgAgTCTTCCTTTACg |
| Raldh1_B1 | gAggAgggCAgCAAACggAATCCTTGTCGCCTTCTTCGATTTCAC |
| Raldh1_B1 | GCAGCCTTGACAGCTTTGTCAACATTAgAAgAgTCTTCCTTTACg |
| Raldh1_B1 | gAggAgggCAgCAAACggAAGGGGAGCCAAGCTGAAACGCATCTC |
| Raldh1_B1 | CTTTCGGACGCGTCCATCCTTCGCCTAgAAgAgTCTTCCTTTACg |
| Raldh1_B1 | gAggAgggCAgCAAACggAATCTGCCAGTTTGTTTAAGATCCTCC |
| Raldh1_B1 | AAAATCAGACGATCCCTTTCGATTATAgAAgAgTCTTCCTTTACg |
| Raldh1_B1 | gAggAgggCAgCAAACggAACCAGCATCAAGGCATTCCAGAGTGC |
| Raldh1_B1 | GCATAGTAGGCAGCAGCAAAAGGTTTAgAAgAgTCTTCCTTTACg |
| Raldh1_B1 | gAggAgggCAgCAAACggAAAATGATTTAATGGCGCCTGGCAAAT |
| Raldh1_B1 | TTATCAGCCCAGCCCGCACAGTAGCTAgAAgAgTCTTCCTTTACg |
| Raldh1_B1 | gAggAgggCAgCAAACggAACGGCTCATGTCTTGTAAAAGTAAAA |
| Raldh1_B1 | GGGGATAATTTGTCCACACACACCATAgAAgAgTCTTCCTTTACg |
| Raldh1_B1 | gAggAgggCAgCAAACggAAAACCAACATAACCAAAGGGAAGTTC |
| Raldh1_B1 | ACAGCAGAGAGCAGGGCCAAGCTTCTAgAAgAgTCTTCCTTTACg |
| Raldh1_B1 | gAggAgggCAgCAAACggAATGCTGGTTTAATGACAACAGTATTT |
| Raldh1_B1 | GTAGACGGCAGAAAGGGGTGTTTGCTAgAAgAgTCTTCCTTTACg |
| Raldh1_B1 | gAggAgggCAgCAAACggAACCCTGCCTCAATGATCAATGATCCC |
| Raldh1_B1 | CACAATGTTGACTACGCCCGGTGGATAgAAgAgTCTTCCTTTACg |
| Raldh1_B1 | gAggAgggCAgCAAACggAAGGCTCCGGCTGTTGGCCCATAACCG |
| Raldh1_B1 | GTCAACGTCCATATGATGTGAAATGTAgAAgAgTCTTCCTTTACg |
| Raldh1_B1 | gAggAgggCAgCAAACggAACTCAGTTGACCCTGTGAAGGCAACT |
| Raldh1_B1 | AGCAGCTTCTTTGATGAGTTTGCCATAgAAgAgTCTTCCTTTACg |
| Raldh1_B1 | gAggAgggCAgCAAACggAAGGAGACCCTTTTCAGATTGCTCTTT |
| Raldh1_B1 | GATTGGGCTCTTTCCTCCCAGCTCCTAgAAgAgTCTTCCTTTACg |
| Raldh1_B1 | gAggAgggCAgCAAACggAAGTCCAAGTCAGCATCAGCAAATATG |
| Raldh1_B1 | ACCATTGTGAGCATGCTCGACTGCATAgAAgAgTCTTCCTTTACg |
| Raldh1_B1 | gAggAgggCAgCAAACggAAGCAGCATTGGCCCTGGTGGTAGAAC |
| Raldh1_B1 | CTCCACAAATATCCGGGAGCAGGCATAgAAgAgTCTTCCTTTACg |
| Raldh1_B1 | gAggAgggCAgCAAACggAAACGAACAAACTCTTCATAGATTGGC |
| Raldh1_B1 | CCTTGCCTTCGCTTTCACAATGCTCTAgAAgAgTCTTCCTTTACg |
| Raldh1_B1 | gAggAgggCAgCAAACggAATCCAGGAGTCACAGGGTCTCCTAGG |
| Raldh1_B1 | CTGGTCAATCTGAGGTCCCTGATTTTAgAAgAgTCTTCCTTTACg |
| Raldh1_B1 | gAggAgggCAgCAAACggAACATTCCAGCTTTGCTCCTTCCTTTT |
| Raldh1_B1 | CCTTTATTTCCCCAGGCCGCGCCTCTAgAAgAgTCTTCCTTTACg |
| Raldh1_B1 | gAggAgggCAgCAAACggAAGAGAAGACCGTGGGCTGTATAAAGA |
| Raldh1_B1 | GCGATGCGCATGTCATCCTTAACATTAgAAgAgTCTTCCTTTACg |
| Raldh1_B1 | gAggAgggCAgCAAACggAATGAACGGGTCCAAATATCTCTTCTT |
| Raldh1_B1 | TCTACTGTTTTGAATTTCATGATTTTAgAAgAgTCTTCCTTTACg |
| Raldh1_B1 | gAggAgggCAgCAAACggAAGTGTTGTTGGCTCGCTTAATAACTT |
| Raldh1_B1 | AAAACTCCTGCCGCTAAGCCATAGGTAgAAgAgTCTTCCTTTACg |
| Raldh1_B1 | gAggAgggCAgCAAACggAAGTCAGTGCCTTGTCGATGTCTTTGG |
| Raldh1_B1 | GTGCCCGCCTGGAGAGCAGTAGCAATAgAAgAgTCTTCCTTTACg |
| Raldh1_B1 | gAggAgggCAgCAAACggAAATGGCGCTGTAGCAATTAACCCATA |
| Raldh1_B1 | AATCCTCCAAAAGGACACTGTGAGGTAgAAgAgTCTTCCTTTACg |
| Raldh1_B1 | gAggAgggCAgCAAACggAAATTTCTCTCCCGTTGCCTGACATTT |
| Raldh1_B1 | GTATATTCTGAAATGCCATATTCTCTAgAAgAgTCTTCCTTTACg |
| Raldh1_B1 | gAggAgggCAgCAAACggAAATTTTGATTGTGACAGTCTTAACTT |
| Raldh1_B1 | AGGAACGTTTAGGAATTCTTCTGTGTAgAAgAgTCTTCCTTTACg |
| Raldh1_B1 | gAggAgggCAgCAAACggAATTAATGGAGTCAGGTTCATTTCAAA |
| Raldh1_B1 | AAAGGCCCATGCTTGATGTTTGTTATAgAAgAgTCTTCCTTTACg |
| Raldh1_B1 | gAggAgggCAgCAAACggAAGCACATTTCTAACAACTTTGCTTTT |
| Raldh1_B1 | TTTATTGTAAGCATACGTAAAGTTATAgAAgAgTCTTCCTTTACg |
| **Raldh2** | |
| Raldh2_B2 | CCTCgTAAATCCTCATCAAAGGGTCCGGCTTCACCTCCCCGGGCA |
| Raldh2_B2 | AGGTGCAGGGAGGCCATGAGAGCCGAAATCATCCAgTAAACCgCC |
| Raldh2_B2 | CCTCgTAAATCCTCATCAAAGTTCTGCCACTCGTTGTTGATAAAA |
| Raldh2_B2 | CACTGGGAATACTTTGCCGCTCACGAAATCATCCAgTAAACCgCC |
| Raldh2_B2 | CCTCgTAAATCCTCATCAAAAATCATCTCTGACGTGGCTGGATTG |
| Raldh2_B2 | AGCCTTGTCAGCTTCCTGGACTTCAAAATCATCCAgTAAACCgCC |
| Raldh2_B2 | CCTCgTAAATCCTCATCAAACGCAGCTCGCACTGCTTTGTCTATA |
| Raldh2_B2 | GACGGAGCCCATGGAGAAGGCAAGTAAATCATCCAgTAAACCgCC |
| Raldh2_B2 | CCTCgTAAATCCTCATCAAACCTCTCCGATGCGTCCATTCTCCGC |
| Raldh2_B2 | GTCAGCCAGTTTGCTCAAGAGTCGGAAATCATCCAgTAAACCgCC |
| Raldh2_B2 | CCTCgTAAATCCTCATCAAAAAGAAGGGCCAAGTCCCTTTCTACG |
| Raldh2_B2 | GCCGCTGTCCAGAGATTCAATGGTAAAATCATCCAgTAAACCgCC |
| Raldh2_B2 | CCTCgTAAATCCTCATCAAAAATATCTCAGTGTCTTTATCGCCCC |
| Raldh2_B2 | CATGAATCTTGTCGGCCCAGCCTGCAAATCATCCAgTAAACCgCC |
| Raldh2_B2 | CCTCgTAAATCCTCATCAAAAATCGCCATCTGCTGGGATGGTTAT |
| Raldh2_B2 | TGGGCTCGTGCCTCGTGAACGTGAAAAATCATCCAgTAAACCgCC |
| Raldh2_B2 | CCTCgTAAATCCTCATCAAATCTTCCAGGCAAACATCAGCAGGGG |
| Raldh2_B2 | TGTTACCACAGCACAGAGCTGGAGCAAATCATCCAgTAAACCgCC |
| Raldh2_B2 | CCTCgTAAATCCTCATCAAATTTGTTCGGCCGGTTTAATAACCAC |
| Raldh2_B2 | ATCCCATGTGGAGCGCGCTGAGCGGAAATCATCCAgTAAACCgCC |
| Raldh2_B2 | CCTCgTAAATCCTCATCAAAGCGGAAACCCAGCCTCTTTGATGAG |
| Raldh2_B2 | ATCCCGGCAAAATATTGACGACTCCAAATCATCCAgTAAACCgCC |
| Raldh2_B2 | CCTCgTAAATCCTCATCAAACCATTGCTGCGCCTGCCGTCGGACC |
| Raldh2_B2 | CTACTTTATCGATGCCGGAGTGAGAAAATCATCCAgTAAACCgCC |
| Raldh2_B2 | CCTCgTAAATCCTCATCAAATCCCAACTTCAGTGGATCCAGTAAA |
| Raldh2_B2 | TCCTTCCAGCTGCCTCCTGGATCAGAAATCATCCAgTAAACCgCC |
| Raldh2_B2 | CCTCgTAAATCCTCATCAAAGCTCCAGGGTAACTCTTTTCAAATT |
| Raldh2_B2 | AAATAATGTTGGGGCTCTTCCCACCAAATCATCCAgTAAACCgCC |
| Raldh2_B2 | CCTCgTAAATCCTCATCAAACTGCATAGTCCAAATCGGCATCTGC |
| Raldh2_B2 | AGAAAACGCCTTGGTGGGCTTGCTCAAATCATCCAgTAAACCgCC |
| Raldh2_B2 | CCTCgTAAATCCTCATCAAACTGCAGTGCAACACTGCCCTTGATT |
| Raldh2_B2 | TGGTCTCCTCAACAAATGTCCGGGAAAATCATCCAgTAAACCgCC |
| Raldh2_B2 | CCTCgTAAATCCTCATCAAACACTCCGCCGGACAAACTCATCGTA |
| Raldh2_B2 | CTACTATACGCCTCTTGGCACGCTCAAATCATCCAgTAAACCgCC |
| Raldh2_B2 | CCTCgTAAATCCTCATCAAAGTTCAGTGGAAGGATCAAAGGGACT |
| Raldh2_B2 | ACTGTTTCATATCCGTCTGTGGACCAAATCATCCAgTAAACCgCC |
| Raldh2_B2 | CCTCgTAAATCCTCATCAAATTTGGATGAGATCCAGAATTTTGTT |
| Raldh2_B2 | CTAGTTTAGCGCCTTCCATGACACCAAATCATCCAgTAAACCgCC |
| Raldh2_B2 | CCTCgTAAATCCTCATCAAACCCTTTCTTCCTAAGCCTTTGCCCC |
| Raldh2_B2 | GAAAACACAGTTGGTTCCACGAATAAAATCATCCAgTAAACCgCC |
| Raldh2_B2 | CCTCgTAAATCCTCATCAAAGAGTATTTGTTGGACGGGTCCAAAA |
| Raldh2_B2 | GATCACTTCTTCCATGGTTTTAAACAAATCATCCAgTAAACCgCC |
| Raldh2_B2 | CCTCgTAAATCCTCATCAAATCCAAACTCCGAGCTGTTCGCTCTC |
| Raldh2_B2 | GTCATTTGTGAAAACGGCTGCTACCAAATCATCCAgTAAACCgCC |
| Raldh2_B2 | CCTCgTAAATCCTCATCAAACGAGGAAACCGTCAGGGCTTTGCTG |
| Raldh2_B2 | AATCCAGACTGTTCCAGCTTGCATTAAATCATCCAgTAAACCgCC |
| Raldh2_B2 | CCTCgTAAATCCTCATCAAACTGGGCATTCAAGGCATTGTAGCAA |
| Raldh2_B2 | GGACATCTTATATCCTCCGAAAGGAAAATCATCCAgTAAACCgCC |
| Raldh2_B2 | CCTCgTAAATCCTCATCAAAGTACTCGCCCATTTCCCTCCCATTA |
| Raldh2_B2 | CTTGGCTTCTGTGTACTCCCGCAATAAATCATCCAgTAAACCgCC |
| Raldh2_B2 | CCTCgTAAATCCTCATCAAAAGCGCTCTTAGGAGTTCTTTTGGGG |
| Raldh2_B2 | GCGTGCTCTTCGCCCCACGTGGTCTAAATCATCCAgTAAACCgCC |
| Raldh2_B2 | CCTCgTAAATCCTCATCAAAGTTGCTGTAACGAATACTGATGGGG |
| Raldh2_B2 | CTGTATAACACTGCATTCCTGAGTGAAATCATCCAgTAAACCgCC |
| Raldh2_B2 | CCTCgTAAATCCTCATCAAATTTGCCAACTAGGATCTGAAATATA |
| Raldh2_B2 | ATTTACGGCAAGGCATCTGGTTCACAAATCATCCAgTAAACCgCC |
| Raldh2_B2 | CCTCgTAAATCCTCATCAAAGGACACCATGCGGAAACCCTTGATA |
| Raldh2_B2 | CGTACATTGTGTAACGGTGGGGTGGAAATCATCCAgTAAACCgCC |
| Raldh2_B2 | CCTCgTAAATCCTCATCAAAGTAAGACTTAGCATGTTATTTTGTG |
| Raldh2_B2 | TGTTGAACGTTCATTAAATGGTTTAAAATCATCCAgTAAACCgCC |
| Raldh2_B2 | CCTCgTAAATCCTCATCAAAGCACTGTATATGTTTAATAGCTTGA |
| Raldh2_B2 | TCTAGCATCCTTCAACTTACTGCACAAATCATCCAgTAAACCgCC |
| Raldh2_B2 | CCTCgTAAATCCTCATCAAATGGGAAACAGACCCCAACCTGCAGT |
| Raldh2_B2 | CAAAGTATACAGGGCTGTTCATCCAAAATCATCCAgTAAACCgCC |
| Raldh2_B2 | CCTCgTAAATCCTCATCAAATAGCGTGTACACGCACTCTTGTTTT |
| Raldh2_B2 | AGGGGCGTCGAAACACTGATGATATAAATCATCCAgTAAACCgCC |
| Raldh2_B2 | CCTCgTAAATCCTCATCAAAACTCTCAGTTTCCTGAAGAAGCATT |
| Raldh2_B2 | GTCAGAGCAGAGAAGCAATTTAATTAAATCATCCAgTAAACCgCC |
| Raldh2_B2 | CCTCgTAAATCCTCATCAAAAAGCACAACAGTCTGGATTGGTTGA |
| Raldh2_B2 | CATAAATATCCTAGGTTATGGTGACAAATCATCCAgTAAACCgCC |
| Raldh2_B2 | CCTCgTAAATCCTCATCAAATAAAGACAAAGTGGGTTTATATTTT |
| Raldh2_B2 | TAGACACCAAAGTTTATCATACAGAAAATCATCCAgTAAACCgCC |
| Raldh2_B2 | CCTCgTAAATCCTCATCAAACATGTTGACATGACAGTGTTATCTA |
| Raldh2_B2 | GATCAGCATCTCCTGCATGATGTGGAAATCATCCAgTAAACCgCC |
| **Raldh3** | |
| Raldh3_B3 | gTCCCTgCCTCTATATCTTTTTTCTGATGGGATGGGGCAGCGGGG |
| Raldh3_B3 | AATATCTTGCTGTATTTGACGGGCATTCCACTCAACTTTAACCCg |
| Raldh3_B3 | gTCCCTgCCTCTATATCTTTTTGGATTCGTGCCATTCATTGTTAA |
| Raldh3_B3 | TTGTGTGTCGCAAATTTCTTCCCGCTTCCACTCAACTTTAACCCg |
| Raldh3_B3 | gTCCCTgCCTCTATATCTTTTCGCAGATTTTCTCATTGGTGGAAG |
| Raldh3_B3 | ACATCTGGCTTGTCTCCTTCCTCCATTCCACTCAACTTTAACCCg |
| Raldh3_B3 | gTCCCTgCCTCTATATCTTTGCTCGTGCAGCTTCCACTGCCCTAT |
| Raldh3_B3 | CTCCACGGTGAGCCCTTCTCAAAGGTTCCACTCAACTTTAACCCg |
| Raldh3_B3 | gTCCCTgCCTCTATATCTTTCGGCCTCGGCTCAGAGCATCCATCT |
| Raldh3_B3 | ATGAGGTCAGCCAGCTTGTGGAGCATTCCACTCAACTTTAACCCg |
| Raldh3_B3 | gTCCCTgCCTCTATATCTTTGTCGCCAGGATGACTCGATCACGCT |
| Raldh3_B3 | GGTTTTCCTGTGTCCATTGTTTCCATTCCACTCAACTTTAACCCg |
| Raldh3_B3 | gTCCCTgCCTCTATATCTTTAGGTCAATCAGGAAGGCATGGAGGA |
| Raldh3_B3 | TATCGCAGGGTCTTTATGCAGCCCTTTCCACTCAACTTTAACCCg |
| Raldh3_B3 | gTCCCTgCCTCTATATCTTTTGTATTTTATCGGCCCAGCCTGCGT |
| Raldh3_B3 | CTGTCATCAATGGGGATGGTCCGGCTTCCACTCAACTTTAACCCg |
| Raldh3_B3 | gTCCCTgCCTCTATATCTTTGGCTCATGCATAGTGAAGCAGACAA |
| Raldh3_B3 | GGAGTTATAGCCCCGCAGACACCGATTCCACTCAACTTTAACCCg |
| Raldh3_B3 | gTCCCTgCCTCTATATCTTTACGAGCATCAGTAACGGGAAGTTCC |
| Raldh3_B3 | CAGCACAGCGCGGGCGCCATTTTCCTTCCACTCAACTTTAACCCg |
| Raldh3_B3 | gTCCCTgCCTCTATATCTTTGCGGGTTTGATAACCAGCGTGTTGC |
| Raldh3_B3 | TAGAGTGACGTCAGTGGTGTCTGCTTTCCACTCAACTTTAACCCg |
| Raldh3_B3 | gTCCCTgCCTCTATATCTTTCCCACCTCCTTGATTAATGAGCCAA |
| Raldh3_B3 | ACTATGTTCACTACACCAGGAGGAATTCCACTCAACTTTAACCCg |
| Raldh3_B3 | gTCCCTgCCTCTATATCTTTGCCCCAGCCTTTGGTCCGTAGCCGG |
| Raldh3_B3 | TCAATATCTGGATGGCTCGAGATGGTTCCACTCAACTTTAACCCg |
| Raldh3_B3 | gTCCCTgCCTCTATATCTTTTCTGTGGATCCAGTGAAGGCCACTT |
| Raldh3_B3 | GCAGCTTCCTTGATGAGTTTACCAATTCCACTCAACTTTAACCCg |
| Raldh3_B3 | gTCCCTgCCTCTATATCTTTTCAGCAAACACAATGCAGGGATTTT |
| Raldh3_B3 | CACTCCACGGCCAGCTCCAGATCACTTCCACTCAACTTTAACCCg |
| Raldh3_B3 | gTCCCTgCCTCTATATCTTTTGGTTGAAGAAGGCGCCCTGGTGAG |
| Raldh3_B3 | CTGGAGGCCGCGGTGCAGCACTGCCTTCCACTCAACTTTAACCCg |
| Raldh3_B3 | gTCCCTgCCTCTATATCTTTTGGTAGATCTTCTCCTCGACAAAGA |
| Raldh3_B3 | CAGTCGACGCTCCGTCGGACAAATTTTCCACTCAACTTTAACCCg |
| Raldh3_B3 | gTCCCTgCCTCTATATCTTTGGGTCTCCGAGAAGTCTCTTCTTGG |
| Raldh3_B3 | GGGCCTTGTTCAGTTTTGGCATCCATTCCACTCAACTTTAACCCg |
| Raldh3_B3 | gTCCCTgCCTCTATATCTTTTTATCAAATTGGGCTCTGTCGATCT |
| Raldh3_B3 | TTCCCACTTTCTATGAGCCCCAAAATTCCACTCAACTTTAACCCg |
| Raldh3_B3 | gTCCCTgCCTCTATATCTTTGGTGGGTTTGATGAACAGTCCTTTT |
| Raldh3_B3 | CATGCCGTCCGTCACATCGGAAAACTTCCACTCAACTTTAACCCg |
| Raldh3_B3 | gTCCCTgCCTCTATATCTTTTCCAAATATCTCCTCCTTGGCGATC |
| Raldh3_B3 | CTTGAACTTCATTATCGACTGCACTTTCCACTCAACTTTAACCCg |
| Raldh3_B3 | gTCCCTgCCTCTATATCTTTGGCTCGCTTAATGACGTCTTCTATA |
| Raldh3_B3 | GGCAGTGAGTCCATACTGCGTGTTGTTCCACTCAACTTTAACCCg |
| Raldh3_B3 | gTCCCTgCCTCTATATCTTTTTTATCCAGACTTTTGGTGAACACT |
| Raldh3_B3 | CTGCATGGAGGAGGCGATGGTCATGTTCCACTCAACTTTAACCCg |
| Raldh3_B3 | gTCCCTgCCTCTATATCTTTGTAACAGTTGATCCAGACTGTGCCA |
| Raldh3_B3 | AAACGGAGCCTGGGCATGAAGTGCATTCCACTCAACTTTAACCCg |
| Raldh3_B3 | gTCCCTgCCTCTATATCTTTTCCGTTTCCCGACATTTTAAACCCT |
| Raldh3_B3 | CGCCAGACTGTATTCACCTAGTTCTTTCCACTCAACTTTAACCCg |
| Raldh3_B3 | gTCCCTgCCTCTATATCTTTATTTATTTGTATTGTGTTGGCTTTT |
| Raldh3_B3 | TGTCTCTTTCACATTCAGAATGTCATTCCACTCAACTTTAACCCg |
| Raldh3_B3 | gTCCCTgCCTCTATATCTTTCAGAAGGTTATAATTTACAATATTT |
| Raldh3_B3 | TCATTGAGCACATTGCAGGTATAAATTCCACTCAACTTTAACCCg |
| Raldh3_B3 | gTCCCTgCCTCTATATCTTTCCAACATGAGGAAGAGCTATGCCAG |
| Raldh3_B3 | CACAGCAGAATATGGTGGAATAGAATTCCACTCAACTTTAACCCg |
| Raldh3_B3 | gTCCCTgCCTCTATATCTTTGGGATTTCTCTTCTCCGAAACAAAA |
| Raldh3_B3 | ATGCGTCATTTCATGTTTTCATAGTTTCCACTCAACTTTAACCCg |
| Raldh3_B3 | gTCCCTgCCTCTATATCTTTCAGCAAAATGTAGTTAATTATTTAT |
| Raldh3_B3 | TGCATTGACGTAACACATCCCTGTCTTCCACTCAACTTTAACCCg |
| Raldh3_B3 | gTCCCTgCCTCTATATCTTTACCAGTTGGCCACCACACGGTGATC |
| Raldh3_B3 | GTTATTTTATTGTCTCTGTTCTTAATTCCACTCAACTTTAACCCg |
| Raldh3_B3 | gTCCCTgCCTCTATATCTTTATTAATCTGAAGATGCCCCTGTTTT |
| Raldh3_B3 | TCAATGGAAGACCACATTCCCTTCATTCCACTCAACTTTAACCCg |
| Raldh3_B3 | gTCCCTgCCTCTATATCTTTTAAAACTAACTTCCACATGGTCTCG |
| Raldh3_B3 | TCCTTCTCATGCTATGTGGCAAAGCTTCCACTCAACTTTAACCCg |
| Raldh3_B3 | gTCCCTgCCTCTATATCTTTGAAACAAAATAAGAACAAAAGATAA |
| Raldh3_B3 | CGCGCTCTATAAGCATACACACTAATTCCACTCAACTTTAACCCg |
| Raldh3_B3 | gTCCCTgCCTCTATATCTTTCAACTGGGAGAGCTCAATGACAAAC |
| Raldh3_B3 | AGACAATCAACTTAATTTTCTTTGTTTCCACTCAACTTTAACCCg |
| Raldh3_B3 | gTCCCTgCCTCTATATCTTTAAGACGTATAAGATGTGTGACTCTG |
| Raldh3_B3 | GGACCACACTAATACCTTTTCCAATTTCCACTCAACTTTAACCCg |
| Raldh3_B3 | gTCCCTgCCTCTATATCTTTAAGATGGTGTACTACATTATCTTTC |
| Raldh3_B3 | TTTTGTTGGACTCACAAAAGAAATGTTCCACTCAACTTTAACCCg |
| **Cyp26a1** | |
| Cyp26a1_B2 | CCTCgTAAATCCTCATCAAATATACATTTCCCCAAGCCTAGCCCC |
| Cyp26a1_B2 | ACTGGGAAGCATTGCCTCTCCTCCTAAATCATCCAgTAAACCgCC |
| Cyp26a1_B2 | CCTCgTAAATCCTCATCAAATCTGAGGAGCCCACTGTCCTAGTGT |
| Cyp26a1_B2 | AGCTCTCTACCCACGCAGCTCTCCTAAATCATCCAgTAAACCgCC |
| Cyp26a1_B2 | CCTCgTAAATCCTCATCAAACCCTGCCTGCAGATCTCTCTCCTAG |
| Cyp26a1_B2 | GCTCCTGCAGCCCTGTTAGCACTATAAATCATCCAgTAAACCgCC |
| Cyp26a1_B2 | CCTCgTAAATCCTCATCAAACTATTAGCACTTTCTCTTTGTACTC |
| Cyp26a1_B2 | GCAGTATCTCGTGTGTGGACCGCGCAAATCATCCAgTAAACCgCC |
| Cyp26a1_B2 | CCTCgTAAATCCTCATCAAACCCCTACCCTACCAGCACCCTCTCT |
| Cyp26a1_B2 | CCCTGCTGCAGGTCTCTTCCCCGGTAAATCATCCAgTAAACCgCC |
| Cyp26a1_B2 | CCTCgTAAATCCTCATCAAAACTAGGCTGGCTCCACCACAACTCT |
| Cyp26a1_B2 | TGCACACTACCTGCAGCGTTGTCAGAAATCATCCAgTAAACCgCC |
| Cyp26a1_B2 | CCTCgTAAATCCTCATCAAACACCGGCACAGCGCGTTCCTCGGAC |
| Cyp26a1_B2 | CCCCACAGTAGCCGCGGCAGCAGTCAAATCATCCAgTAAACCgCC |
| Cyp26a1_B2 | CCTCgTAAATCCTCATCAAAGCACAGCTACTCCCTCTGCCCGGGG |
| Cyp26a1_B2 | CTGGCGAGCAGGGTGGAGAAGCCCAAAATCATCCAgTAAACCgCC |
| Cyp26a1_B2 | CCTCgTAAATCCTCATCAAAAACGGCAGTATCAAGGTGCACAGCG |
| Cyp26a1_B2 | AGCTTCACTGCGGCCAAGAAGAGCAAAATCATCCAgTAAACCgCC |
| Cyp26a1_B2 | CCTCgTAAATCCTCATCAAACGGGTGCTCACGCAGTACAGGTCCC |
| Cyp26a1_B2 | GGCAGCGGGCACCGGGAGAGGCGGTAAATCATCCAgTAAACCgCC |
| Cyp26a1_B2 | CCTCgTAAATCCTCATCAAAAAGAACGGGAGCCCCATGGTTCCCG |
| Cyp26a1_B2 | TGTAGCACCAATTGCAGGGTCTCCCAAATCATCCAgTAAACCgCC |
| Cyp26a1_B2 | CCTCgTAAATCCTCATCAAACGTTTCATCTGTAGGAATTTTCGTC |
| Cyp26a1_B2 | GTCTTGTAAATAAACCCGTACTTCCAAATCATCCAgTAAACCgCC |
| Cyp26a1_B2 | CCTCgTAAATCCTCATCAAACGCACCGTGGGTCTGCCGAACAGGT |
| Cyp26a1_B2 | TGCTTCACGTTCTCCGCGCCCATCAAAATCATCCAgTAAACCgCC |
| Cyp26a1_B2 | CCTCgTAAATCCTCATCAAAGCACAGACACGAGCCTGTGTTCCCC |
| Cyp26a1_B2 | AGATGGTCCGCACCGAGGCGGGCCAAAATCATCCAgTAAACCgCC |
| Cyp26a1_B2 | CCTCgTAAATCCTCATCAAAGCAGGTTGGACAGGCAGCCGGCCCC |
| Cyp26a1_B2 | TTTTGCGGTTCTTGTGTAGGGAGTCAAATCATCCAgTAAACCgCC |
| Cyp26a1_B2 | CCTCgTAAATCCTCATCAAACCCGAGAAAACGCCTGCATGATAAC |
| Cyp26a1_B2 | TCACAGGGATGTAGTGCTGCAGCGCAAATCATCCAgTAAACCgCC |
| Cyp26a1_B2 | CCTCgTAAATCCTCATCAAACTATCGAGCCGCGCACCTCTTCCTC |
| Cyp26a1_B2 | ACGATGCGCCCCTTGCAAGCCACTGAAATCATCCAgTAAACCgCC |
| Cyp26a1_B2 | CCTCgTAAATCCTCATCAAAGCTTCACCTCAGGGTAGACCAGCAC |
| Cyp26a1_B2 | TCCTCATGGCGATGCGGAACATGAGAAATCATCCAgTAAACCgCC |
| Cyp26a1_B2 | CCTCgTAAATCCTCATCAAATCTGGTGCGGCTCGAAGCCCAGCAA |
| Cyp26a1_B2 | CTAGCTGCTGTTCAGTCTCTGGGTCAAATCATCCAgTAAACCgCC |
| Cyp26a1_B2 | CCTCgTAAATCCTCATCAAATGCGGGTCATTTCCTCGAAGGCTTC |
| Cyp26a1_B2 | GCACATCGATGGGCAGCGAAAAGAGAAATCATCCAgTAAACCgCC |
| Cyp26a1_B2 | CCTCgTAAATCCTCATCAAAGATGACATTTCTAGCCTTCAGCCCC |
| Cyp26a1_B2 | CTTAATGTTCTCTTCGATTTTAGAGAAATCATCCAgTAAACCgCC |
| Cyp26a1_B2 | CCTCgTAAATCCTCATCAAACGTGTTGGAGTCCTTGGCCATTTTC |
| Cyp26a1_B2 | GAGTAATTGCAGCGCGTCCTTGTACAAATCATCCAgTAAACCgCC |
| Cyp26a1_B2 | CCTCgTAAATCCTCATCAAACTCGCCATTCTTCTGCGTGTGCTCG |
| Cyp26a1_B2 | CTCCTTCAGCTCCTGCATGTTCAGTAAATCATCCAgTAAACCgCC |
| Cyp26a1_B2 | CCTCgTAAATCCTCATCAAAGCCTCCAAAGAGGAGCTCGGTGGCA |
| Cyp26a1_B2 | GGTGGCGGCGCTGGCTGTGGTCTCGAAATCATCCAgTAAACCgCC |
| Cyp26a1_B2 | CCTCgTAAATCCTCATCAAAGTGCAGCGCCAAGAAAGTCATAAGC |
| Cyp26a1_B2 | CTTTCTGACTTTGTGCAGCACGTCAAAATCATCCAgTAAACCgCC |
| Cyp26a1_B2 | CCTCgTAAATCCTCATCAAAGGACAATAAATCCTTGATTTGAAGT |
| Cyp26a1_B2 | CGGCTTGTTCTCCTGGCTGCTGCTGAAATCATCCAgTAAACCgCC |
| Cyp26a1_B2 | CCTCgTAAATCCTCATCAAAGAGTTGCTCCAGGGCTTCCACGGAC |
| Cyp26a1_B2 | CTCCTTAATGACGCAGGCCGTGTACAAATCATCCAgTAAACCgCC |
| Cyp26a1_B2 | CCTCgTAAATCCTCATCAAAAGGCACGGGCGGGCTGAGCCGCAGA |
| Cyp26a1_B2 | GGTTTTGAGCGCCACGCGGAACCCCAAATCATCCAgTAAACCgCC |
| Cyp26a1_B2 | CCTCgTAAATCCTCATCAAAGGGAATCTGATATCCATTCAACTCG |
| Cyp26a1_B2 | GATGCTGTAGATGACATTCCAGCCTAAATCATCCAgTAAACCgCC |
| Cyp26a1_B2 | CCTCgTAAATCCTCATCAAAGACGTCGGCCACGTCGTGGGTGTCG |
| Cyp26a1_B2 | CGGATTGAACTCCTCCTTGTCTGTGAAATCATCCAgTAAACCgCC |
| Cyp26a1_B2 | CCTCgTAAATCCTCATCAAACTCCGGGTGGGAGGACATGAACCTG |
| Cyp26a1_B2 | GGGAATGAAGTTGAAGCGGGCGTGGAAATCATCCAgTAAACCgCC |
| Cyp26a1_B2 | CCTCgTAAATCCTCATCAAACCCACACAGCTCCTCAGGCCGCCCC |
| Cyp26a1_B2 | TTCAGGAGGAGCTTGGCGAACTCCTAAATCATCCAgTAAACCgCC |
| Cyp26a1_B2 | CCTCgTAAATCCTCATCAAAGTCCGTGCCAGCTCCACGATGAAGA |
| Cyp26a1_B2 | GCCCCGTTGAGGAGATGCCAGTCGCAAATCATCCAgTAAACCgCC |
| Cyp26a1_B2 | CCTCgTAAATCCTCATCAAAACAGTGGGCCCAGTCTTCATGGTGG |
| Cyp26a1_B2 | TTGGCGGGGAGATTGTCCACGGGGTAAATCATCCAgTAAACCgCC |
| Cyp26a1_B2 | CCTCgTAAATCCTCATCAAATACATTATCCCGTTGAAACGTATAA |
| Cyp26a1_B2 | GGGTTTACGTGTAGCCCTGGCAGCGAAATCATCCAgTAAACCgCC |
| Cyp26a1_B2 | CCTCgTAAATCCTCATCAAACACATATACATATGTTACTATAAAA |
| Cyp26a1_B2 | AGTTAAATACAGAATAGGTAAAATTAAATCATCCAgTAAACCgCC |
| Cyp26a1_B2 | CCTCgTAAATCCTCATCAAAAACGCAGAGACAAGAGTTTACAAAA |
| Cyp26a1_B2 | AACAGACCCATAATGTTAGTGCTTGAAATCATCCAgTAAACCgCC |
| **Cyp26b1** | |
| Cyp26b1_B1 | gAggAgggCAgCAAACggAAAGGACCACTGACACTAGGCAGGCGG |
| Cyp26b1_B1 | CACAGCTGTTGGGAGACGGCCAGCATAgAAgAgTCTTCCTTTACg |
| Cyp26b1_B1 | gAggAgggCAgCAAACggAATCCCGGGTGGCAGCCCAGCGGAGTT |
| Cyp26b1_B1 | TTGGGGATGGGCAGCTTGCAGCTCTTAgAAgAgTCTTCCTTTACg |
| Cyp26b1_B1 | gAggAgggCAgCAAACggAACGTTGCCATATTTCTCCCTTCTGGA |
| Cyp26b1_B1 | GCCGCCCCAACAAGTGTGTCTTGAATAgAAgAgTCTTCCTTTACg |
| Cyp26b1_B1 | gAggAgggCAgCAAACggAATCTCTGCCCCGGTCACTCTGATTAA |
| Cyp26b1_B1 | GCTCACCCATCAGAATCTTGCGGACTAgAAgAgTCTTCCTTTACg |
| Cyp26b1_B1 | gAggAgggCAgCAAACggAAGAGGCCACTCTGTGCTCACCAGGCT |
| Cyp26b1_B1 | TGGGGCCTAGTAGGGTCCTTGTGCTTAgAAgAgTCTTCCTTTACg |
| Cyp26b1_B1 | gAggAgggCAgCAAACggAACCAGGGCCTCGTGGCTGAAGATCTT |
| Cyp26b1_B1 | CTAGCTGGATCTTGGGAAGGTAACTTAgAAgAgTCTTCCTTTACg |
| Cyp26b1_B1 | gAggAgggCAgCAAACggAATCCACATCCTCAATGTGTCCTGGAT |
| Cyp26b1_B1 | AGACATTAATGGGATCAGGGTTACTTAgAAgAgTCTTCCTTTACg |
| Cyp26b1_B1 | gAggAgggCAgCAAACggAAGGAAGGTTAACTTCTGGGCCTCGAA |
| Cyp26b1_B1 | ATCCCAGAAGGACCCGGATGGCCATTAgAAgAgTCTTCCTTTACg |
| Cyp26b1_B1 | gAggAgggCAgCAAACggAAGATTGAGTTCCTCATCAGAGAGGCG |
| Cyp26b1_B1 | CAAACTGCTGAAAGACCTGGAAGAGTAgAAgAgTCTTCCTTTACg |
| Cyp26b1_B1 | gAggAgggCAgCAAACggAACCACAGGCAGGGAGAACACGTTCTC |
| Cyp26b1_B1 | CCCTTCTGTAGCCACTGAATGGCATTAgAAgAgTCTTCCTTTACg |
| Cyp26b1_B1 | gAggAgggCAgCAAACggAATCTGCAGGGTCTCGCGAGCCCGAAT |
| Cyp26b1_B1 | TCTCTCGAATGGCCTTCTCCAGACTTAgAAgAgTCTTCCTTTACg |
| Cyp26b1_B1 | gAggAgggCAgCAAACggAAAGTCCTTCCCCTGCGAATTCTGGAA |
| Cyp26b1_B1 | CAATCAAAATATCGAGCGCGTCTGCTAgAAgAgTCTTCCTTTACg |
| Cyp26b1_B1 | gAggAgggCAgCAAACggAAGGCTGTTGTAGCATAGGCAGCAAAA |
| Cyp26b1_B1 | CTGCATTATGAGTGAGGTGCTGGAGTAgAAgAgTCTTCCTTTACg |
| Cyp26b1_B1 | gAggAgggCAgCAAACggAATTCGAAGACAGCAGGATGCTTCAAG |
| Cyp26b1_B1 | GTTGCCCCGGAGTTCCTCCCGCAGCTAgAAgAgTCTTCCTTTACg |
| Cyp26b1_B1 | gAggAgggCAgCAAACggAAACAGATGCATCCATTGTGGAGGATA |
| Cyp26b1_B1 | GATGTTGTCTACCCGGAACGCTCCGTAgAAgAgTCTTCCTTTACg |
| Cyp26b1_B1 | gAggAgggCAgCAAACggAACACACAGTCCAAGTAATGGAGGCTG |
| Cyp26b1_B1 | ACTAAAGAGCCGGAGGACTTCCTTGTAgAAgAgTCTTCCTTTACg |
| Cyp26b1_B1 | gAggAgggCAgCAAACggAACATTACACTCCAGCCTTTTGGGATT |
| Cyp26b1_B1 | TGTGTCATGTGTATCCCGTATACTATAgAAgAgTCTTCCTTTACg |
| Cyp26b1_B1 | gAggAgggCAgCAAACggAAAACGTCCACATCCTTGAAGACCGGT |
| Cyp26b1_B1 | GTCTTGACCGAAGCGGTCTGGGTCATAgAAgAgTCTTCCTTTACg |
| Cyp26b1_B1 | gAggAgggCAgCAAACggAAGAACCTTCCATCCTTGTCTTCAGTG |
| Cyp26b1_B1 | TATTCCACCGCCAAATGGGAGATAGTAgAAgAgTCTTCCTTTACg |
| Cyp26b1_B1 | gAggAgggCAgCAAACggAATTGAGGAAAAGCTTTGCCAGATTTT |
| Cyp26b1_B1 | GTGCTGGCCAGTTCGATGGCTAGGGTAgAAgAgTCTTCCTTTACg |
| Cyp26b1_B1 | gAggAgggCAgCAAACggAAGTCCGTGTGGCCAGCTCAAATCGGC |
| Cyp26b1_B1 | ACAGGAACCGGCATCACACGTGGAATAgAAgAgTCTTCCTTTACg |
| Cyp26b1_B1 | gAggAgggCAgCAAACggAAGTTCTGGTTGGAATCAAGTCCAAAA |
| Cyp26b1_B1 | CATTGTTTCTGTCTCTGTTATGATTTAgAAgAgTCTTCCTTTACg |
| Cyp26b1_B1 | gAggAgggCAgCAAACggAAGGAAGAATGGAAACATACACTGGGG |
| Cyp26b1_B1 | TTTGCATTATTCCCCTGACAGGGGCTAgAAgAgTCTTCCTTTACg |
| Cyp26b1_B1 | gAggAgggCAgCAAACggAAGTTTCTGTACTTGATTTGTATTGTT |
| Cyp26b1_B1 | TGATGTCGCTGTTTAGCTTCTCTGTTAgAAgAgTCTTCCTTTACg |
| **Shox** | |
| Shox_B1 | gAggAgggCAgCAAACggAAGTCCGAGATGGGCTGCGCCGGGCTT |
| Shox_B1 | GGGAGGGGCTTTGGCGTATCGAAGATAgAAgAgTCTTCCTTTACg |
| Shox_B1 | gAggAgggCAgCAAACggAACTCTCCTCTCCGCTCGTCCTCTGGT |
| Shox_B1 | GGTCCTCGGCTGGTCCAGGCCACCCTAgAAgAgTCTTCCTTTACg |
| Shox_B1 | gAggAgggCAgCAAACggAAAGCCCTGTCCGCGGACGCCCGCTCC |
| Shox_B1 | CCGAGCCCGAGAGAGAGGGAGAGTGTAgAAgAgTCTTCCTTTACg |
| Shox_B1 | gAggAgggCAgCAAACggAAAGACAAACGCCGTGAGCTCCTCCAT |
| Shox_B1 | CCTTGCTCTTCTGGTCGAAAGACTTTAgAAgAgTCTTCCTTTACg |
| Shox_B1 | gAggAgggCAgCAAACggAAGCCTTCCTTGCGGCCGGCTCCGGGG |
| Shox_B1 | GCTCTCCAGCACCTCCCGGTACGTGTAgAAgAgTCTTCCTTTACg |
| Shox_B1 | gAggAgggCAgCAAACggAAAGTGGTGCGCGCCGCCATCGCCGGG |
| Shox_B1 | TCTCCGGCAGCTCCTTGAAGAGGGGTAgAAgAgTCTTCCTTTACg |
| Shox_B1 | gAggAgggCAgCAAACggAACACTCGTAGATGCCTTCCGAACCCC |
| Shox_B1 | GACTTCACGTCCTCCCTCTTCTCCTTAgAAgAgTCTTCCTTTACg |
| Shox_B1 | gAggAgggCAgCAAACggAAAGCTTCGTCTGCCCGTCCTCGTCCT |
| Shox_B1 | AAGTTGGTGCGGCTCCGCCGCTGCTTAgAAgAgTCTTCCTTTACg |
| Shox_B1 | gAggAgggCAgCAAACggAATCCAGCTCGTTGAGCTGCTCCAGGG |
| Shox_B1 | GGGTAGTGGGTCTCGTCGAAGAGCCTAgAAgAgTCTTCCTTTACg |
| Shox_B1 | gAggAgggCAgCAAACggAACTCAGCTCCTCCCGCATGAAGGCGT |
| Shox_B1 | CGCGCTTCGGAGAGGCCCAGCCGCTTAgAAgAgTCTTCCTTTACg |
| Shox_B1 | gAggAgggCAgCAAACggAACTTCTATTCTGGAACCAGACCTGCA |
| Shox_B1 | TGATTTTCCTGCTTCCGACACTTTGTAgAAgAgTCTTCCTTTACg |
| Shox_B1 | gAggAgggCAgCAAACggAAGTGCCGAGGATCACACCTTTATGCA |
| Shox_B1 | ACCCGGCAGGCGTCCAGGTGGCTGGTAgAAgAgTCTTCCTTTACg |
| Shox_B1 | gAggAgggCAgCAAACggAATAAGGCTCCCATGTTCACATAGGGG |
| Shox_B1 | AGCTTGTACCTGTTGGAAAGGCATCTAgAAgAgTCTTCCTTTACg |
| Shox_B1 | gAggAgggCAgCAAACggAAATGGGTCACTCCTTCCAGCTGCAAC |
| Shox_B1 | CAGGTGTGGGTGGAGGTGGTGGTGTTAgAAgAgTCTTCCTTTACg |
| Shox_B1 | gAggAgggCAgCAAACggAAGAACATAAGGTACGGTGCGTGGGCA |
| Shox_B1 | AATAGGAAGTCCAAAGTGAGGAGGTTAgAAgAgTCTTCCTTTACg |
| Shox_B1 | gAggAgggCAgCAAACggAAAGCAGAGGCTGTTTCCGCCAGCGAT |
| Shox_B1 | CTTTGCCGCAGCTGCCACCACAGCATAgAAgAgTCTTCCTTTACg |
| Shox_B1 | gAggAgggCAgCAAACggAACGCAAGTCGGCAATGCTGGAGTTTT |
| Shox_B1 | GCCTCGGCATGTTTCCTGGCTTTGATAgAAgAgTCTTCCTTTACg |
| Shox_B1 | gAggAgggCAgCAAACggAAAAGGACTGGAATTGTCAAAGACCCA |
| Shox_B1 | AGAAGAGTTTCACTTCACCTTCTTATAgAAgAgTCTTCCTTTACg |
| Shox_B1 | gAggAgggCAgCAAACggAACCGGAGAAACTCTTGCAGGCCTTTT |
| Shox_B1 | GGCCAACGCATGTTCTCTGCTGATCTAgAAgAgTCTTCCTTTACg |
| Shox_B1 | gAggAgggCAgCAAACggAATGTGAATGGGAGAGCGCCTTTGACC |
| Shox_B1 | GCAGGCCAATCATCTGCATTTGGATTAgAAgAgTCTTCCTTTACg |
| Shox_B1 | gAggAgggCAgCAAACggAAGGCGGGAAGCACGAGAGGAGGGCGA |
| Shox_B1 | TATGTTATTGATCATTGTCTTCGGATAgAAgAgTCTTCCTTTACg |
| Shox_B1 | gAggAgggCAgCAAACggAAGGCAGAGCGTGGTCCAGAATAGCAT |
| Shox_B1 | GCTCATATCTTAATTATTTTCTGGCTAgAAgAgTCTTCCTTTACg |
| Shox_B1 | gAggAgggCAgCAAACggAATTCTGAACGGTGACAAAATGATGCC |
| Shox_B1 | CGATTGTACCATATTTCTTCCTCCATAgAAgAgTCTTCCTTTACg |
| Shox_B1 | gAggAgggCAgCAAACggAATGTTTATTGTCTGCAAATAAGTGCT |
| Shox_B1 | TTGGCCAAGGTTCAAAGATTGCCAATAgAAgAgTCTTCCTTTACg |
| Shox_B1 | gAggAgggCAgCAAACggAACAGGTGGTGTGATGCTGAAGTGGAC |
| Shox_B1 | GCACAGGGAAGGCAAAACTGACAGCTAgAAgAgTCTTCCTTTACg |
| **Shox2** | |
| Shox2_B2 | CCTCgTAAATCCTCATCAAACTCCTTCTTCTCCTTCACCTTCTGA |
| Shox2_B2 | CTCCAGCACCTCCCGGTACGTGATCAAATCATCCAgTAAACCgCC |
| Shox2_B2 | CCTCgTAAATCCTCATCAAATGCCGGCTCCTCGGGCCCCGTCCTC |
| Shox2_B2 | GGTCCAGCTCCAGCGCCGGGGAGCGAAATCATCCAgTAAACCgCC |
| Shox2_B2 | CCTCgTAAATCCTCATCAAAGGGAGTCCCGGCTCCTTTCGATGGT |
| Shox2_B2 | GCTCCGGGGACACGTCAGTCAGTTTAAATCATCCAgTAAACCgCC |
| Shox2_B2 | CCTCgTAAATCCTCATCAAACCTTCAGGTCCTCCTTCCTCTCCTT |
| Shox2_B2 | TCTTGGTCTGGCCCTCGTCGTCGAGAAATCATCCAgTAAACCgCC |
| Shox2_B2 | CCTCgTAAATCCTCATCAAAAGTTGGTGCGGCTCCTGCGCTGCTT |
| Shox2_B2 | CCAGCTCGTTCAGCTGCTCCAGAGTAAATCATCCAgTAAACCgCC |
| Shox2_B2 | CCTCgTAAATCCTCATCAAAGGTAGTGGGTCTCGTCGAACAGCCG |
| Shox2_B2 | TCAGCTCCTCCCGCATGAAGGCGTCAAATCATCCAgTAAACCgCC |
| Shox2_B2 | CCTCgTAAATCCTCATCAAAGGGCCTCGGAGAGCCCCAGCCGCTG |
| Shox2_B2 | TTCGGTTCTGGAACCAAACCTGCACAAATCATCCAgTAAACCgCC |
| Shox2_B2 | CCTCgTAAATCCTCATCAAAGGTTCTCCTGTTTCCGGCACTTCGC |
| Shox2_B2 | CACCGATCAGGACCCCTTTGTGCAGAAATCATCCAgTAAACCgCC |
| Shox2_B2 | CCTCgTAAATCCTCATCAAACTCGGCAGGCTTCAAACTGGCTGGC |
| Shox2_B2 | GGGCGCCCACGTTGACATAGGGTGCAAATCATCCAgTAAACCgCC |
| Shox2_B2 | CCTCgTAAATCCTCATCAAAGACTATCCTGCTGAAACGGCATTCT |
| Shox2_B2 | GAAAGGAGAAGGGCGGCACGTTGCAAAATCATCCAgTAAACCgCC |
| Shox2_B2 | CCTCgTAAATCCTCATCAAATGTCCAGCTGCAGCTGTGCCTGAAC |
| Shox2_B2 | GGTGGTGGTGCGCGTGCGCCACTGCAAATCATCCAgTAAACCgCC |
| Shox2_B2 | CCTCgTAAATCCTCATCAAAGTGCGTGCGCGGCCAGGTGCGGGTG |
| Shox2_B2 | AGGGCGGCCCGGGGAACATCATGTAAAATCATCCAgTAAACCgCC |
| Shox2_B2 | CCTCgTAAATCCTCATCAAACCGCCAGCGTGGCCAATGGGAGTCC |
| Shox2_B2 | CGACTGACGCAGCCGACGCCGTCTCAAATCATCCAgTAAACCgCC |
| Shox2_B2 | CCTCgTAAATCCTCATCAAATCGTGGTCTTGGCCGCCGCCGCCGC |
| Shox2_B2 | GCAGGTCGGCGATGCTGGAGTTCTTAAATCATCCAgTAAACCgCC |
| Shox2_B2 | CCTCgTAAATCCTCATCAAACGGCCGCGTGCTTCTTGGCCTTGAG |
| Shox2_B2 | CTTCCGCTGCCAGTCACAGCCCCAGAAATCATCCAgTAAACCgCC |
| Shox2_B2 | CCTCgTAAATCCTCATCAAAGGAAGTACGTAGGAGCGGCGCGGGG |
| Shox2_B2 | CATTTCCGGCTCACAGATACGGGCAAAATCATCCAgTAAACCgCC |
| Shox2_B2 | CCTCgTAAATCCTCATCAAAAACCCATTCCTTCTCCGAGTGGCAG |
| Shox2_B2 | GGTCTTTCCAGGCCGATCGTTGATCAAATCATCCAgTAAACCgCC |
| Shox2_B2 | CCTCgTAAATCCTCATCAAAGCTTGGGCCTGTTTTGCCACCTTTC |
| Shox2_B2 | ACCTAGCTGTGCGCAGAGCGCCTCCAAATCATCCAgTAAACCgCC |
| Shox2_B2 | CCTCgTAAATCCTCATCAAAAGAGTGGTGTTTATTCGTCCTCAGG |
| Shox2_B2 | GGTTCCCAGATTTGGAAGCAGGCCTAAATCATCCAgTAAACCgCC |
| Shox2_B2 | CCTCgTAAATCCTCATCAAAAGGGCTGTGAAGAGACAACATGGCC |
| Shox2_B2 | ATGAACAGTGGTCCATGGATTGGATAAATCATCCAgTAAACCgCC |
| Shox2_B2 | CCTCgTAAATCCTCATCAAAGATTTCACCACTCGCTGCATTTTGT |
| Shox2_B2 | TGCATCAGAGGAATTTGCCTAGATGAAATCATCCAgTAAACCgCC |
| Shox2_B2 | CCTCgTAAATCCTCATCAAATCACTGCAGATTCCTCAGAATAAGA |
| Shox2_B2 | AGCGGTTGAAATGTGCTGACAAGGTAAATCATCCAgTAAACCgCC |
| Shox2_B2 | CCTCgTAAATCCTCATCAAACGCACCTCAGGTCATTGACCAGAGG |
| Shox2_B2 | GATGCACGTGCAGGAAGAGCAGTGAAAATCATCCAgTAAACCgCC |
| Shox2_B2 | CCTCgTAAATCCTCATCAAATGGGCGTCAATGCCACAGTTCACCC |
| Shox2_B2 | ATGGGGTCGTGCTGGCTGTCCTTGCAAATCATCCAgTAAACCgCC |
| Shox2_B2 | CCTCgTAAATCCTCATCAAATATTGTTGCATATTCCTAGCATGGC |
| Shox2_B2 | CAGACAAACGAACTGTCGTCAGTGGAAATCATCCAgTAAACCgCC |
| Shox2_B2 | CCTCgTAAATCCTCATCAAACGTGCTCGCCCATGAGGGCTATGAT |
| Shox2_B2 | AAACATTGAAGAATATATTGTTTGAAAATCATCCAgTAAACCgCC |
| Shox2_B2 | CCTCgTAAATCCTCATCAAATTTCTATGGAAGTGCCTGTGGCCGG |
| Shox2_B2 | TCCCATTCATGGACCTTATCCTGCAAAATCATCCAgTAAACCgCC |
| Shox2_B2 | CCTCgTAAATCCTCATCAAATTGATAGACGAGCATCGGCACCTGG |
| Shox2_B2 | TTCCTGGGCTTACCAAGGACACACGAAATCATCCAgTAAACCgCC |
| Shox2_B2 | CCTCgTAAATCCTCATCAAAGCTGTGCGCGTCAACGCCGAGCATT |
| Shox2_B2 | TGTTGGATGGATTTTGGCCTATGGCAAATCATCCAgTAAACCgCC |
| Shox2_B2 | CCTCgTAAATCCTCATCAAACCCAGGGGTTGCTATGTACTCCTGT |
| Shox2_B2 | GATGCTGCTCGCATTAGATGCCATAAAATCATCCAgTAAACCgCC |
| Shox2_B2 | CCTCgTAAATCCTCATCAAACTGAATGTTGACGCACGGGTTGCCA |
| Shox2_B2 | GTGTATTCTACATTGGCCAGTGTTTAAATCATCCAgTAAACCgCC |
| Shox2_B2 | CCTCgTAAATCCTCATCAAAGTGTAGAAATATAATCTTCTCTTTT |
| Shox2_B2 | ATTTGCCCATCGTCTGTCGAATATCAAATCATCCAgTAAACCgCC |
| Shox2_B2 | CCTCgTAAATCCTCATCAAATTTTCTCTCCCTCAAGTGACATTTT |
| Shox2_B2 | ATTCTTAAATAATGTTATTCGCATCAAATCATCCAgTAAACCgCC |
| Shox2_B2 | CCTCgTAAATCCTCATCAAATTGCCGGCTGCATTTCTCCTTAATC |
| Shox2_B2 | AAAGGGTCAATTGCAAAACCTGTGAAAATCATCCAgTAAACCgCC |
| Shox2_B2 | CCTCgTAAATCCTCATCAAACTTGTCATAGTAAAGTGAACTTGCA |
| Shox2_B2 | ACAGGAGGGCCCGCCTCGCTTATCAAAATCATCCAgTAAACCgCC |
| Shox2_B2 | CCTCgTAAATCCTCATCAAAGGAAACACGAGGATCTGGTTCATGG |
| Shox2_B2 | CCCCAGCCGCTCTTCCAGGAGTGCTAAATCATCCAgTAAACCgCC |
| Shox2_B2 | CCTCgTAAATCCTCATCAAATTCCGTTCTGTGCACGGAGTCTTTT |
| Shox2_B2 | CAAGTAAGTACATGCAAAGCAGAACAAATCATCCAgTAAACCgCC |
| **Sox9** | |
| Sox9_B3 | TCTCCTGCTCCTCGGTCATCTTCATTTCCACTCAACTTTAACCCg |
| Sox9_B3 | gTCCCTgCCTCTATATCTTTAGGGGCTGGGCGCTCCGGACAGGCA |
| Sox9_B3 | GCGAGCCCGCCGAGTCCTCGGACATTTCCACTCAACTTTAACCCg |
| Sox9_B3 | gTCCCTgCCTCTATATCTTTCGTCCGAGCTGGAGCCCGACGGGCA |
| Sox9_B3 | CGTTCTCCAGGGGCCGGGTGTTCTCTTCCACTCAACTTTAACCCg |
| Sox9_B3 | gTCCCTgCCTCTATATCTTTGCTCCTGCAGCTCGCCCTTCGGGAA |
| Sox9_B3 | ACTTGTCCTCCTCGCTCTCCTTCTTTTCCACTCAACTTTAACCCg |
| Sox9_B3 | gTCCCTgCCTCTATATCTTTTGACGGCCTCGCGGATGCAGACGGG |
| Sox9_B3 | TCCAGTCGTAGCCCTTGAGCACCTGTTCCACTCAACTTTAACCCg |
| Sox9_B3 | gTCCCTgCCTCTATATCTTTTCACCCGCACGGGCATGGGCACCAG |
| Sox9_B3 | CGTGCGGCTTGCTCTTGCTGGAGCCTTCCACTCAACTTTAACCCg |
| Sox9_B3 | gTCCCTgCCTCTATATCTTTCCATGAAGGCGTTCATGGGCCGCTT |
| Sox9_B3 | GCTTCCTGCGCGCCGCCTGCGCCCATTCCACTCAACTTTAACCCg |
| Sox9_B3 | gTCCCTgCCTCTATATCTTTTGTGCAGGTGCGGGTACTGGTCGGC |
| Sox9_B3 | TGCCCAGGGTCTTGCTGAGCTCGGCTTCCACTCAACTTTAACCCg |
| Sox9_B3 | gTCCCTgCCTCTATATCTTTCGCCCTCGTTCAGCAGCCTCCAGAG |
| Sox9_B3 | CGGCCTCCTCCACGAAGGGACGTTTTTCCACTCAACTTTAACCCg |
| Sox9_B3 | gTCCCTgCCTCTATATCTTTCTTTCTTGTGCTGCACCCGCAGCCT |
| Sox9_B3 | TGGGCTGGTACTTGTAGTCGGGGTGTTCCACTCAACTTTAACCCg |
| Sox9_B3 | gTCCCTgCCTCTATATCTTTGGCCGTTCTTCACCGACTTCCTCCG |
| Sox9_B3 | GTTCGGCGCCCTCCTCCTGGTCGGCTTCCACTCAACTTTAACCCg |
| Sox9_B3 | gTCCCTgCCTCTATATCTTTAGATGGCGTTGGGCGAGATGTGCGT |
| Sox9_B3 | GCGGCGAGTCGGCCTGTAGCGCCTTTTCCACTCAACTTTAACCCg |
| Sox9_B3 | gTCCCTgCCTCTATATCTTTGCACCTCGCTCATGCCGGAGGAGGA |
| Sox9_B3 | ACTGACCGGAGTGTTCGCCAGGGGATTCCACTCAACTTTAACCCg |
| Sox9_B3 | gTCCCTgCCTCTATATCTTTTGGTGGGAGGGGTTGGTGGCCCCTG |
| Sox9_B3 | TGCCAGGTTGCACGTCGGTTTTGGGTTCCACTCAACTTTAACCCg |
| Sox9_B3 | gTCCCTgCCTCTATATCTTTGGCGCCCTTCTCTCTTAAGGTCTGG |
| Sox9_B3 | GAGGCTGCCTGCCTTCCTCCTGGAGTTCCACTCAACTTTAACCCg |
| Sox9_B3 | gTCCCTgCCTCTATATCTTTTGTCCACGTCCCCAAAGTCAATATG |
| Sox9_B3 | AGATGACGTCGCTACTTAGCACTGATTCCACTCAACTTTAACCCg |
| Sox9_B3 | gTCCCTgCCTCTATATCTTTGTCGAACTCCTGGATGTCGATGGGG |
| Sox9_B3 | AGGGTGACTATTGGGAGGAAGGTACTTCCACTCAACTTTAACCCg |
| Sox9_B3 | gTCCCTgCCTCTATATCTTTGGCCTGTCCATGGGTGACGGGCACC |
| Sox9_B3 | GTAGCTGCTGGTGTAGGTGCCGGGGTTCCACTCAACTTTAACCCg |
| Sox9_B3 | gTCCCTgCCTCTATATCTTTGCAAGCCAGGCATGGGCGGGACCCC |
| Sox9_B3 | AGGGTGGTCAGCGTGTGCTGCTGCTTTCCACTCAACTTTAACCCg |
| Sox9_B3 | gTCCCTgCCTCTATATCTTTTGCTGGCCCTGCCCCTGCTCGTTGC |
| Sox9_B3 | AGCTGCTCCGTCTTGATGTGTGTCCTTCCACTCAACTTTAACCCg |
| Sox9_B3 | gTCCCTgCCTCTATATCTTTTGCTGCTCGCTGTAGTGTGTGGGGC |
| Sox9_B3 | TAGCTCAGCTGCTGCGGGGAATGTTTTCCACTCAACTTTAACCCg |
| Sox9_B3 | gTCCCTgCCTCTATATCTTTTAGTGCTGCTGCAGGTTGAAAGGGC |
| Sox9_B3 | CGCGTGATGGTGGGGTAGGTGGCGTTTCCACTCAACTTTAACCCg |
| Sox9_B3 | gTCCCTgCCTCTATATCTTTTGGTGGTCGGTGTAGTCGTACTGGG |
| Sox9_B3 | GCGTGGCTGTAGTAGCTGTTAGAGCTTCCACTCAACTTTAACCCg |
| Sox9_B3 | gTCCCTgCCTCTATATCTTTGAGTACAGGCTTGAGCTCTGACCCG |
| Sox9_B3 | TGTGTGGGGTTCATGTAGGAGAAGGTTCCACTCAACTTTAACCCg |
| Sox9_B3 | gTCCCTgCCTCTATATCTTTTCTGCGATAGGGGTGTACATGGGGC |
| Sox9_B3 | TGCGGGATGGAGGGGACGCCCGTCGTTCCACTCAACTTTAACCCg |
| Sox9_B3 | gTCCCTgCCTCTATATCTTTAGACTGGCTGCTCCCAGTGCTGGGG |
| Sox9_B3 | GCCTCTAGGGCCGGGTGAGCTGTGTTTCCACTCAACTTTAACCCg |
| Sox9_B3 | gTCCCTgCCTCTATATCTTTTCTTCAAAGTCTGCAGGGCAGCTTG |
| Sox9_B3 | CGGTAACCAAGATGGCCGACTTAAGTTCCACTCAACTTTAACCCg |
| Sox9_B3 | gTCCCTgCCTCTATATCTTTACAAGCTCTTGGTCCTTCCAACATG |
| Sox9_B3 | GTTCGAGCCCAAGCTGTAGTGCAGATTCCACTCAACTTTAACCCg |
| Sox9_B3 | gTCCCTgCCTCTATATCTTTGGGTTCACTGTCCATCCGTTTCCCC |
| Sox9_B3 | AATGTACAGGGTTTCTGGGCCCCGATTCCACTCAACTTTAACCCg |
| Sox9_B3 | gTCCCTgCCTCTATATCTTTAGTGTTCGGTTCTGAGATGTCCTCT |
| Sox9_B3 | TTTCCAGTCTTTGTGAGAACAGAGGTTCCACTCAACTTTAACCCg |
| Sox9_B3 | gTCCCTgCCTCTATATCTTTCTGTTCCTTAATTACCGTTATAAAA |
| Sox9_B3 | TATTCCTCACAGAGGATTTTCTATTTTCCACTCAACTTTAACCCg |
| Sox9_B3 | gTCCCTgCCTCTATATCTTTTTCCACCCATAGGTTTAAATTGGAA |
| Sox9_B3 | GCACCAGTCGTGCCTTTGTCTGCAGTTCCACTCAACTTTAACCCg |
| Sox9_B3 | gTCCCTgCCTCTATATCTTTGTGTCTGCTGCGTTTCGTGTTCCTC |
| Sox9_B3 | GCATCCCGATTGGCTGGCTTCGAGGTTCCACTCAACTTTAACCCg |
| Sox9_B3 | gTCCCTgCCTCTATATCTTTCCCCTCAGGTGCAAGCGTGCACCCC |
| Sox9_B3 | AGAATTGTGCAGGCAAAGGCGGCGTTTCCACTCAACTTTAACCCg |
| Sox9_B3 | gTCCCTgCCTCTATATCTTTACACAGTACATACTACAACGTGTGG |
| Sox9_B3 | TCTCCACAATAAAAGTCGATGAAACTTCCACTCAACTTTAACCCg |
| Sox9_B3 | gTCCCTgCCTCTATATCTTTGGAGAAACAGTTTTGCTTTCTGGGG |
| Sox9_B3 | TGCAAGAAAACTATGGGAAACAGTGTTCCACTCAACTTTAACCCg |
| Sox9_B3 | gTCCCTgCCTCTATATCTTTAATGAAGCAAAGCTGAACTGGAAAA |
| Sox9_B3 | TGTATCACTCCACGTCACAAGTTACTTCCACTCAACTTTAACCCg |
| Sox9_B3 | gTCCCTgCCTCTATATCTTTGCACAAACAAACGGCCCACAATTTT |
