## Supplementary material for "Retinoic acid breakdown is required for proximodistal positional identity during amphibian limb regeneration": Table S1

| Primer name | 5' --> 3' sequence |
| --- | --- |
| Meis1_qPCR_F | TCCCCCTGAACGTGTTGATG |
| Meis1_qPCR_R | AGTCGGACAGCGAAGACATG |
| Meis2_qPCR_F | TGTCATGCCCAGCAGTATGG |
| Meis2_qPCR_R | ACTTCTCAAAGACCAGGGCG |
| Hoxa9_qPCR_F | GGAGACAAGCCTGCCATTGA |
| Hoxa9_qPCR_R | TGTAAGGGCAGCGCTTCTTG |
| Hoxa11_qPCR_F | GAGTCCTCTTCCGGCAACAA |
| Hoxa11_qPCR_R | TTCGCGGATCTGGTACTTGG |
| Hoxa13_qPCR_F | ACTGTTCCAAGGAGCAAGGG |
| Hoxa13_qPCR_R | GTACGAGTTCGCATCCGAGG |
| Rara_qPCR_F | CATTATGGTGTCAGCGCGTG |
| Rara_qPCR_R | GATGATGCACGTCTTGTCGC |
| Rarg_qPCR_F | CAAAGGTCAGCAAGGCACAC |
| Rarg_qPCR_R | TCGCCAGCTCGCTGAATTTA |
| Raldh1_qPCR_F | ATCGACAAGGCACTGACGTT |
| Raldh1_qPCR_R | TTTCTCTCCCGTTGCCTGAC |
| Raldh2_qPCR_F | ATGTCCGGTAATGGGAGGGA |
| Raldh2_qPCR_R | CTGCAGCGCTCTTAGGAGTT |
| Raldh3_qPCR_F | GAACATAGTCCCCGGCTACG |
| Raldh3_qPCR_R | TGGAGGCAGCTTCCTTGATG |
| Cyp26a1_qPCR_F | CATCGATGTGCCTTTCTGCG |
| Cyp26a1_qPCR_R | GAGTCCTTGGCCATTTTCGC |
| Cyp26b1_qPCR_F | AAAGAGCATGGCAAGGAGCT |
| Cyp26b1_qPCR_R | TTGTGGAGGATACCGTTGCC |
| GFP_qPCR_F | AGCTGAAGGGCATCGACTTC |
| GFP_qPCR_R | TTCTGCTTGTCGGCCATGAT |
| Shox_qPCR_F | TCAGTTGCAGCTGGAAGGAG |
| Shox_qPCR_R | GTCCAAAGTGAGGAGGTGGG |
| Shox2_qPCR_F | GAAGTGCCGGAAACAGGAGA |
| Shox2_qPCR_R | CGTTGACATAGGGTGCGACT |
| Ef1a_qPCR_F | AACATCGTGGTCATCGGCCAT |
| Ef1a_qPCR_R | GGAGGTGCCAGTGATCATGTT |
